# Resolving challenges in quantitative modeling of microbial community dynamics

**DOI:** 10.1101/356519

**Authors:** Samuel F. M. Hart, Hanbing Mi, Robin Green, Li Xie, Jose Mario Bello Pineda, Babak Momeni, Wenying Shou

## Abstract

Microbial communities can perform biochemical activities that monocultures cannot. Controlling communities requires an understanding of community dynamics. Here, we mathematically predict the growth rate of an engineered community consisting of two *S. cerevisiae* strains, each releasing a metabolite required and consumed by the partner. Initial model parameters were based on strain phenotypes measured in batch mono-cultures with zero or excess metabolite, and failed to quantitatively predict experimental results. To resolve model-experiment discrepancy, we chemically identified the correct exchanged metabolites, but this did not improve model performance. We then re-measured strain phenotypes in chemostats mimicking the metabolite-limited community environments, while mitigating or incorporating effects of rapid evolution. Almost all phenotypes we measured varied significantly with the metabolite environment. Once we used parameters measured in community-like chemostat environments, prediction agreed with experimental results. In summary, using a simplified community, we uncovered, and devised means to resolve, modeling challenges that are likely general.

## Introduction

Multi-species microbial communities are ubiquitous. In a community, member species *interact* in that one species alters the physiology of another species. Consequently, communities often display community-level properties not achievable by member species in isolation. For example, a community assembled from six intestinal bacterial species, but not any of the individual species alone, clears *Clostridium difficile* infection in mice ^1^. As another example, a two-species community is required for efficient industrial production of Vitamin C ^2,3^. Thus, even simplified communities of a small number of species can be useful for biotechnology applications ^4–8^.

An important community-level property is community dynamics, including how species abundance changes over time ^9^. Community dynamics can be predicted using statistical correlation models. For example, community dynamics observed over a period of time can be used to construct a model which correlates the abundance of one species with the growth rate of another, and the model can then be used to predict future dynamics ^10–12^. However even for two-species communities, statistical correlation models might generate false predictions on species coexistence ^13^.

Alternatively, mathematical models based on species interaction mechanisms should avoid the pitfalls of statistical correlations, but species interactions are generally difficult to identify ^9^. Genome-scale metabolic models use genome sequences, sometimes in conjunction with RNA and protein expression profiles, to predict metabolic fluxes within species as well as metabolic fluxes among-species (i.e. metabolic interactions) ^14,15^. However, these models face multiple challenges including unknown protein functions or metabolic fluxes ^16^. Even when interaction mechanisms are known ^14,17–20^, often a fraction of model parameters are “free” (unmeasured) due to difficulties in measuring parameters. The values assigned to free parameters are generally “guesstimates” or literature values “borrowed” from a different strain or even a different species, and can vary by orders of magnitude ^21^. Using free parameters may work for qualitative modeling, but often not for quantitative modeling. If there are no free parameters, then model-experiment disagreements would suggest that important pieces are missing in either the model or the experiments.

Previously, we have constructed and mathematically modeled an engineered yeast community CoSMO (<u>Co</u>operation that is <u>S</u>ynthetic and <u>M</u>utually <u>O</u>bligatory) ^17^. CoSMO consists of two differentially-fluorescent, non-mating haploid *S. cerevisiae* strains (Fig 1A; Fig 1-Table Supplement 1). One strain, designated *A*^*−*^*L*^*+*^, cannot synthesize adenine because of a deletion mutation in the *ADE8* gene, and over-activates the lysine biosynthetic pathway due to a feedback-resistant *LYS21* mutation ^22^. The other strain, designated *L*^*−*^*A*^*+*^, requires lysine because of a deletion mutation in the *LYS2* gene, and over-activates the adenine biosynthetic pathway due to a feedback-resistant *ADE4* mutation ^23^. Overproduced metabolites in both strains are released into the environment, which are consumed by the partner. In minimal medium lacking adenine and lysine supplements, the two strains engage in obligatory cooperation. Released metabolites are rapidly consumed and thus are present at very low concentrations in CoSMO.

**Fig. 1.**
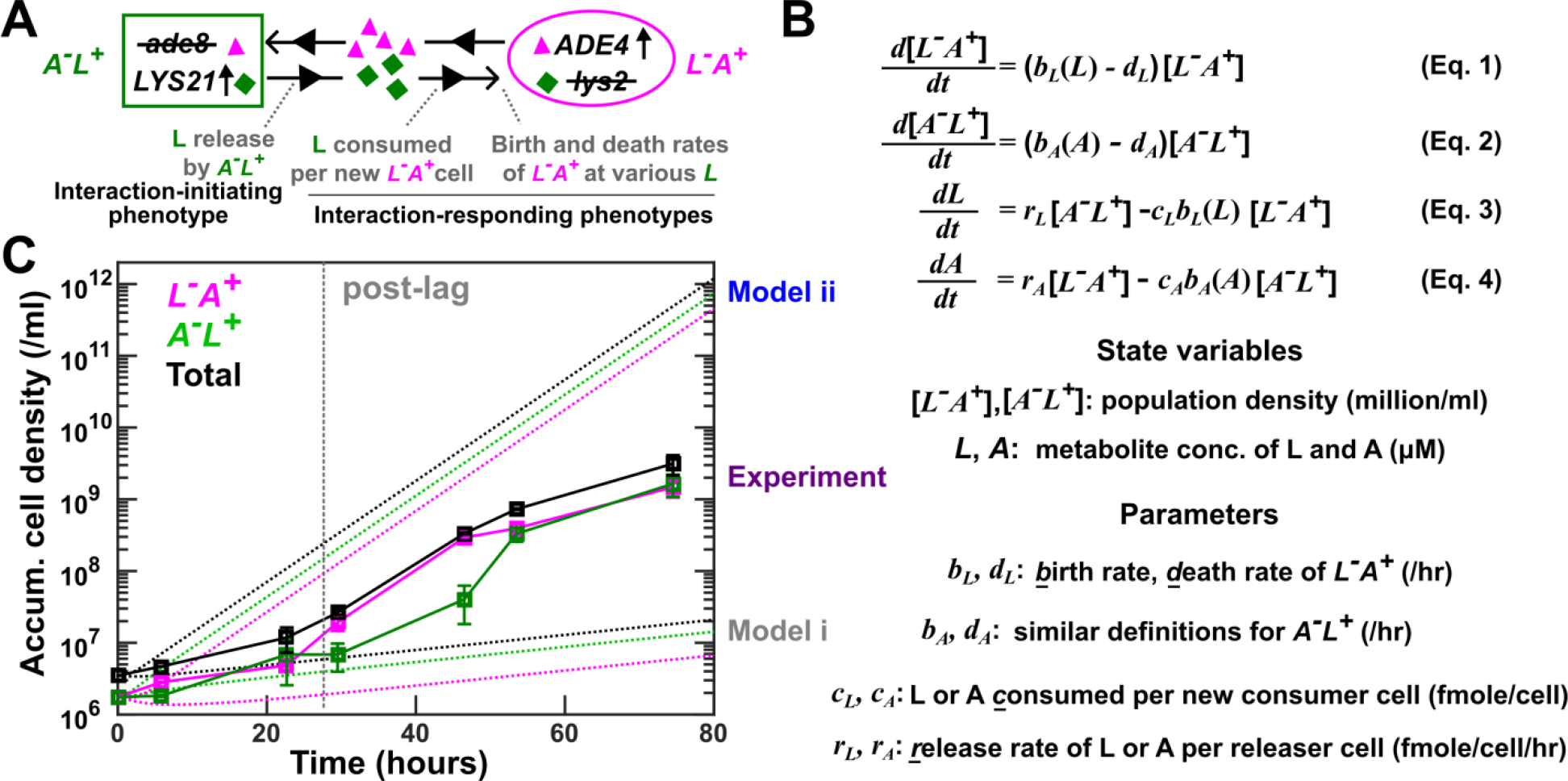
Model-experiment discrepancy in CoSMO growth dynamics. (**A**) CoSMO comprises two non-mating, cross-feeding yeast strains expressing different fluorescent proteins. Here, we use the lysine-mediated interaction as an example to illustrate interaction-initiating and interaction-responding phenotypes. (**B**) A mathematical model of well-mixed CoSMO. For example, [*L*^*−*^*A*^*+*^], the density of *L*^*−*^*A*^*+*^, increases due to cell birth (which is a function of lysine concentration *L*) and decreases due to cell death. Lysine concentration *L* increases due to release by *A*^*−*^*L*^*+*^, and decreases due to consumption coupled to *L*^*−*^*A*^*+*^ birth. (**C**) Model-experiment discrepancy. In experiments (squares; data listed in Fig 1-Table Supplement 2), *L*^*−*^*A*^*+*^ and *A*^*−*^*L*^*+*^ growing exponentially in excess lysine or adenine were washed and preconditioned (“Strain culturing and preconditioning” in Methods). They were mixed at 1:1 in SD at time zero to form CoSMO, which was then cultured in a well-mixed environment and diluted as needed to ensure that no other nutrients were limiting. Population dynamics were tracked using flow cytometry (“Flow cytometry” in Methods), and dilutions were taken into account when calculating accumulative cell densities. CoSMO initially grew slowly (prior to grey dotted line), and then grew faster. We simulated CoSMO population dynamics (dotted lines; Fig 1-Code Supplement 1) using Eq. 1-4 (**B**) where parameters were measured in batch cultures (Fig 1-Table Supplement 2). Models i and ii differed in that the release rates of the two strains were “borrowed” from our previous measurements in the S288C background (Model i) or directly measured in the RM11 background (Model ii).

We have formulated a differential equation model of the CoSMO dynamics (e.g. Eq. 1-4 in Fig 1B). Population growth is dictated by cell birth (which in turn depends on the concentration of the required metabolite) as well as cell death, while metabolite concentration is dictated by release and consumption. Model parameters correspond to strain phenotypes including metabolite release rate, metabolite consumption per birth, and cell birth and death rates (Fig 1B). Even though these phenotypes reflect strain interactions (“interaction phenotypes” in Fig 1A), we measured them in mono-cultures to eliminate partner feedback. In our earlier studies, we quantified some of these phenotypes and borrowed other from literature values ^17,24,25^. Our models correctly predicted various properties of CoSMO, including the steady state strain ratio ^17^ as well as qualitative features of spatial patterning ^24,25^.

In the current study, we aim to predict CoSMO growth rate (the rate of total population increase), a measure of how likely the community can survive periodic dilutions such as those in industrial fermenters ^26^. Our initial model predictions of CoSMO growth rate significantly deviated from experimental measurements. In the process of resolving this model-experiment discrepancy, we have uncovered and overcome multiple challenges. Since these challenges are likely general, our work serves as a “roadmap” that can be applied to quantitative modeling of other simplified microbial communities.

## Results

Experimentally, CoSMO growth followed a reproducible pattern: after an initial lag marked by slow growth, the two populations and thus the entire community grew at a faster rate (Fig 1C, “Experiment”). The latter growth rate eventually reached a steady state (Fig 7-Figure Supplement 4A, bottom panels). We wanted to quantitatively predict CoSMO’s post-lag steady state growth rate (“growth rate”), with the criterion that model prediction should fall within experimental error bars. To predict community growth rate, we either quantified it from simulated community dynamics (Fig 1C, dotted lines), or calculated it from an analytical formula (Eq. 15 or 16 in Methods “Calculating steady state community growth rate”), with similar results (e.g. Fig 1-Figure Supplement 1). The analytical formula suggests that community growth rate is affected by metabolite release rate, metabolite consumption per birth, and death rate.

### Initial models based on batch-culture parameters poorly predict community growth rate

Our first model (Model i) under-estimated community growth rate. Unlike the published strains of *A*^*−*^*L*^*+*^ and *L*^*−*^*A*^*+*^ in the S288C background ^17^, strains in this study were constructed in the RM11 background to reduce mitochondrial mutation rate ^27^. Thus, we had to consider re-quantifying strain phenotypes. For each RM11 strain, we measured death rate during starvation using a microscopy batch culture assay ^28^. We also quantified the amount of metabolite consumed per birth in batch cultures grown to saturation (see Fig 4B for details), similar to our earlier work ^17^. Since release rates were tedious to measure, we “borrowed” published release rates of *L*^*−*^*A*^*+*^ and *A*^*−*^*L*^*+*^ in the S288C background in batch starved cultures ^17^. Predicted community growth rates were much slower than experimental measurements (Fig 1C “Model i”; Fig 7 grey).

A revised model (Model ii) without any borrowed parameters over-estimated community growth rate. For this model, we directly measured the release rates of RM11 *L*^*−*^*A*^*+*^ and *A*^*−*^*L*^*+*^ in batch starved cultures (see Fig 5A and B for details). The release rates of both strains in the RM11 background were ~3-fold higher than those in the S288C background (Fig 1-Table Supplement 2). Concomitantly, the predicted community growth rate greatly exceeded experiments (Fig 1 C, “Model ii”; Fig 7 blue).

### Identifying interaction mediators

One possible cause for the model-experiment discrepancy could be that cells engineered to overproduce adenine or lysine ^22,23^ might instead release derivatives of adenine or lysine. Consequently, when we quantified phenotypes such as metabolite consumption, we could have supplemented the wrong metabolite and been misled. A genome-scale metabolic model of *S. cerevisiae* predicted that although *A*^*−*^*L*^*+*^ likely released lysine, *L*^*−*^*A*^*+*^ likely released hypoxanthine or adenosine-(3,5)-biphosphate instead of adenine ^29,30^. Nanospray desorption electrospray ionization mass spectrometry imaging (nanoDESI MS) ^31^ performed by Julia Laskin lab revealed a lysine gradient emanating from *A*^*−*^*L*^*+*^ and hypoxanthine and inosine gradients emanating from *L*^*−*^*A*^*+*^, although the signals were noisy (data not available, but see below).

Indeed, lysine mediates the interaction from *A*^*−*^*L*^*+*^ to *L*^*−*^*A*^*+*^. We subjected *A*^*−*^*L*^*+*^ supernatant to HPLC (high pressure liquid chromatography; Methods, “HPLC”) and yield bioassay (Methods, “Bioassays”). In HPLC, a compound in *A*^*−*^*L*^*+*^ supernatant eluted at the same time as the lysine standards (Fig 2A), and its concentration could be quantified by comparing the peak area against those of lysine standards (Fig 2A inset). In bioassay, we quantified the total lysine-equivalent compounds in an *A*^*−*^*L*^*+*^ supernatant by growing *L*^*−*^*A*^*+*^ in it and comparing the final turbidity with those of minimal medium supplemented with lysine standards. HPLC quantification agreed with the yield bioassay (Fig 2B). Thus, lysine-equivalent compounds released by *A*^*−*^*L*^*+*^ were primarily lysine.

**Fig. 2.**
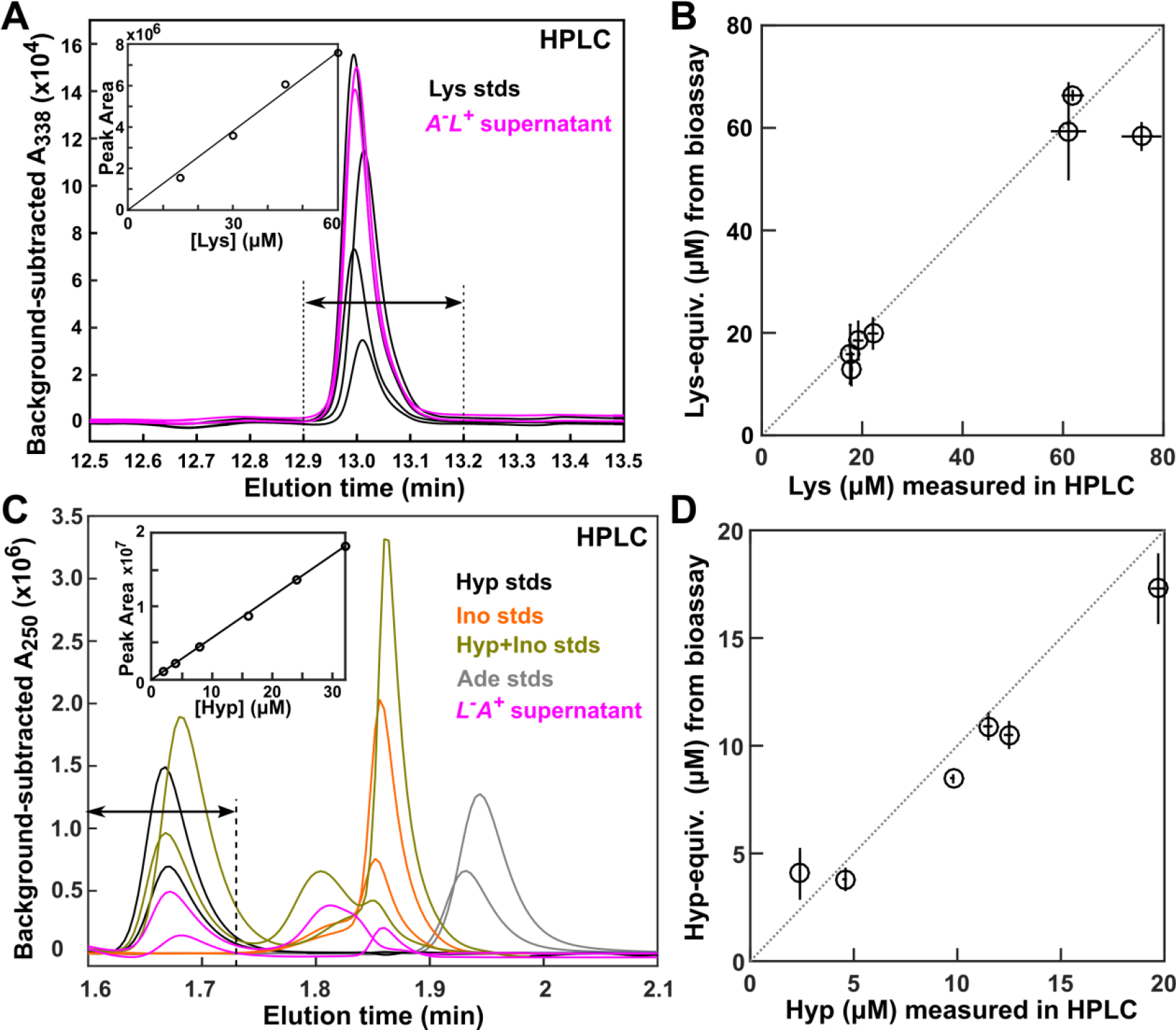
Lysine and hypoxanthine mediate interactions in CoSMO. (**A, B**) Lysine mediates the interaction from *A*^*−*^*L*^*+*^ to *L*^*−*^*A*^*+*^. (**A**) *A*^*−*^*L*^*+*^ releases lysine. Supernatants of *A*^*−*^*L*^*+*^ (magenta), as well as SD supplemented with various concentrations of lysine standards (black) were derivatized and run on HPLC (Methods, “HPLC”). Inset: standard curve where the peak areas between 12.9 min and 13.2 min (arrows) were plotted against lysine concentrations. (**B**) In *A*^*−*^*L*^*+*^ supernatants, lysine concentrations quantified by HPLC agreed with concentrations of lysine-equivalents that supported *L*^*−*^*A*^*+*^ in a yield bioassay (Methods, “Bioassays”). Circles indicate average values, and bars represent the spread of two measurements. (**C, D**) Hypoxanthine mediates the interaction from *L*^*−*^*A*^*+*^ to *A*^*−*^*L*^*+*^. (**C**) *L*^*−*^*A*^*+*^ releases hypoxanthine and inosine. HPLC traces of *L*^*−*^*A*^*+*^ supernatants (magenta) and standards of SD supplemented with two different concentrations of hypoxanthine (black), inosine (orange), adenine (grey), or a mixture of hypoxanthine and inosine (olive). Inset: standard curve where the peak areas in the window between 1.6 min and 1.732 min (arrows) were plotted against hypoxanthine concentrations and used to quantify hypoxanthine. Since the HPLC elution profile could vary between independent runs (e.g. compare the two olive curves), quantification windows were adjusted accordingly. (**D**) In *L*^*−*^*A*^*+*^ supernatants, hypoxanthine concentrations quantified by HPLC agreed with concentrations of purines that supported *A*^*−*^*L*^*+*^ growth as quantified by the yield bioassay. In **B** and **D**, dotted lines have a slope of 1.

Hypoxanthine mediates the interaction from *L*^*−*^*A*^*+*^ to *A*^*−*^*L*^*+*^. When we subjected *L*^*−*^*A*^*+*^ supernatants to HPLC, we found compounds at the elution times of hypoxanthine and inosine, but not of adenine (Fig 2C). Hypoxanthine but not inosine supported *A*^*−*^*L*^*+*^ growth, and inosine did not affect how hypoxanthine stimulated *A*^*−*^*L*^*+*^ growth (Fig 2-Figure Supplement 1). Hypoxanthine concentration quantified by HPLC agreed with the concentration of purines consumable by *A*^*−*^*L*^*+*^ in the yield bioassay (Fig 2D; Methods “Bioassays”). Thus, *A*^*−*^*L*^*+*^ primarily consumed hypoxanthine released by *L*^*−*^*A*^*+*^.

Using phenotypes of *A*^*−*^*L*^*+*^ measured in hypoxanthine versus adenine happened to not affect model performance. Death and release rates were not affected since they were measured in the absence of purine supplements. A similar amount of hypoxanthine and adenine were consumed to produce a new *A*^*−*^*L*^*+*^ cell (Fig 2-Figure Supplement 1). Although the birth rate of *A*^*−*^*L*^*+*^ was slower in the presence of hypoxanthine compared to adenine, especially at low concentrations (Fig 2-Figure Supplement 2), this difference did not affect community growth rate. Thus, distinguishing whether hypoxanthine or adenine was the interaction mediator did not make a difference in predicting community growth rate (Fig 1-Figure Supplement 1). Here, we continue to use “A” to represent the adenine precursor hypoxanthine.

### Rapid evolution during chemostat measurements of strain phenotypes

Model-experiment discrepancy (Fig 1C) could be caused by phenotypes being quantitatively different when measured in batch cultures containing zero or excess metabolite versus in the metabolite-limited CoSMO-like environments. Thus, we re-measured strain phenotypes in chemostats ^32^ that mimicked CoSMO environments. Specifically in a chemostat, fresh medium containing the required metabolite (lysine or hypoxanthine) was pumped into the culturing vessel at a fixed rate (“dilution rate”), while cell-containing medium exited the culturing vessel at the same rate (Methods, “Chemostat culturing”). After an initial adjustment stage, live population density reached a steady state (Fig 3A) which meant that the population grew at the same rate as the dilution rate (Eq. 5-9 in Methods) ^32^. By setting chemostat dilution rate to various growth rates experienced by CoSMO (i.e. 5.5~8 hr doubling), we mimicked the environments experienced by CoSMO and measured strain phenotypes. However, as we demonstrate below, rapid evolution occurred during measurements and needed to be mitigated or incorporated in experiments and in modeling.

**Fig. 3.**
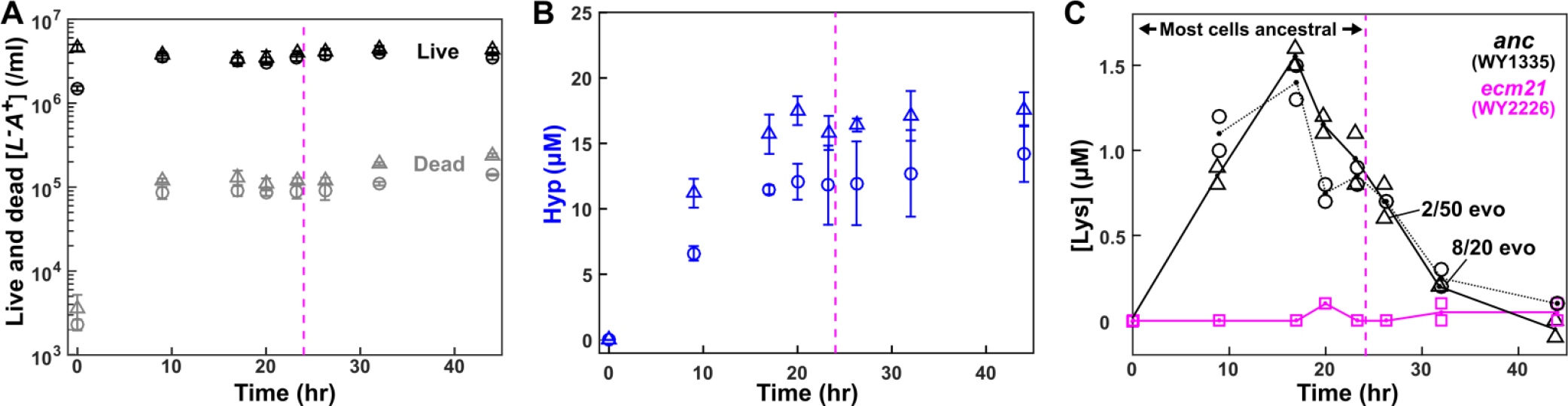
Chemostat dynamics reveals rapid evolution. *L*^*−*^*A*^*+*^ (WY1335) cells growing exponentially in excess lysine were washed free of lysine and inoculated into the culturing vessel (triangles: inoculation at near steady state density in the absence of lysine; circles: inoculation at 1/3 steady state density in the presence of 5~10 lysine). Minimal medium containing 20 lysine was dripped into the culturing vessel (19 ml) to achieve an 8-hr doubling time (19 ml*ln(2)/8 hr = 1.646 ml/hr; “Chemostat culturing” in Methods). (**A**) Live and dead cell densities (Methods, “Flow cytometry”) and (**B**) released hypoxanthine (Methods, “Bioassays”) reached steady state by ~10 hr and ~20 hr, respectively. Error bars represent two standard deviations. (**C**) Lysine concentrations in culturing vessels failed to maintain a steady state due to rapid evolution (Methods, “Detecting evolved clones”). Instead, lysine concentrations rapidly declined to a level similar to that in a chemostat inoculated with an evolved *L*^*−*^*A*^*+*^ mutant with improved affinity for lysine ^64,65^ (“*ecm21*”, magenta). Indeed, when we tested chemostat samples (triangles), 8 out of 20 tested clones were evolved by 32 hours (~4 generations). For each sample, two measurements of lysine concentrations and their average were plotted. Magenta dashed lines mark the time before which >90% of population remained ancestral.

In chemostat measurements, ancestral *L*^*−*^*A*^*+*^ was rapidly overtaken by mutants with dramatically improved affinity for lysine ^33^ (Fig 3C; Fig 3-Figure Supplement 2; Methods, “Detecting evolved clones”). These mutants, likely being present in the inoculum at a low (~10^−6^) frequency, grew 3.6-fold faster than the ancestor (Fig 3-Figure Supplement 2). Thus, to measure ancestral *L*^*−*^*A*^*+*^ phenotypes, we terminated measurements before mutants could take over (<10%; before magenta dashed lines in Fig 3).

In contrast, the evolutionary effects of *A*^*−*^*L*^*+*^ mutants on CoSMO growth were captured during phenotype measurements. Unlike *L*^*−*^*A*^*+*^ mutants, *A*^*−*^*L*^*+*^ mutants were constantly generated from ancestral cells at an extremely high rate (on the order of 0.01/cell/generation; Methods “Evolutionary dynamics of mutant A-L+”), presumably via frequent chromosome duplication (Fig 3-Figure Supplement 3C). Thus, these mutants were already present at a significant frequency (1~10%) even before our measurements started, and slowly rose to 30~40% during measurements due to their moderate fitness advantage over the ancestor under hypoxanthine limitation (Fig 3-Figure Supplement 4; Fig 3-Figure Supplement 3A; Methods, “Detecting evolved clones”). Consequently, we actually measured the average phenotypes of an evolving mixture of ancestors and mutants. Fortunately these averaged phenotypes could be used to model CoSMO since mutants accumulated in similar fashions during phenotype measurements and during CoSMO measurements so long as the two time windows were compatible (Fig 3-Figure Supplement 4B).

### Metabolite consumption is sensitive to the environment

Metabolite consumption per birth depends on the growth environment. Consistent with our previous work ^17^, consumption during exponential growth was higher than that in a culture grown to saturation (Fig 4; Methods, “Measuring consumption in batch cultures”), presumably due to exponential cells storing excess metabolites ^34^. Consumption in chemostats (Methods “Quantifying phenotypes in chemostats”, Eq. 10) was in-between exponential and saturation consumption (Fig 4C for *L*^*−*^*A*^*+*^ and Fig 4-Figure Supplement 1 for *A*^*−*^*L*^*+*^). Since consumption in chemostat was relatively constant across the range of doubling times usually encountered in CoSMO (5.5~8 hr), we used the average value in Model iii (dashed line in Fig 4C and Fig 4-Figure Supplement 1; Table 1).

**Fig. 4.**
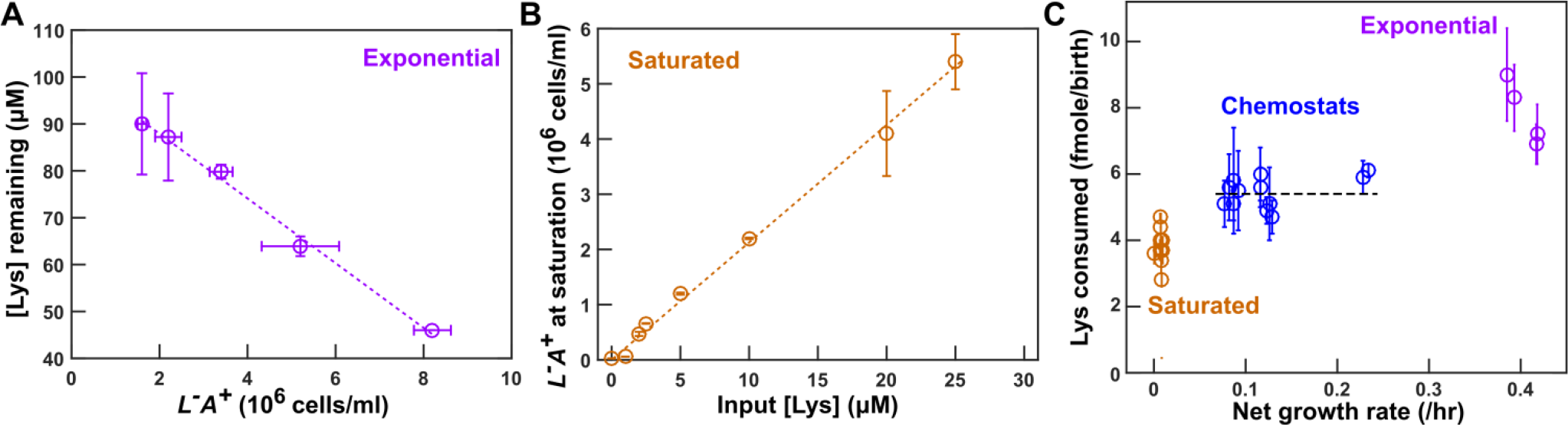
Metabolite consumption is sensitive to the environment. (**A**) Consumption in excess lysine. *L*^*−*^*A*^*+*^ population density and lysine concentration remaining in the medium were measured over time (Methods, “Measuring consumption in batch cultures”). Consumption per birth was calculated from the slope of the lavender line. (**B**) Consumption in cultures grown to saturation. *L*^*−*^*A*^*+*^ cells were inoculated into SD supplemented with various concentrations of lysine. After cultures had reached saturation, total cell densities were measured by flow cytometry. Lysine consumed per birth was quantified from 1/(slope of the orange line). This value was used in Models i and ii. (**C**) Consumption per birth in different environments. Lysine consumption was measured in lysine-limited chemostats at various doubling times (Methods, “Quantifying phenotypes in chemostats” and Eq. 10), and data were jittered slightly along the x-axis to facilitate visualization. For chemostat measurements, error bars represent 2 standard deviations caused by fluctuations in steady state population density. For exponential and saturation consumption, error bars mark 2 standard error of mean for slope estimation. The black dashed line marks the average lysine consumption per *L*^*−*^*A*^*+*^ birth in chemostats (Table 1; Table 1-Table Supplement 1), which we used in Model iii.

**Table 1.**
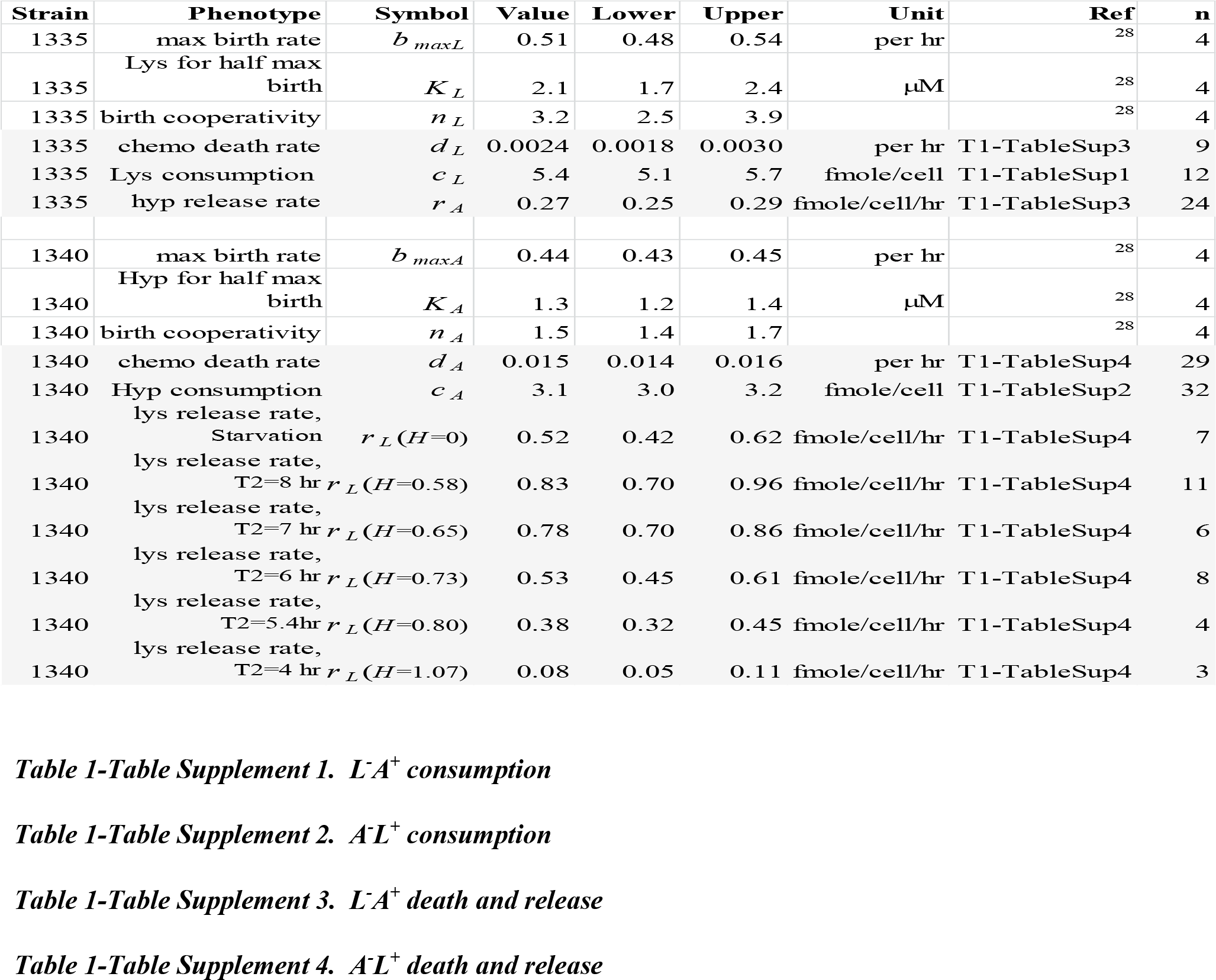
Experimentally-measured strain phenotypes. The upper and lower bounds of consumption amount per birth, release rate, and death rate in chemostats were calculated from the mean value plus and minus two standard errors of mean, respectively. Birth rates *b* at various concentrations *s* of a limiting metabolite were measured using a high-throughput microscopy batch-culture assay, and then fitted into the Moser’s growth model ^66^ *b*(*s*) = *b*_*max*_*s*^*n*^/(*s*^*n*^ + *K*^*n*^), where *b*_*max*_ is the maximal birth rate, *K* is the *s* at which *b*_*max*_/2 is achieved, and *n* is the growth cooperativity (akin to Hill’s coefficient). The averages from four independent experiments were fitted into Moser’s model to infer birth parameters (*b*_*max*_, *K*, and *n*) and their confidence intervals (for details, see ^28^). Birth parameters were used in simulations.

| Strain | Phenotype | Symbol | Value | Lower | Upper | Unit | Ref | n |
| --- | --- | --- | --- | --- | --- | --- | --- | --- |
| 1335 | max birth rate | $b_{maxL}$ | 0.51 | 0.48 | 0.54 | per hr | <sup>28</sup> | 4 |
| 1335 | Lys for half max birth | $K_L$ | 2.1 | 1.7 | 2.4 | $\mu\text{M}$ | <sup>28</sup> | 4 |
| 1335 | birth cooperativity | $n_L$ | 3.2 | 2.5 | 3.9 | | <sup>28</sup> | 4 |
| 1335 | chemo death rate | $d_L$ | 0.0024 | 0.0018 | 0.0030 | per hr | T1-TableSup3 | 9 |
| 1335 | Lys consumption | $c_L$ | 5.4 | 5.1 | 5.7 | fmole/cell | T1-TableSup1 | 12 |
| 1335 | hyp release rate | $r_A$ | 0.27 | 0.25 | 0.29 | fmole/cell/hr | T1-TableSup3 | 24 |
| 1340 | max birth rate | $b_{maxA}$ | 0.44 | 0.43 | 0.45 | per hr | <sup>28</sup> | 4 |
| 1340 | Hyp for half max birth | $K_A$ | 1.3 | 1.2 | 1.4 | $\mu\text{M}$ | <sup>28</sup> | 4 |
| 1340 | birth cooperativity | $n_A$ | 1.5 | 1.4 | 1.7 | | <sup>28</sup> | 4 |
| 1340 | chemo death rate | $d_A$ | 0.015 | 0.014 | 0.016 | per hr | T1-TableSup4 | 29 |
| 1340 | Hyp consumption | $c_A$ | 3.1 | 3.0 | 3.2 | fmole/cell | T1-TableSup2 | 32 |
| 1340 | lys release rate, Starvation | $r_L(H=0)$ | 0.52 | 0.42 | 0.62 | fmole/cell/hr | T1-TableSup4 | 7 |
| 1340 | lys release rate, T2=8 hr | $r_L(H=0.58)$ | 0.83 | 0.70 | 0.96 | fmole/cell/hr | T1-TableSup4 | 11 |
| 1340 | lys release rate, T2=7 hr | $r_L(H=0.65)$ | 0.78 | 0.70 | 0.86 | fmole/cell/hr | T1-TableSup4 | 6 |
| 1340 | lys release rate, T2=6 hr | $r_L(H=0.73)$ | 0.53 | 0.45 | 0.61 | fmole/cell/hr | T1-TableSup4 | 8 |
| 1340 | lys release rate, T2=5.4hr | $r_L(H=0.80)$ | 0.38 | 0.32 | 0.45 | fmole/cell/hr | T1-TableSup4 | 4 |
| 1340 | lys release rate, T2=4 hr | $r_L(H=1.07)$ | 0.08 | 0.05 | 0.11 | fmole/cell/hr | T1-TableSup4 | 3 |

### Live L-A+ releases hypoxanthine upon lysine limitation

Metabolites can be released by live cells or leaked from dead cells. We want to distinguish between live versus dead release for the following reasons. First, if death rate were to evolve to be slower, then live release would predict increased metabolite supply whereas dead release would predict the opposite. Second, dead release would imply non-specific release and thus cell-cell interactions may be highly complex. Finally, leakage from dead cells is thermodynamically inevitable, whereas active release of costly molecules would require an evolutionary explanation.

Hypoxanthine is likely released by live *L*^*−*^*A*^*+*^. In the absence of lysine (Methods, “Starvation release assay”), we tracked the dynamics of live and dead *L*^*−*^*A*^*+*^ (Fig 5A, magenta and grey), and of hypoxanthine accumulation (Fig 5A, lavender). If live cells released hypoxanthine, then hypoxanthine should increase linearly with live cell density integrated over time (i.e. the sum of live cell density*hr, Fig 5B Left), and the slope would represent the live release rate (fmole/cell/hr). If cells released hypoxanthine upon death, then hypoxanthine should increase linearly with dead cell density, and the slope would represent the amount of metabolite released per cell death (Fig 5B Right). Since the live release model explained our data better than the dead release model (Fig 5B), hypoxanthine was likely released by live cells during starvation. In lysine-limited chemostats, we could not use dynamics to distinguish live from dead release (note the mathematical equivalence between Eqs. 8a and 8b in Methods “Quantifying phenotypes in chemostats”). Instead, we harvested cells and chemically extracted intracellular metabolites (Methods, “Extraction of intracellular metabolites”). Each *L*^*−*^*A*^*+*^ cell on average contained 0.12 (+/−0.02, 95% CI) fmole of hypoxanthine (Methods, “HPLC”). If hypoxanthine was released by dead cells (~10^5^ dead cells/ml, Fig 3A), we should see 0.012 μM instead of the observed ~10 μM hypoxanthine in the supernatant (Fig 3B). Thus, hypoxanthine was likely released by live *L*^*−*^*A*^*+*^ in chemostats.

**Fig. 5.**
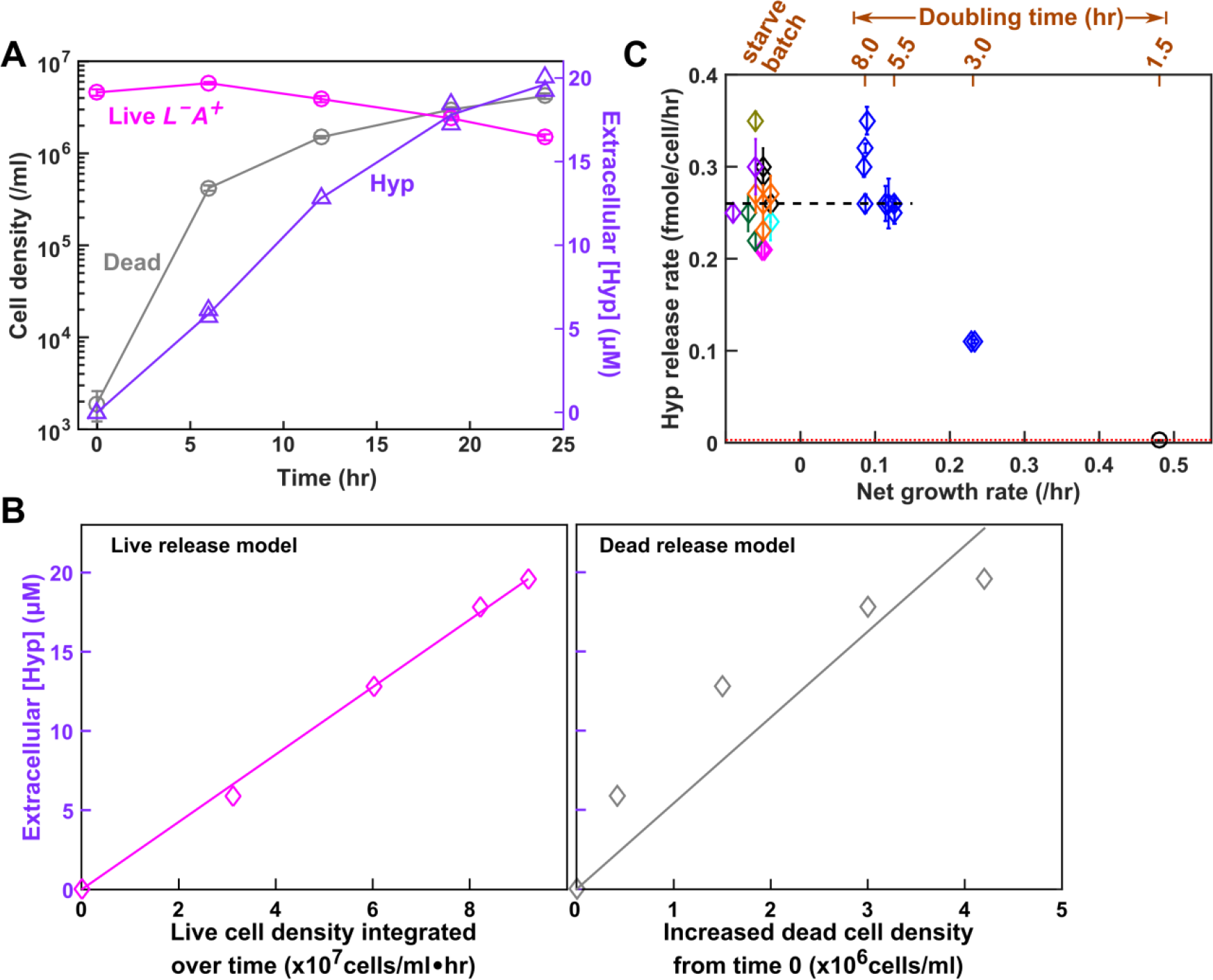
Hypoxanthine is released by live *L*^*−*^*A*^*+*^ upon limitation for lysine. (**A**) Hypoxanthine release during lysine starvation. Exponential *L*^*−*^*A*^*+*^ cells were washed free of lysine, and diluted into SD without lysine. Live and dead population densities and supernatant hypoxanthine concentrations were measured over time (Methods, “Flow cytometry”; the yield bioassay in “Bioassays”). (**B**) The “live release” model fits data better than the “dead release” model. (Left) If hypoxanthine was released by live cells at a constant rate, then hypoxanthine concentration should scale linearly against live cell density integrated over time. (Right) If hypoxanthine was released upon cell death, then hypoxanthine concentration should scale linearly against dead cell density. Live release model has better linearity than dead release model, and therefore hypoxanthine is likely released by live cells. The release rate measured in the left panel of **B** was used in Models i and ii. (**C**) Hypoxanthine release rate as a function of growth rate. Hypoxanthine release rates were plotted for *L*^*−*^*A*^*+*^ in the presence of excess lysine (black circle on the right), in lysine-limited chemostats with doubling times ranging from 3 hr to 8hr (blue; Fig 5-Figure Supplement 1 blue), and during lysine starvation (plotted at <0 net growth rate, different colors indicating experiments on different days). The black dashed line marks the average release rate measured from starvation up to 5.5 hr chemostats (Table 1) which was used in Model iii. Hypoxanthine release rate in excess lysine was lower than the detection limit (i.e. <0.003 fmole/cell/hr marked by the red dotted line; Methods “Measuring the upper bound of release rate in excess metabolites”). Release rates of *L*^*−*^*A*^*+*^ are in Table 1-Table Supplement 3.

Hypoxanthine release rates of *L*^*−*^*A*^*+*^ are similar in lysine-limited chemostats mimicking the CoSMO environments (Methods “Quantifying phenotypes in chemostats” Eq. 13; Fig 5-Figure Supplement 1) versus during starvation (Fig 5C). Thus, we used the average hypoxanthine release rate (Fig 5C black dashed line; Table 1) in Model iii. Note that release rates declined in faster-growing cultures (≤3-hr doubling; Fig 5C), but we did not use these data since CoSMO did not grow that fast.

### *A*^*−*^*L*^*+*^ intracellular lysine content and lysine release rate vary with the environment

Lysine is likely released by live *A*^*−*^*L*^*+*^. When we measured lysine release from starving *A*^*−*^*L*^*+*^ cells (Fig 6-Figure Supplement 1A), a model assuming live release and a model assuming dead release generated similar matches to experimental dynamics (Fig 6-Figure Supplement 1B and C). However, after measuring intracellular lysine content, we concluded that dead release was unlikely since each dead cell would need to release significantly more lysine than that measured inside a cell to account for supernatant lysine concentration, especially during the early stage of starvation (Fig 6-Figure Supplement 2B).

Lysine release rate of *A*^*−*^*L*^*+*^ is highly sensitive to the growth environment (Fig 6B, details in Fig 6-Figure Supplement 6). Release rate in 7~8 hr chemostat were ~60% more than that during starvation. Lysine release rate rapidly declined as hypoxanthine became more available (i.e. as growth rate increased, Fig 6B). Variable release rate could be due to variable intracellular lysine content: lysine content per cell increased by several-fold upon removal of hypoxanthine (from 2.9 fmole/cell to ~19 fmole/cell; Fig 6A black dotted line), and leveled off at a higher level in 8-hr chemostats than during starvation (Fig 6A). We incorporated variable lysine release rate in Model iii (Table 1).

**Fig. 6.**
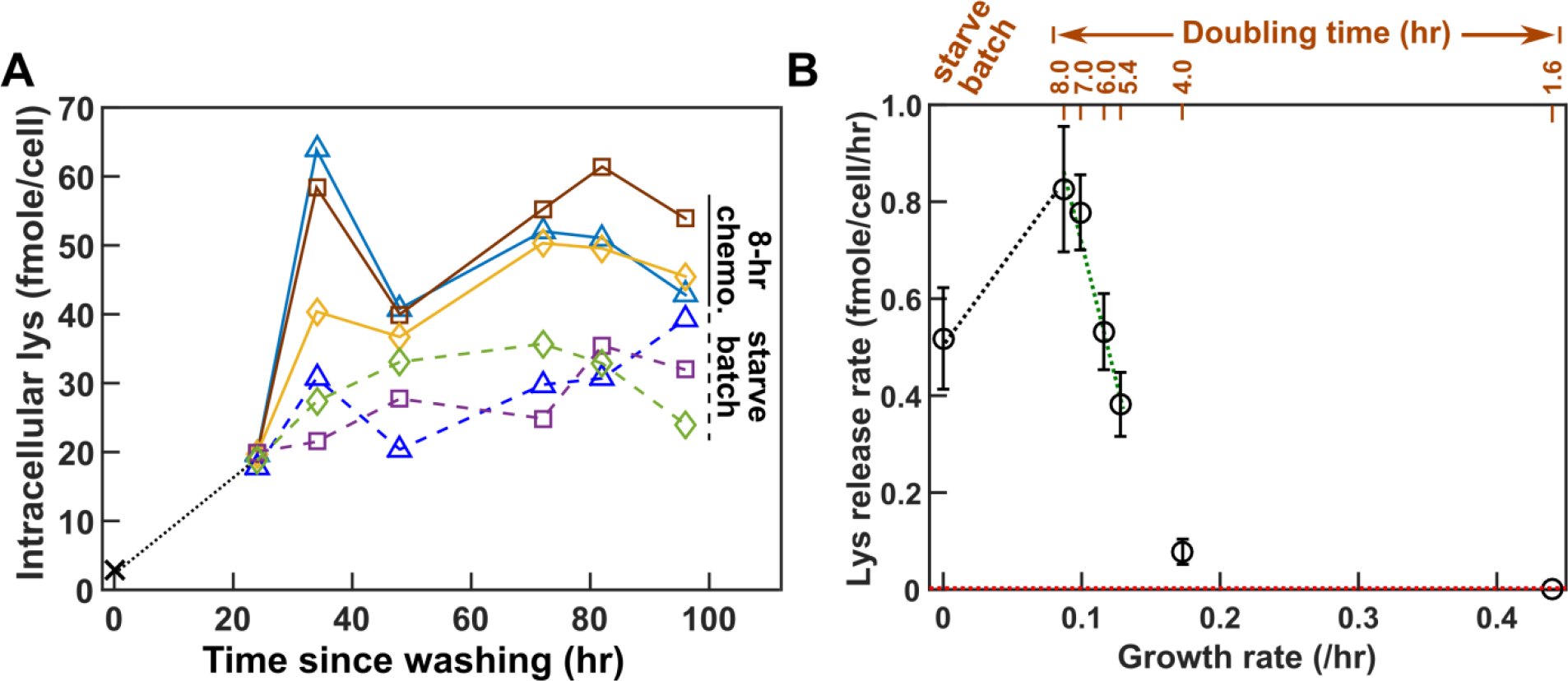
Intracellular lysine content and lysine release rate of *A*^*−*^*L*^*+*^ vary with environment. (**A**) Intracellular lysine content increases upon hypoxanthine limitation. *A*^*−*^*L*^*+*^ cells grown to exponential phase in SD plus excess hypoxanthine were washed and diluted into SD at time zero. Cells were either starved further (“starve batch”, dashed lines) or inoculated into hypoxanthine-limited chemostats after 24 hrs of pre-starvation (“8-hr chemo.”, solid lines). At various times, cells were harvested, and intracellular lysine was extracted (Methods “Extraction of intracellular metabolites”) and measured via yield bioassay (Methods “Bioassays”). Intracellular lysine content increased by six-fold during the 24-hour pre-starvation, even though the average cell size increased by only ~20% (Fig 6-Figure Supplement 3). Intracellular lysine content continued to increase, reaching a higher level in 8-hr chemostats than in starvation. Different colors represent different experiments. (**B**) Lysine release rate varies with the environment. Lysine release rates were quantified for cells at different growth rates (e.g. Fig 6-Figure Supplement 1; doubling times of chemostats marked above) and during starvation. Phenotypes were measured over a similar time window as for CoSMO growth rate measurement to ensure similar evolutionary effects (Fig 3-Figure Supplement 4). Means and their 2 SEM (standard error of mean) were plotted. Lysine release rate in an exponential batch culture was below the level of detection (red dotted line = 0.003 fmole/cell/hr; Methods “Measuring the upper bound of release rate in excess metabolites”). The green dotted regression line was used in analytical calculation (Eq. 16 in Methods “Calculating steady state community growth rate”), while both the black and the green dotted regression lines were used in spatial simulations. Lysine release rates were summarized in Table 1-Table Supplement 4, and plotted in greater detail in Fig 6-Figure Supplement 6.

### Parameters measured in community-like environments enable prediction of CoSMO growth rate

Death rates, which could affect CoSMO growth rate (Methods, Eq. 16), are also sensitive to the environment. We measured death rates in chemostats (Methods, “Quantifying phenotypes in chemostats” Eq. 11 or Eq. 14), and found them to be distinct from the death rates in zero or excess metabolite (Fig 7-Figure Supplement 1). Since death rates were relatively constant in chemostats mimicking the CoSMO environments (Fig 7-Figure Supplement 1, blue lines), we used the averaged values in Model iii (Table 1; Table 1-Table Supplement 3; Table 1-Table Supplement 4).

Our chemostat-measured model parameters are internally consistent: Mathematical models of *L*^*−*^*A*^*+*^ in lysine-limited chemostat (Fig 3-Code Supplement 3) and of *A*^*−*^*L*^*+*^ in hypoxanthine-limited chemostat (Fig 6-Code Supplement 1) captured experimental observations (Fig 3-Figure Supplement 7; Fig 6-Figure Supplement 5).

Using parameters measured in chemostats (Table 1), model prediction quantitatively matches experimental results. Experimentally, since *L*^*−*^*A*^*+*^ mutants quickly took over well-mixed CoSMO (red in Fig 7-Figure Supplement 2A ^33^), we grew CoSMO in a spatially-structured environment so that fast-growing mutants were spatially confined to their original locations and remained minority (red in Fig 7-Figure Supplement 2B). Spatial CoSMO growth rates measured under a variety of experimental setups (e.g. agarose geometry and initial total cell density) remained consistent (0.11± 0.01/hr; Fig 7 purple; Fig 7-Figure Supplement 4). In Model iii based on chemostat-measured parameters (Table 1), an analytical formula (Eq. 16 in Methods) and spatial CoSMO simulations both predicted CoSMO growth rate to be 0.10 ± 0.01/hr (Fig 7 green and brown). Thus, chemostat parameters allowed our model to quantitatively explain experimental CoSMO growth rate (Fig 7 green and brown versus purple).

**Fig. 7.**
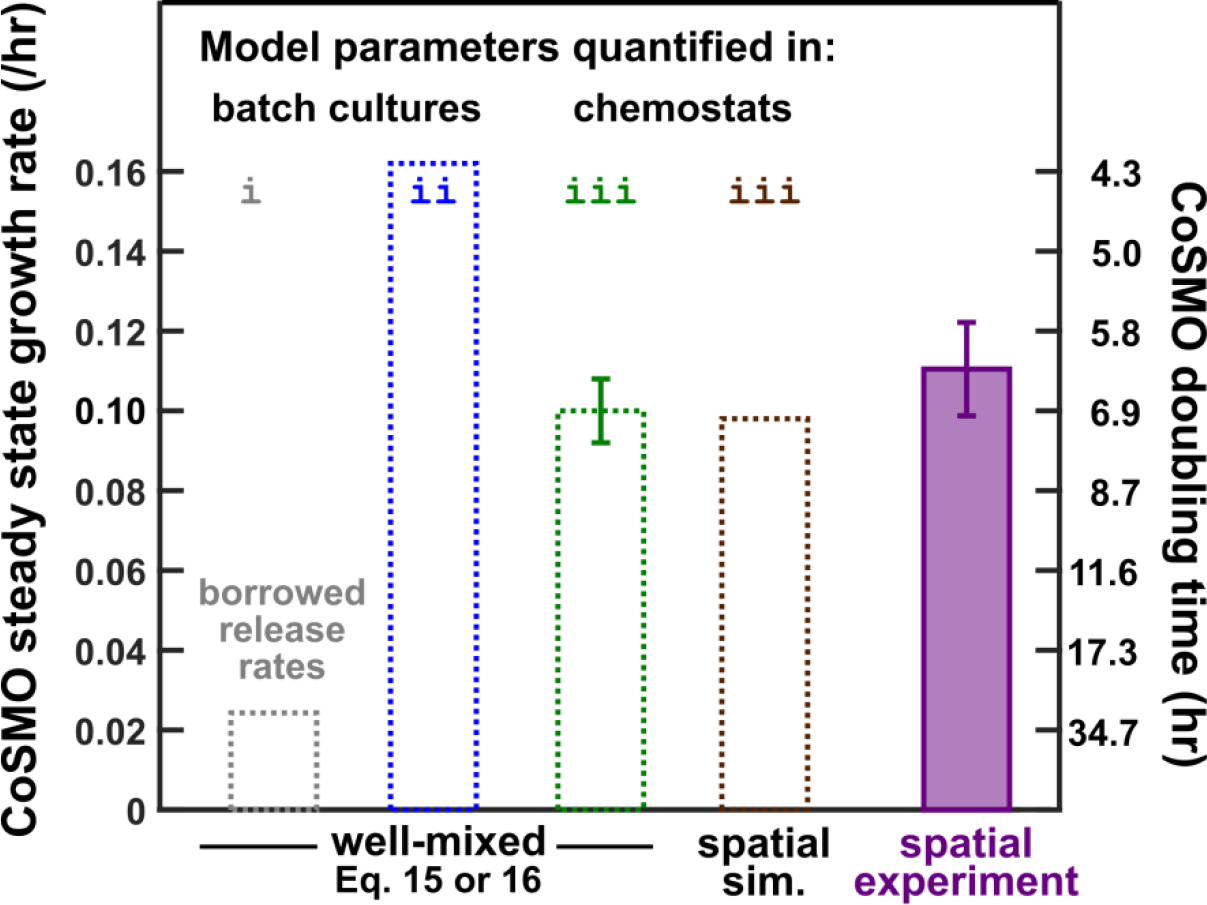
Steady state CoSMO growth rate can be predicted from parameters measured in chemostats. Steady state growth rate of well-mixed CoSMO was calculated from Eq. 15 or Eq. 16 (with nearly identical results) when we “borrowed” release rates from a different strain background in starved batch culture (grey; Model i in Fig 1C), when all parameters were measured from the correct strain background in starved batch cultures (blue; Model ii in Fig 1C), and when all parameters were measured from the correct strain background in chemostats (green; Model iii). A partial differential equation model (i.e. the spatial version of Model iii) was simulated for spatial CoSMO under various experimental configurations, yielding similar predictions on spatial CoSMO growth rate (Fig 7-Figure Supplement 3). The average value was plotted here (brown). In simulations, CoSMO grew at a similar rate in a spatially-structured environment as in a well-mixed environment. This is because in spatial CoSMO, concentrations of metabolites eventually became uniform in the agarose and in the community (Fig 7-Figure Supplement 5), and because the two strains formed small (tens of microns) patches that were inter-mixed ^24^. The green error bar (95% confidence interval) was calculated from uncertainties in parameter estimations (i.e. two standard errors of mean in Table 1) via the method of error propagation (Method “Calculating steady state community growth rate”). Under various experimental configurations (Methods, “Quantifying spatial CoSMO growth dynamics”), steady state CoSMO growth rates were similar, and the average value and two standard deviations from 11 independent experiments were plotted (purple).

## Discussions

Microbial communities are often complex, and thus it is difficult to understand how community-level properties emerge from interactions among member species. Here, using a highly simplified community growing in a well-controlled environment, we illustrate challenges in quantitative modeling and means to overcome them.

Even when interactions are engineered and thus genetic determinants are known, interaction mediators can be non-trivial to identify. In CoSMO, we previously thought that adenine was released by *L*^*−*^*A*^*+*^, whereas in reality, hypoxanthine and inosine are released (Fig 2). Fortuitously, hypoxanthine but not inosine affects *A*^*−*^*L*^*+*^ growth (Fig 2-Figure Supplement 1). Otherwise, we might be forced to quantify how hypoxanthine and inosine, in isolation and in different combinations, might affect *A*^*−*^*L*^*+*^. Even though faster growth of *A*^*−*^*L*^*+*^ in adenine than in hypoxanthine does not affect our prediction of CoSMO growth rate (Fig 2-Figure Supplement 2; Fig 1-Figure Supplement 1), it could affect predictions on other community properties.

Another obstacle for model building and testing is rapid evolution during quantification of species phenotypes and community properties. For *L*^*−*^*A*^*+*^, mutants pre-exist at a low frequency, but they can grow several-fold faster than the ancestor (Fig 3-Figure Supplement 2). Consequently, a population will remain largely (>90%) ancestral only for the first 24 hours in the well-mixed chemostat environment (Fig 3). A short measurement time window poses additional challenges if, for example, the released metabolite has not accumulated to a quantifiable level. Takeover by fast-growing mutants can be impeded by growing cells in a spatially-structured environment (Fig 7-Figure Supplement 2). For *A*^*−*^*L*^*+*^, mutants are generated from ancestral cells at an extremely high rate during phenotype quantification (Methods, “Evolutionary dynamics of mutant *A-L*+”; Fig 3-Figure Supplement 4). Since similar levels of mutants also accumulate in CoSMO (Fig 3-Figure Supplement 4B), we can account for evolutionary effects by using similar quantification time windows for *A*^*−*^*L*^*+*^ phenotypes and for CoSMO growth rate. Thus, unless we are extremely careful, we may not even know what we are measuring. The evolutionary phenomena here are likely common: Given the large population size of microbial populations, pre-existing mutants ^35^ could quickly take over upon drastic environmental shifts. Eventually, evolving communities could be modeled, although replicate communities could evolve divergently ^33^.

Many mathematical models “borrow” parameters from a different strain or even a different species. This practice can work for qualitative models or even quantitative models where parameter variations do not affect predictions. In the case of CoSMO, release rates from two genetic backgrounds differed by several-fold (Fig 1-Table Supplement 2). Thus, borrowing parameters could be problematic for quantitative modeling.

Most mathematical models assume invariant parameters. As we have demonstrated here, phenotypes (e.g. metabolite consumption per birth, metabolite release rate, and death rate) measured in zero or excess metabolite can differ dramatically from those measured under metabolite limitation (Fig 4; Fig 4-Figure Supplement 1; Fig 5C; Fig 6B; Fig 7-Figure Supplement 1). Furthermore, even within the range of metabolite limitation experienced by CoSMO (doubling times of 5.4~8 hrs), lysine release rate varies by as much as two-fold (Fig 6B), which could be caused by variable intracellular metabolite concentrations (Fig 6A). Based on parameters measured in chemostats (including variable lysine release rate), Model iii prediction quantitatively agrees with experimental results (Fig 7).

Our work illustrates how a mathematical model can synergize with quantitative measurements. A model suggests which parameters need to be carefully measured. For example for spatial CoSMO growth rate, parameters such as diffusion coefficients are not critical (Fig 7-Figure Supplement 3), but release rates and consumption are (Methods, Eq. 15). Once key parameters have been measured, model-experiment discrepancy suggests important missing pieces. When predicting CoSMO growth rate, a missing piece is how phenotype parameters are affected by the growth environment.

Physiology, ecology, and evolution are clearly intertwined. Physiology impacts not only organisms’ evolutionary success, but also their ecological interactions with other community members. For example in quorum sensing, bacteria at a high cell density enter a different physiological state and release new chemical compounds or enzymes ^37^. As another example, computational work suggests that metabolite secretion can be costly or costless to the secretors depending on the environmental context ^36^. Thus, it is not surprising that quantitative differences in physiology (e.g. growth rate) can impact the strength of an ecological interaction (metabolite release and consumption). Moreover, evolutionary changes and ecology affect organismal physiology. Thus, extending the “ecology-evolution feedback” concept ^38–44^, we advocate an integrated view of evolution, ecology, and physiology.

## Methods

### Strains and growth medium

We constructed CoSMO in the RM11 background due to its lower rate of mitochondrial mutations ^27^ compared to the S288C background used in our earlier studies ^45^. Thus, phenotypes measured here differed from those measured in strains of the S288C background ^17,24^ We introduced desired genetic modifications into the ancestral RM11 background via transformation ^46,47^ (Fig 1-Table Supplement 1). Strains were stored at −80°C in 15% glycerol.

We used rich medium YPD (10 g/L yeast extract, 20 g/L peptone, 20 g/L glucose) in 2% agar plates for isolating single colonies. Saturated YPD overnight liquid cultures from these colonies were then used as inocula to grow exponential cultures. To prevent purines from being yield-limiting, we supplemented YPD with 100 μM hypoxanthine for *A*^*−*^*L*^*+*^ cells. We sterilized YPD media by autoclaving. YPD overnights were stored at room temperature for no more than 4~5 days prior to experiments.

We used defined minimal medium SD (6.7 g/L Difco™ yeast nitrogen base w/o amino acids, 20 g/L glucose) for all experiments ^48^, with supplemental metabolites as noted ^48^. To achieve higher reproducibility, we sterilized SD media by filtering through 0.22 μm filters. To make SD plates, we autoclaved 20 g/L Bacto™ agar or agarose in H2O, and after autoclaving, supplemented equal volume of sterile-filtered 2XSD. CoSMO steady state growth rates on agar (which contains trace contaminants of metabolites) and agarose generate similar results.

### Strain culturing and preconditioning

All culturing, unless otherwise noted, was performed at 30°C in a well-mixed environment where culture tubes were inserted side-ways into a roller drum (New Brunswick Scientific, Model TC-7). *L*^*−*^*A*^*+*^ cells were pre-grown to exponential phase (OD_600_ generally less than 0.4 in 13mm culture tubes, or < 2.8×10^7^ cells/ml) in SD supplemented with excess (164 μM) lysine and washed 3~5 times with SD. In microscopy assays, when noted, we starved *L*^*−*^*A*^*+*^ cells for 4 hours to deplete intracellular lysine storage. Otherwise, we did not prestarve *L*^*−*^*A*^*+*^. *A*^*−*^*L*^*+*^ cells were pre-grown to exponential phase in SD supplemented with excess hypoxanthine (100 μM) or excess adenine (108 μM) as noted, washed 3~5 times with SD, and pre-starved in SD for 24 hours to deplete cellular purine storage. We pre-starved *A*^*−*^*L*^*+*^ to reduce CoSMO growth lag (Fig 1-Figure Supplement 2), thus facilitating quantification of CoSMO growth rate. To be consistent, we also pre-starved *A*^*−*^*L*^*+*^ during phenotype quantification.

### Flow cytometry

We prepared fluorescent bead stocks (ThermoFisher Cat R0300, 3 μm red fluorescent beads). Beads were autoclaved in a factory-clean glass tube, diluted into sterile 0.9% NaCl, and supplemented with sterile-filtered Triton X-100 to a final 0.05% (to prevent beads from clumping). We sonicated beads and kept them in constant rotation to prevent settling. We quantified bead concentrations by counting beads via hemacytometer and Coulter counter. Final bead stock was generally 4~8×10^6^/ml.

Culture samples were diluted to OD 0.01~0.1 (7×10^5^~7×10^6^/ml) in Milli-Q H2O in unautoclaved 1.6ml Eppendorf tubes. 90 μl of the diluted culture sample was supplemented with 10 μl bead stock and 2 μl of 1 μM ToPro 3 (Molecular Probes T-3605), a nucleic acid dye that only permeates compromised cell membranes (dead cells). Sample preparation was done in a 96-well format for high-throughput processing.

Flow cytometry of the samples was performed on Cytek DxP Cytometer equipped with four lasers, ten detectors, and an autosampler. Fluorescent proteins GFP, Citrine, mCherry, TagBFP-AS-N (Evrogen), and ToPro are respectively detected by 50 mW 488 nm laser with 505/10 (i.e. 500~515nm) detector, 50 mW 488nm laser with 530/30 detector, 75 mW 561nm Laser with 615/25 detector, 50 mW 408nm laser with 450/50 detector, and 25mW 637nm laser with 660/20 detector. Each sample was run in triplicates and individually analyzed using FlowJo^®^ software to identify numbers of events of beads, dead cells, and various live fluorescent cells. Densities of various populations were calculated from the cell:bead ratios. We then calculated the mean cell density from triplicate measurements, with the coefficient of variation generally within 5%~10%.

### HPLC

All HPLC measurements were done on a Shimadzu Nexera X2 series Ultra High Performance Liquid Chromatography (UHPLC) system. All supernatant samples were filtered (0.22 μm filter). For standards, we made a high-concentration solution, filtered it, and stored it at −80 °C. Prior to an HPLC run, we diluted the stock to various concentrations in filtered H_2_O.

To quantify lysine, 100 μl sample was loaded into an Agilent 250 μl pulled point glass small volume insert (Part No: 5183-2085), which was then placed inside a Shimadzu 1.5 ml 12× 32 mm autosampler vial (Part No: 22845450-91). This vial was then placed into an autosampler (Nexera X2 SIL-30AC). Prior to injection into the column, samples were derivatized at 25°C with freshly-made derivatization reagents in the autosampler using a programmed mixing method as following. 7.5 μl of sample was removed and placed into a separate reaction small volume insert and vial. Next, 45 μl of mercaptopropionic acid (10 μl per 10 ml 0.1 M sodium borate buffer, pH 9.2) and 22 μl of o-phthaladehyde (10 mg per 5 ml 0.1 M sodium borate buffer, pH 9.2) were added to this vial, mixed, and incubated for 1 minute. 10 μl of 9-fluorenyl methyl chloroformate (4 mg per 20 ml acetonitrile, HPLC grade) was then added and the sample was re-mixed and incubated for 2 minutes. Finally, 10 μl of the reaction mixture was injected onto Phenomenex Kinetex^®^ 2.6 μm EVO C18 100 Å LC Column (150 × 3.0 mm, Part No: 00F-4725-Y0) fitted with a SecurityGuard™ ULTRA Holder for UHPLC Columns (2.1 to 4.6 mm, Part No AJ0-9000) and a SecurityGuard™ ULTRA cartridge (3.0 mm internal diameter, Part No: AJ0-9297). SecurityGuard™ ULTRA cartridge (pre-column) was periodically replaced in the event of pressure reading exceeding the manufacturer’s specifications.

Compounds were eluted from the column using a gradient of HPLC-grade Solution A (73 mM Potassium Phosphate, pH 7.2) and Solution B (50:50 acetonitrile/methanol). Solution A was filtered through a 0.2 μm filter prior to use. The % solution B follows the following program: 0-2 minutes 11%, 2-4 minutes 17%, 4-5.5 minutes 31%, 5.5-10 minutes 32.5%, 10-12 minutes 46.5%, and 12 - 15.5 minutes 55%. The flow rate is maintained at 0.1 ml/minute. The column was then flushed with 100% solution B for 5 minutes, and re-equilibrated for 5 minutes with 11% solution B at 0.8 ml/min. The column was maintained at a running temperature of 35 °C in a Nexera X2 CTO-20A oven. Absorbance measurements at 338 nm were measured using a high sensitivity flow cell for a SPD-M30A UV-Vis detector.

For purines, we used the above protocol without the derivatization steps. Instead, 5~10 μl sample was directly injected onto the column.

### Bioassays

We used a yield bioassay for relatively high metabolite concentrations (≥ 5 μM for lysine and ≥2 μM for hypoxanthine). For lower concentrations, we used a rate bioassay with a sensitivity of 0.1 μM for both lysine and hypoxanthine (Fig 3-Figure Supplement 1). When necessary, we diluted the sample to get into the assay linear range.

In the yield bioassay, 75 μl sample filtered through a 0.2 μm filter was mixed with an equal volume of a master mix containing 2× SD (to provide fresh medium) as well as tester cells auxotrophic for the metabolite of interest (~1×10^4^ cells/ml, WY1335 for lysine or WY1340 for hypoxanthine) in a flat-bottom 96-well plate. We then wrapped the plate with parafilm and allowed cells to grow to saturation at 30°C for 48 hrs. We re-suspended cells using a Thermo Scientific Teleshake (setting #5 for ~1 min) and read culture turbidity using a BioTek Synergy MX plate reader. Within each assay, SD supplemented with various known concentrations of metabolite were used to establish a standard curve that related metabolite concentration to final turbidity (e.g. Fig 2-Figure Supplement 1A). From this standard curve, the metabolite concentration of an unknown sample could be inferred.

The rate bioassay was used for samples with low metabolite concentrations. For example, to measure lysine concentration in a lysine-limited chemostat, we mixed 150 μl filtered sample with an equal volume of master mix containing 2× SD and *L*^*−*^*A*^*+*^ tester cells (~1×10^4^ cells/ml) in a flat-bottom 96-well plate. As our tester strain for lysine, we used an evolved clone (WY 2270) isolated after *L*^*−*^*A*^*+*^ had grown for tens of generations under lysine limitation. This clone displayed increased affinity for lysine due to an *ecm21* mutation and duplication of Chromosome 14. Growth rates of the tester strain in SD supplemented with various known concentrations of lysine and in the unknown sample were measured using a microscopy assay (Methods, “Microscopy quantification of growth phenotypes”). The growth rate of WY 2270 tester cells scaled linearly with lysine concentrations up to 1 μM (Fig 3-Figure Supplement 1A). Similarly, for hypoxanthine, we used an evolved *A*^*−*^*L*^*+*^ strain (WY1600) as the tester strain. The linear range was up to ~0.3 μM (Fig 3-Figure Supplement 1B). From the standard curve, we could infer the metabolite concentration of a sample.

### Extraction of intracellular metabolites

To extract intracellular metabolites, we poured a cell culture onto a 0.45 μm nitrocellulose membrane filter (BioRad, Cat# 162-0115) in a reusable filtration device (Southern Labware Product FHMA25, glass microanalysis 25mm vacuum filter holder with 15mL funnel), applied vacuum to drain the supernatant, transferred the filter into extraction solution (40% acetonitrile, 40% methanol, 20% water), vortexed to dislodged cells, and then removed the filter. This sequence was carried out as rapidly as possible (<10 seconds). We then flash-froze the extraction solution in liquid nitrogen, and allowed it to thaw at −20°C. After thawing, we subjected the solution through 5 rounds of the following: vortexing for 1 minute, and incubating on ice for 5 minutes between each vortexing. We then spun down the solution in a refrigerated centrifuge for 10 min at 14,000 rpm to pellet ghost cells as well as any membrane filter bits that may have disintegrated into the extraction solution. We transferred the supernatant containing soluble cell extract to a new tube. In order to make sure that all soluble components were extracted, we re-suspended the cell pellet in a half-volume of fresh extraction solution and subjected cells through another round of the same procedure (flash-freezing, five rounds of vortexing-ice incubation, and centrifugation). We then removed the supernatant, and added it to the original supernatant. We then dried off the extraction solution in a centrifugal evaporator and re-suspended soluble components in water. This resultant solution could then be assayed for metabolite concentrations. When properly dried, extracts did not contain inhibitors that might interfere with bioassays (Fig 2-Figure Supplement 3).

For *L*^*−*^*A*^*+*^, cells from 19 ml cultures (4×10^5^ ~ 4×10^6^ cells/ml) were re-suspended in 3 ml extraction buffer. One third of the sample was further processed, and extracted metabolites were re-suspended in 0.5 ml water. For *A*^*−*^*L*^*+*^, metabolites from 1~5 ml cultures (1~6×10^6^ cells/ml) were extracted and re-suspended in 1 ml water.

### Microscopy quantification of growth phenotypes

See ^28^ for details on microscopy and experimental setup, method validation, and data analysis. Briefly, cells were diluted to low densities to minimize metabolite depletion during measurements. Dilutions were estimated from culture OD measurement to result in 1000~5000 cells inoculated in 300 μl SD medium supplemented with different metabolite concentrations in wells of a transparent flat-bottom microtiter plate (e.g. Costar 3370). We filled the outermost wells with water to reduce evaporation.

Microtiter plates were imaged periodically (every 0.5~2 hrs) under a 10x objective in a Nikon Eclipse TE-2000U inverted fluorescence microscope. The microscope was connected to a cooled CCD camera for fluorescence and transmitted light imaging. The microscope was enclosed in a temperature-controlled chamber set to 30°C. The microscope was equipped with motorized stages to allow z-autofocusing and systematic xy-scanning of locations in microplate wells, as well as motorized switchable filter cubes capable of detecting a variety of fluorophores. Image acquisition was done with an in-house LabVIEW program, incorporating bright-field autofocusing ^28^ and automatic exposure adjustment during fluorescence imaging to avoid saturation. Condensation on the plate lid sometimes interfered with autofocusing. Thus, we added a transparent “lid warmer” on top of our plate lid ^28^, and set it to be 0.5°C warmer than the plate bottom, which eliminated condensation. We used an ET DsRed filter cube (Exciter: ET545/30x, Emitter: ET620/60m, Dichroic: T570LP) for mCherry-expressing strains, and an ET GFP filter cube (Exciter: ET470/40x, Emitter: ET525/50m, Dichroic: T495LP) for GFP-expressing strains.

Time-lapse images were analyzed using an ImageJ plugin Bioact2 ^28^. Bioact2 measured the total fluorescence intensity of all cells in an image frame after subtracting the background fluorescence from the total fluorescence. A script plotted background-subtracted fluorescence intensity over time for each well to allow visual inspection. If the dynamics of four positions looked similar, we randomly selected one to inspect. In rare occasions, all four positions were out-of-focus and were not used. In a small subset of experiments, a discontinuous jump in data appeared in all four positions for unknown reasons. We did not calculate rates across the jump. Occasionally, one or two positions deviated from the rest. This could be due to a number of reasons, including shift of focal plane, shift of field of view, black dust particles, or bright dust spots in the field of view. The outlier positions were excluded after inspecting the images for probable causes. If the dynamics of four positions differed due to cell growth heterogeneity at low concentrations of metabolites, all positions were retained.

We normalized total intensity against that at time zero, and averaged across positions. We calculated growth rate over three to four consecutive time points, and plotted the maximal net growth rate against metabolite concentration (e.g. Fig 2-Figure Supplement 2). If maximal growth rate occurred at the end of an experiment, then the experimental duration was too short and data were not used. For *L*^*−*^*A*^*+*^, the initial stage (3~4 hrs) residual growth was excluded from analysis. For *A*^*−*^*L*^*+*^, since cells had already been pre-starved, fluorescence intensity did not continue to increase in the absence of supplements.

For longer *A*^*−*^*L*^*+*^ imaging (30+ hr), we observed two maximal growth rates at low hypoxanthine concentrations (e.g. ~0.4 μM) possibly due to mutant clones (Fig 3-Figure Supplement 3). We used the earlier maximal growth rate even if it was lower than the later maximal growth rate, since the latter was probably caused by faster-growing mutants.

### Chemostat culturing

We have constructed an eight-vessel chemostat with a design modified from ^49^. For details of construction, modification, calibration, and operation, see ^50^.

For *L*^*−*^*A*^*+*^, due to rapid evolution, we tried to devise experiments so that live and dead populations quickly reached steady state. Two conditions seemed to work well. In both, we first calculated the expected steady state cell density by dividing the concentration of lysine in the reservoir (20 μM) by fmole lysine consumed per new cell. Condition 1: Wash exponentially-growing cells to completely remove any extracellular lysine and inoculate the full volume (19 ml) at 100% of expected steady state density. Start chemostat to drip in lysine at the pre-specified flow rate. Condition 2: Wash exponentially growing cells to remove extracellular lysine and inoculate 50%~75% of the volume at 1/3 of the expected steady state density. Fill the rest of the 19ml vessel with reservoir media (resulting in less than the full 20 μM of starting lysine, but more than enough for maximal initial growth rate, ~10-15 μM). The two conditions yielded similar results (Fig 3). We predominantly used Condition 2.

We set the pump flow rate to achieve the desired doubling time *T* (19ml*ln(2)/T). We collected and weighed waste media for each individual culturing vessel to ensure that the flow rate was correct (i.e. total waste accumulated over time *t* was equal to the expected flow rate\**t*). We sampled cultures periodically to track population dynamics using flow cytometry (Methods, “Flow cytometry”), and filtered supernatant through a 0.45 μm nitrocellulose filter and froze the supernatant for metabolite quantification at the conclusion of an experiment (Methods, “Bioassays”). At the conclusion of an experiment, we also tested input media for each individual culturing vessel to ensure sterility by plating a 300 μl aliquot on an YPD plate and checking for growth after two days of growth at 30°C. If a substantial number of colonies grew (>5 colonies), the input line was considered contaminated and data from that vessel was not used.

*A*^*−*^*L*^*+*^ cells exponentially growing in SD+100 μM hypoxanthine were washed and prestarved for 24 hrs. We then filled the chemostat culturing vessel with starved cells in SD at 100% of the expected starting density and pumped in fresh medium (SD+ 20 μM hypoxanthine) to achieve the desired doubling time. Cultures were otherwise treated as described above for *L*^*−*^*A*^*+*^.

For most experiments, we isolated colonies from end time point and checked percent evolved (Methods, “Detecting evolved clones”). For *L*^*−*^*A*^*+*^, we only analyzed time courses where >90% of population remained ancestral. For *A*^*−*^*L*^*+*^, significant levels of mutants were generated before and throughout quantification (Fig 3-Figure Supplement 4). Since quantified phenotypes did not correlate strongly with % mutants (Fig 3-Figure Supplement 6) and since mutants accumulated similarly during chemostat measurements and during CoSMO growth rate measurements (Fig 3-Figure Supplement 4B), we used the time window for CoSMO growth rate quantification (~96 hrs) in *A*^*−*^*L*^*+*^ chemostat experiments.

### Quantifying phenotypes in chemostats

We illustrate how we quantify release rate, consumption amount per birth, and death rate in chemostats, using *L*^*−*^*A*^*+*^ as an example. In a lysine-limited chemostat, live cell density [*L*^*−*^*A*^*+*^]_*live*_ is increased by birth (at a rate *b*_*L*_), and decreased by death (at a rate *d*_*L*_) and dilution (at a rate *dil*):

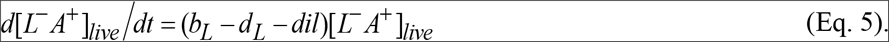

Dead cell density [*L*^*−*^ *A*^*+*^]_*dead*_ is increased by death and decreased by dilution

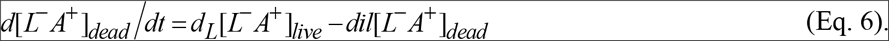

*L*, lysine concentration in the culturing vessel, is increased by the supply of fresh medium (at concentration *L*_0_), and decreased by dilution and consumption (with each birth consuming *c*_*L*_ amount of lysine).

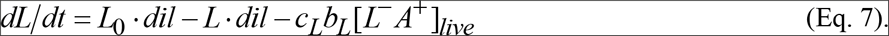

Finally, hypoxanthine concentration *A* is increased by release (either from live cells at *r*_*A*_ per live cell per hr or from dead cells at *r*_*A*,*d*_ per death), and decreased by dilution.

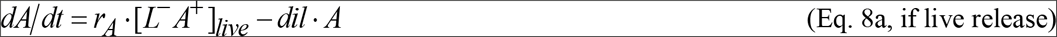

or

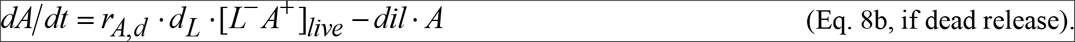

Note that at the steady state (denoted by subscript “*ss*”), net growth rate is equal to dilution rate (setting Eq. 5 to zero):

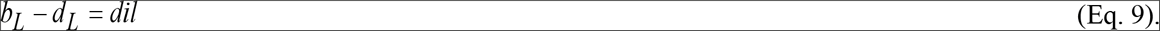

To measure metabolite consumed per cell at steady state, we set Eq. 7 to zero

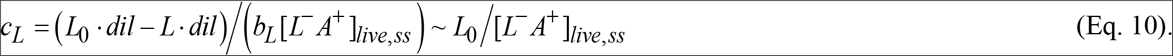

Here, the approximation holds because the concentration of lysine in chemostat (*L*) is much smaller than that in reservoir (*L*_*0*_) and because birth rate *b*_*L*_ is similar to dilution rate *dil*.

To measure death rate at steady state, we set Eq. 6 to zero, and get

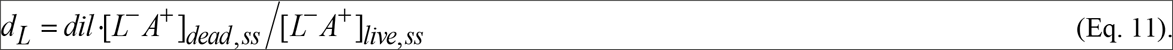

Thus, we can measure death rate by measuring the steady state dead and live population densities averaged over time.

To measure release rate at steady state, we can set Eq. 8a to zero and obtain:

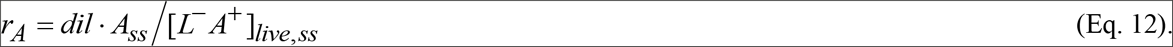

Alternatively, we can use both the pre-steady state and steady state chemostat dynamics to quantify release rate and death rate if these rates are constant. For release rate, we multiply both sides of Eq. 8a with *e*^*dil.t*^

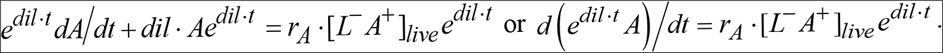

We have

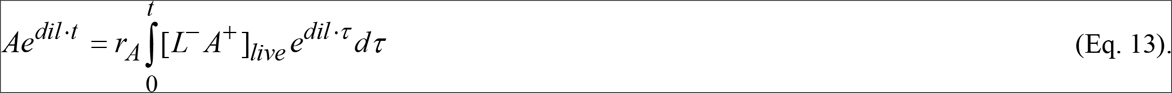

How do we calculate 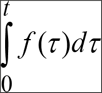 from experimental data? The value of integral is always zero at *t*=0. For each time point *t* + Δ*t*, the integral is the integral at the previous time point *t* (i.e. 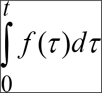 plus 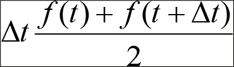. If we plot *Ae*^*dil.t*^ against 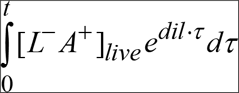, since the initial *A* is zero, we should get a straight line through origin with a slope of *r*_*A*_ (Fig 5-Figure Supplement 1, blue).

Similarly from Eq. 6, if death rate is constant, we have

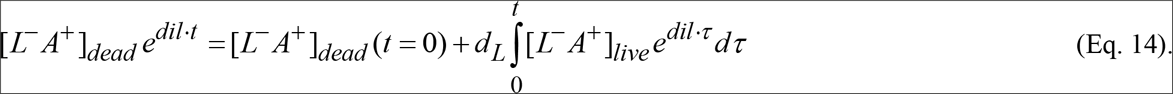

If we plot [*L*^*−*^*A*^*+*^]_*deade*_*e*^*dil.t*^ against 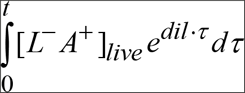, we should get a straight line with a slope of *d*_*L*_ (Fig 5-Figure Supplement 1, grey).

The two methods (using only the steady state data versus performing linear regression on the entire data range) yielded similar results. We have opted for the latter method since it takes advantage of pre-steady-state data.

### Detecting evolved clones

To detect evolved clones in an *L*^*−*^*A*^*+*^ culture, we diluted it to <1000 cells/ml and plated 300 μl on YPD plate and allowed colonies to grow for 2~3 days. We randomly picked 20~50 colonies to inoculate into YPD and grow saturated overnights. We diluted each saturated overnight 1:6000 into SD+164 μM lysine, and allowed cultures to grow overnight at 30°C to exponential phase. We washed cells 3x with SD, starved them for 4-6 hours to deplete vacuolar lysine stores, and diluted each culture so that a 50 μl spot had several hundred cells. We spotted 50 μl on SD plate supplemented with 1.5 μM lysine (10 spots/plate), and allowed these plates to grow overnight. When observed under a microscope, evolved cells with increased lysine affinity would grow into “microcolonies” of ~20~100 cells, while the ancestral genotype failed to grow (Fig 3-Figure Supplement 2C). Occasionally an intermediate phenotype was observed where smaller microcolonies with variable sizes formed, and this phenotype was counted as evolved as well. For a high-throughput version of this assay, we diluted YPD saturated culture 10,000x into SD and waited for 3 hrs at room temperature. We then directly spotted 50 μl on SD plates supplemented with 1.5 μM lysine. Ancestral cells formed ≤ 10 cell-clusters, but we could still clearly distinguish ancestor versus evolved clones.

To detect evolved clones in an *A*^*−*^*L*^*+*^ culture, we took advantage of the observation that evolved clones with improved affinity for hypoxanthine grew slowly when hypoxanthine concentration was high (Fig 3-Figure Supplement 3A). Similar fitness tradeoff has been observed for *L*^*−*^*A*^*+*^ ^33^ and in many other examples ^51–54^ From an *A*^*−*^*L*^*+*^ culture, we randomly picked colonies and made individual YPD overnights in a 96-well plate. We diluted YPD overnights 1:3,600 fold into SD+100 μM hypoxanthine or 108 μM adenine, and grew for 16~24 hr. Some of these cultures were not turbid while other cultures and the ancestor reached near saturation (Fig 3-Figure Supplement 3B). We considered these low-turbidity cultures as evolved, and they generally grew faster than the ancestor in low (0.4 μM) hypoxanthine (Fig 3-Figure Supplement 3A, compare blue, grey, and green against magenta).

### Starvation release assay

For *L*^*−*^*A*^*+*^, we washed exponential phase cells and diluted each sample to OD~0.1 to roughly normalize cell density. We took an initial cell density reading of each sample by flow cytometry, wrapped tube caps in parafilm to limit evaporation, and incubated in a rotator at 30°C. Prep time (from the start of washing to the initial cell density reading) took approximately two hours, during which time the majority of residual growth had taken place. At each time point we measured live and dead cell densities by flow cytometry, and froze an aliquot of supernatant where supernatant had been separated from cells by filtering through sterile nitrocellulose membrane. We concluded the assay after approximately 24 hours, generally aiming for time points every six hours. At the conclusion of the assay, we quantified hypoxanthine concentration for each sample using the yield bioassay (Methods, “Bioassays”). The slope of the linear regression of integrated live cell density over time (cells/ml*hr) versus hypoxanthine concentration (μM) gave us the release rate.

For *A*^*−*^*L*^*+*^, the starvation release assay was similar, except that the assay lasted longer with less frequent time points to accommodate the longer assay. Pre-growth in 108 μM Ade versus 100 μM hypoxanthine generated similar release rates, and thus we pooled the data.

### Evolutionary dynamics of mutant *A*^*−*^*L*^*+*^

Mutant *A*^*−*^*L*^*+*^ clones were alike, and they grew ~50% slower than the ancestor in excess hypoxanthine (Fig 3-Figure Supplement 3A). This has allowed us to rapidly quantify mutant abundance (Fig 3-Figure Supplement 3B; Methods, “Detecting evolved clones”). The high abundance of mutants during exponential growth is surprising, especially given the large (~50%) fitness disadvantage of mutants (Fig 3-Figure Supplement 3A). Whole-genome sequencing of a randomly-chosen evolved *A*^*−*^*L*^*+*^ clone (WY2447) revealed evidence for aneuploidy (Fig 3-Figure Supplement 3C; Methods “Genomic Analysis”). Assuming a chromosomal mis-segregation rate of 0.01/generation/cell and incorporating the fitness difference between ancestor and mutant in various hypoxanthine concentrations (Fig 3-Figure Supplement 5A), our mathematical models (Fig 3-Code Supplements 1 and 2) qualitatively captured experimental observations (Fig 3-Figure Supplement 5B and C). This extraordinarily high mutation rate is possibly due to an imbalance in purine intermediates in a purine auxotroph, and is in-line with the highest chromosomal mis-segregation rate observed in chromosome transmission fidelity mutants (up to 0.015/generation/cell) ^55^. In low concentrations of hypoxanthine (<1 μM), the fitness difference between mutant and ancestral *A*^*−*^*L*^*+*^ varied from −30% to 70% (right panel of Fig 3-Figure Supplement 5A), consistent with the dynamics of mutant *A*^*−*^*L*^*+*^ in chemostats.

Ancestral and evolved clones exhibited distinct phenotypes (Fig 3-Figure Supplement 3A and D). However, measured phenotype values were not significantly correlated with % mutants at the end of an experiment. This was due to the relatively narrow spread in % mutants and the relatively large measurement errors (Fig 3-Figure Supplement 6).

### Measuring consumption in batch cultures

To measure consumption in exponential cultures, we diluted exponentially growing cells to ~1×10^6^ cell/ml in SD supplemented with ~100μM metabolite, and measured cell density (Methods, “Flow cytometry”) and metabolite concentration (Methods, yield assay in “Bioassays”) every hour over six hours. For an exponential culture of size *N*(*t*) growing at a rate *g* while consuming metabolite M, we have

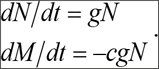

Thus, 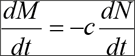. Integrating both sides, we have *M(t)*-*M(0)*=−*c(N(t)*-*N(0))*. Thus, if we plot M(t) against N(t), the slope is consumption per birth. We disregarded time points after which M had declined to less than 10 μM even though cells could still grow at the maximal growth rate.

We also measured consumption after cells fully “saturated” the culture and used intracellular stores for residual growth. We starved exponentially-growing cells (3-6 hours for *L*^*−*^*A*^*+*^, 24 hours for *A*^*−*^*L*^*+*^) to deplete initial intracellular stores and inoculated ~1×10^5^ cells/ml into various concentrations of the cognate metabolite up to 25 μM. We incubated for 48 hours and then measured cell densities by flow cytometry. We performed linear regression between input metabolite concentrations and final total cell densities within the linear range, forcing the regression line through origin. Consumption per birth in a saturated culture was quantified from 1/slope.

### Measuring the upper bound of release rate in excess metabolites

To measure release rate in an exponentially-growing population in excess metabolites, we note that

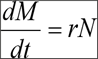

where *M* is metabolite concentration, *r* is the release rate, and *N* is live population density. Let *g* be growth rate, then after integration, we have

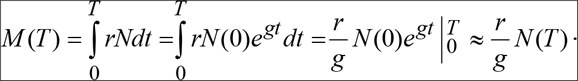

The approximation holds when *N(T)*>>*N(0)*, which is true experimentally.

We grew cells in excess metabolite (lysine or hypoxanthine) exponentially to time *T* when OD_600_ < 0.5 (i.e. <1.6×10^7^/ml). Supernatants were assayed for released metabolite using the rate bioassay (Methods, “Bioassays”). Since *M(T)* was below the sensitivity of detection (~0.1 μM; Fig 3-Figure Supplement 1) for both strains, we used 0.1 μM as *M(T)*, growth rate (0.47~0.48/hr for *L*^*−*^*A*^*+*^ and 0.43~0.44/hr for *A*^*−*^*L*^*+*^), and *N(T)* (1.4~1.6×10^7^/ml) to calculate the upper bound for release rate *r*.

### Genomic Analysis

High-quality genomic DNA was extracted using the QIAGEN Genomic-tip 20G kit (CAT No. 10223) or the Zymo Research YeaStar Genomic DNA Kit (CAT No. D2002). DNA fragmentation and libraries were prepared ^56^ using a Nextera DNA Sample Preparation Kit with 96-custom barcode indices ^57^ and TruSeq Dual Index Sequencing Primers. Libraries were pooled and multiplexed on a HiSeq2000 lane (Illumina) for 150 cycle pairedend reading. A custom analysis pipeline written in Perl incorporated the bwa aligner ^58^ and samtools ^59^ for alignment file generation, GATK for SNP/indel calling ^60^, and cn.MOPs for local copy-number variant calling ^61^. Finally, a custom Perl script using vcftools ^62^ was used to automate the comparison of an evolved clone with its ancestor. All called mutations were validated by visual inspection in the IGV environment ^63^.

Ploidy was calculated using custom python and R scripts. Read depth was counted for each base, and averaged within consecutive 1000-bp windows. Then, the average coverage of each 1000-bp window was normalized against the median of these values from the entire genome, and log_2_ transformed. Transformed data were plotted as box-plots for each chromosome/supercontig. All code is publically available at https://github.com/robingreen525/ShouLab_NGS_CloneSeq.

### Calculating death rate in non-limited batch culture

We grew cells to exponential phase in SD + excess supplements. While still at a low density (<10^7^ cells/ml), we measured live and dead cell densities using flow cytometry to yield *dead/live* ratio. Since the percentage of dead cells was small, we analyzed a large volume of sample via flow cytometry to ensure that at least 400 ToPro3-stained dead cells were counted so that the sampling error 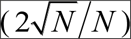 was no more than 10%. We also calculated growth rates using optical density readings for the two hours before and after flow cytometry measurement to yield net growth rate *g*. In exponentially-growing cells,

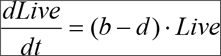

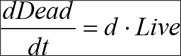

where *b* and *d* are respectively birth and death rates of cells.

Thus, 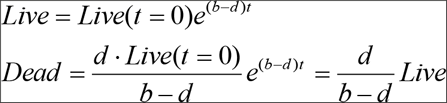, or

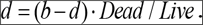

Thus, the ratio of dead to live cells is the ratio of death rate to net growth rate. The death rate of *lys2*-cells in excess lysine ranged from 10^−4^ to 10^−3^/hr. This large variability persisted despite our using the same culture master mix to grow independent cultures.

### Quantifying spatial CoSMO growth dynamics

In all experiments, *L*^*−*^*A*^*+*^ cells were grown to exponential phase in SD plus lysine, and washed free of lysine. *A*^*−*^*L*^*+*^ cells were grown to exponential phase in SD plus hypoxanthine, washed, pre-starved in SD for 24 hrs, and washed again. Pre-starvation was intended to deplete cellular hypoxanthine storage and to shorten CoSMO growth lag (Fig 1-Figure Supplement 2). We grew spatial CoSMO in two configurations: “column” versus “spotting”.

In the column setting, to prevent potential metabolite cross-contamination, we over-filled non-neighboring wells (i.e. 24 wells per deep 96-well plate) with 2×SD+2% agar, and covered the surface with a sterile glass plate to form a flat agar surface with no air bubbles. After solidification, we removed the glass plate and removed extra agar between filled wells using sterile tweezers. This results in agar depth of 3 cm. For the rest of the experiment, when not setting up or sampling, we covered the plate with a sterile lid, suspended above wells by thick toothpicks. We wrapped plates with parafilm to reduce agar drying. We mixed strains at a 1:1 ratio and filtered them through MF membrane (HAWP04700 from Millipore, Billerica, MA) to achieve 3000 cells/mm^2^ density on the filter (see ^24^ for details). We then punched 8 mm diameter disks and transferred one disk to each agarose well, resulting in ~1.5×10^5^ cells/disk. For each time point, we used tweezers (ethanol-flame sterilized between samples) to pick 2~3 disks, and suspended each in water prior to flow cytometry measurements.

In the spotting setting, in an 85 mm petri dish we poured ~25ml 2×SD + 2% agarose + a small amount of lysine (generally 0.7 μM to minimize the lag phase during CoSMO growth) to achieve an agar/agarose depth of 5mm. After solidification, we used a sterile blade to cut and remove ~2mm strips out of the agar to create six similarly sized sectors on the plate with no agarose connections between them (Fig 7-Figure Supplement 4A). We inoculated plates by spotting 15μl of strains at a 1:1 ratio onto plates to result in ~4×10^4^ cells/patch (4 mm inoculum radius). Cells were grown and sampled as in the column setting, except that we cut out the agarose patch containing cells, submerged it in water, vortexed for a few seconds, and discarded agarose.

For both setups, we used 9×10^7^ total cells as a cutoff for CoSMO growth rate calculation. We used this cut-off because exponential CoSMO growth rate was observed beyond 9×10^7^ total cells, suggesting that no other metabolites were limiting by then.

### Simulating spatial CoSMO growth

We modified our previous individual-based spatial CoSMO model ^24^ so that in each time step, metabolite consumption and release of each cell scaled linearly with cell’s biomass to reflect exponential growth. The model used parameters in Table 1. The release rate of lysine for each *A*^*−*^*L*^*+*^ cell at each time-step was linearly interpolated based on the local concentration of hypoxanthine (Table 1-Table Supplement 4). We simulated CoSMO growth in two different settings: (1) cells were initially uniformly distributed on the surface of an agar column; (2) cells were initially spotted in the middle of an agar pad according to the experimental setup. The simulation domain used for setting (1) was 500 μm × 500 μm in the lateral x and y dimensions; for setting (2), the agarose domain was 800~960 μm on each side (5 μm/grid), and the size of the inoculation spot was ¼×¼=1/16 of the agarose domain. In both (1) and (2) settings, the z dimension in simulation varied according to the experimental setup (5 mm ~3 cm). For metabolite diffusion within the community, we used either a single diffusion coefficient (D = 360 μm^2^/s; Fig 7-Code Supplement 1) or two diffusion coefficients (*D* = 360 μm^2^/s measured in agarose and *D* = 20 μm^2^/s measured in yeast community ^24^; Fig 7-Code Supplement 2). Both codes are for spotting inoculation, but the inoculation spot can be increased to cover the entire surface. Regardless of the simulation setup, we obtained a similar steady state community growth rate.

### Calculating steady state community growth rate

When CoSMO achieves the steady state growth rate, both strains will grow at the same rate as the community (*g*_*comm*_). This means that *L* and *A* concentrations do not change, and Eqs 1-4 (Fig 1) become:

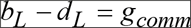

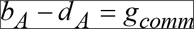

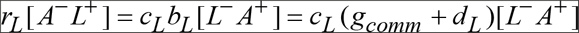

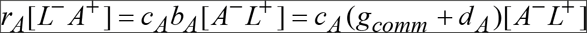

Combine the last two equations, we get

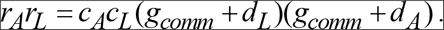

Solving this, we get

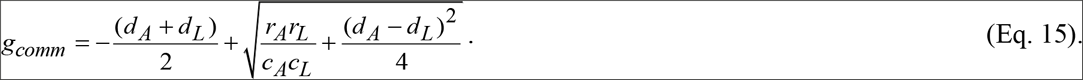

For Model iii, given our parameter values (Table 1), 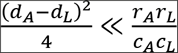. Thus, we obtain

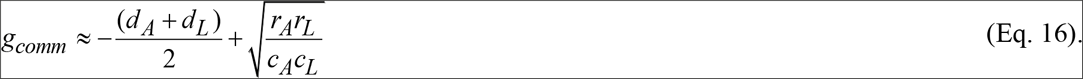

*rL*, lysine release rate of *A*^*−*^*L*^*+*^, varies with growth rate (Fig 7). When we focus on doubling times between 5.5 to 8 hrs, a range experienced by CoSMO, then we arrive at the following correlation (Fig 6B green dotted line): *r*_*L*_ =1.853-11.388*g*_*A*_ where *gA* is the net growth rate of *A*^*−*^*L*^*+*^.

Since at steady state growth rate *g*_*A*_ = *g*_*comm*_, we have

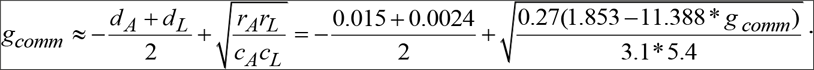

(*g*_*comma*_ + 0.0087)^2^ =(1.853-11.388\**g*_*comma*_)0.0161 = 0.0298-0.1833*g*_*comma*_. That is,

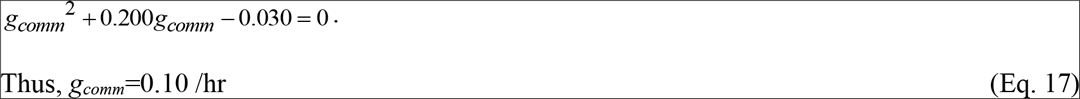

corresponding to a doubling time of 6.9 hrs.

To estimate the uncertainty in our prediction of *g*_*comm*_, we use the variance formula for error propagation. Specifically, let *f* be a function of *x*_*i*_ (*i*=1, 2,…, *n*). Then, the error of *f*, *s*_*f*_, can be expressed as

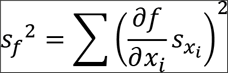

where 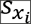. is the uncertainty of *x*_*i*_.

Thus, for each of the six parameters in Eq. 16, we divide its 95% confidence interval (Table 1) by 2 to obtain error *s*. For lysine release rate *rL*, we use the value measured in chemostats with a 7-hr doubling time which closely corresponds to CoSMO doubling time.

1. 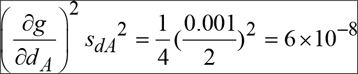
2. 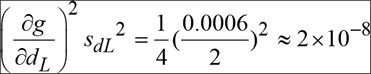
3. 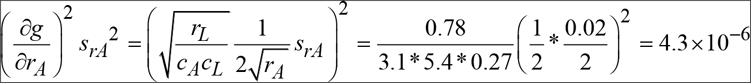
4. 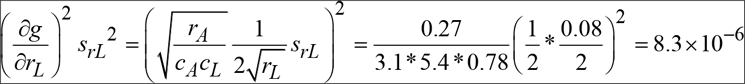
5. 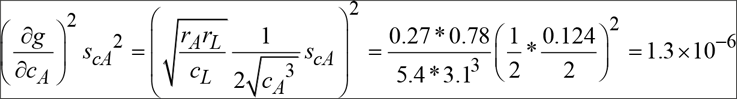
6. 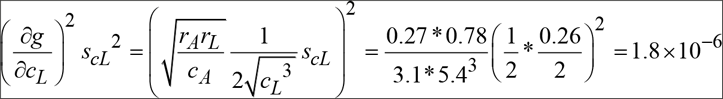

Summing all terms and taking the square root, we have an error of 0.004 for *g*_*comm*_. Thus, the 95% confidence interval is +/− 0.01.

We did not calculate the uncertainty of our spatial simulation prediction, since we did not solve the spatial model analytically. However, given that predicted community growth rates with or without diffusion are similar (Fig 7), we expect that the two predictions should share similar uncertainty.

---

***Table 1-Table Supplement 1. ^*−*^A^*+*^ consumption***

***Table 1-Table Supplement 2. A^*−*^L^*+*^ consumption***

***Table 1-Table Supplement 3. L^*−*^A^*+*^ death and release***

***Table 1-Table Supplement 4. A^*−*^L^*+*^ death and release***

**Fig 1-Figure Supplement 1.**
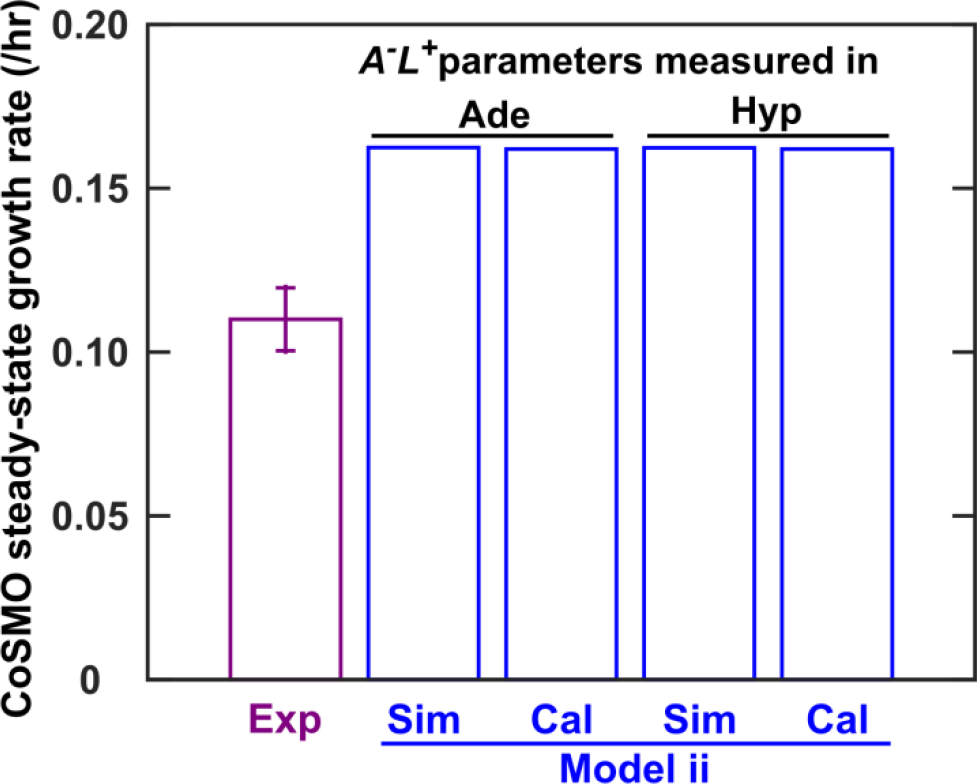
Using *A*-*L*+ phenotypes measured in batch monocultures supplemented with adenine versus hypoxanthine did not affect model performance. Fig 1-Figure Supplement 1. “Exp”: community growth rates were calculated from seven independent experiments in a well-mixed environment (from ~30 hr to 70~80 hr) and averaged, with the error bar representing two standard deviations. “Model ii”: all model parameters were derived from RM11 *L*^*−*^*A*^*+*^ and *A*^*−*^*L*^*+*^ phenotypes measured in batch mono-cultures. We predicted steady state community growth rate either via quantifying the simulated post-lag dynamics (e.g. Fig 1C “Model ii”) (“Sim”) or via an analytical formula (Eq. 15 in Methods) (“Cal”). The experimental and predicted doubling times were 6.5 hr and 4.3 hr, respectively. Experimental data and model parameters are listed in Fig 1-Table Supplement 2

**Fig 1-Figure Supplement 2.**
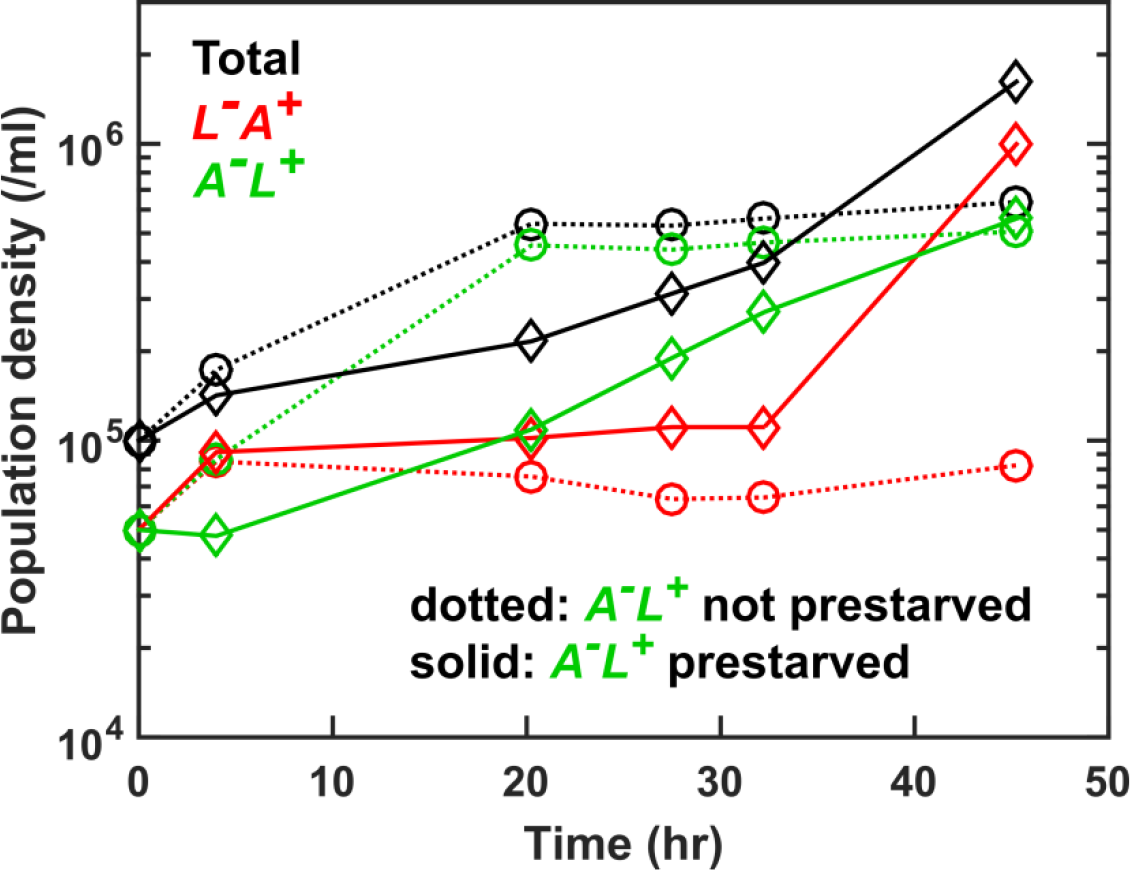
Prestarving *A*^*−*^*L*^*+*^ reduces the lag phase of community growth. Fig 1-Figure Supplement 2. Exponential *A*^*−*^*L*^*+*^ (WY1340) cells were washed free of hypoxanthine, and either prestarved for 24 hrs in SD (solid lines) or not pre-starved (dotted lines) before being mixed with exponentially-grown and washed *L*^*−*^*A*^*+*^ (WY1335) to form CoSMO. Pre-starvation of *A*^*−*^*L*^*+*^ leads to less growth lag compared to no pre-starvation.

***Fig 1-Table Supplement 1. Strain table***

***Fig 1-Table Supplement 2. Experimental measurements of well-mixed CoSMO population dynamics and initial model parameters***

***Fig 1-Code Supplement 1. Initial model for well-mixed CoSMO***

**Fig 2-Figure Supplement 1.**
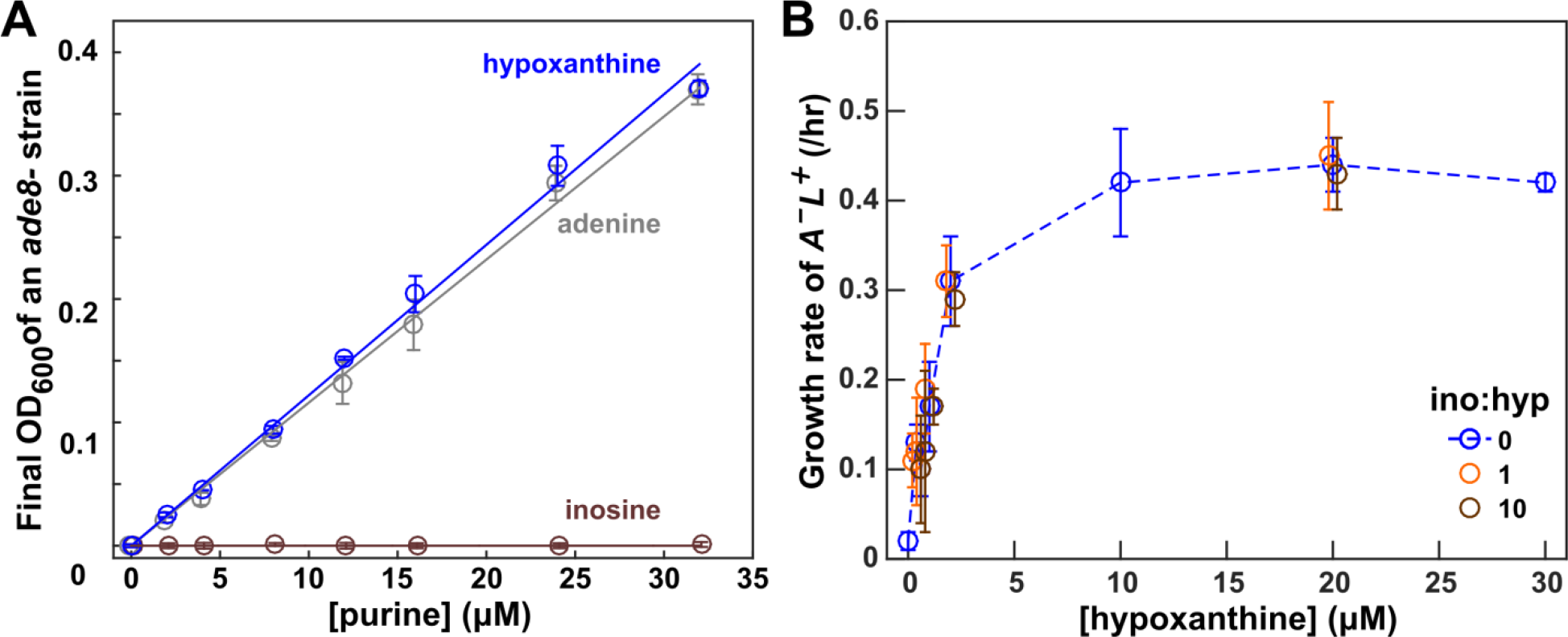
Inosine does not mediate the interaction from *L*^*−*^*A*^*+*^ to *A*^*−*^*L*^*+*^. Fig 2-Figure Supplement 1: (**A**) Hypoxanthine but not inosine is consumed by *A*^*−*^*L*^*+*^. The final turbidity of an *ade8-* (WY1340) tester strain increases with increasing concentrations of hypoxanthine (blue) and adenine (grey) but not inosine (brown). The slopes of the blue and grey lines are similar, suggesting that a similar amount of hypoxanthine and adenine are consumed to produce one new *A*^*−*^*L*^*+*^ cell. (**B**) Stimulation of *A*^*−*^*L*^*+*^ (WY1340) growth rate by hypoxanthine (blue) is not affected by the presence of inosine at 1x (orange) or 10x (brown) concentration.

**Fig 2-Figure Supplement 2.**
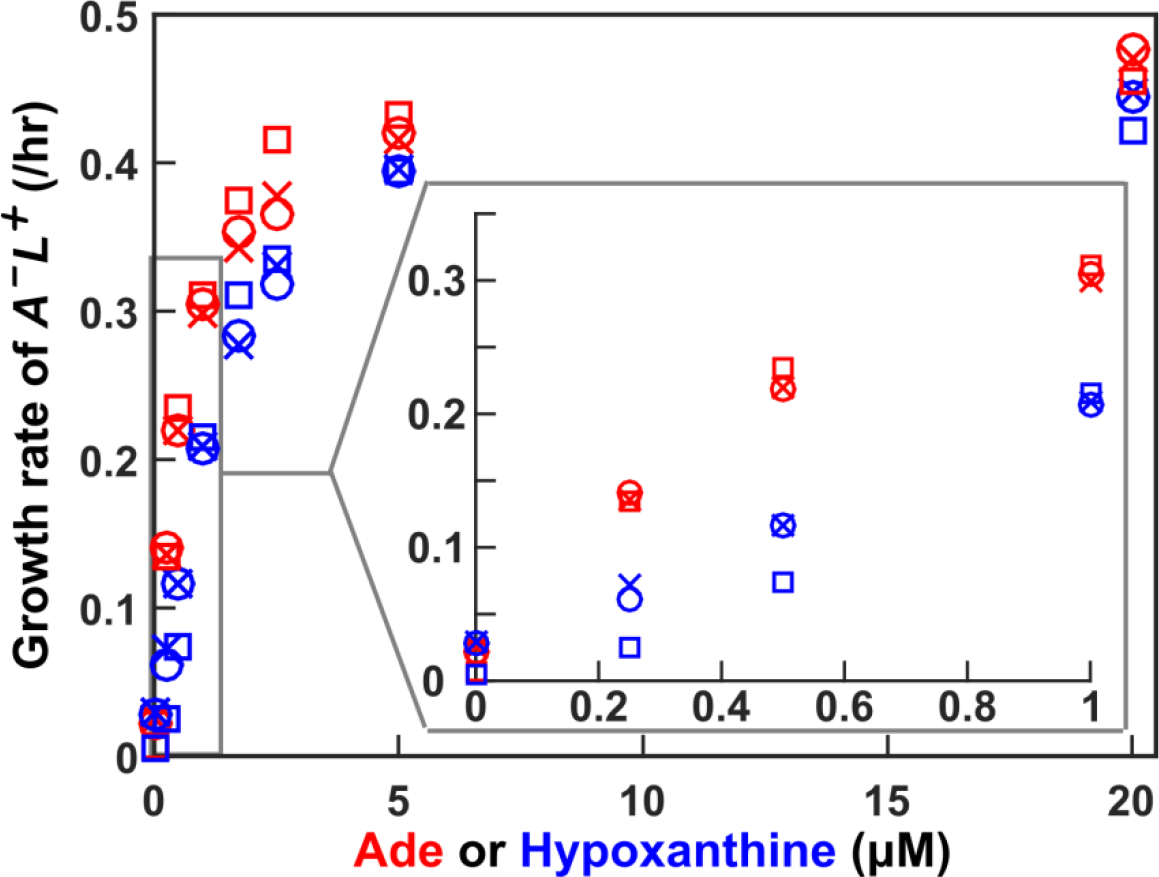
Hypoxanthine and adenine lead to quantitatively different growth phenotypes in *A*^*−*^*L*^*+*^. *A*^*−*^*L*^*+*^ cells grow faster when fed with adenine (red) than when fed with hypoxanthine (blue) when metabolite concentration is low (inset). *A*^*−*^*L*^*+*^ (WY1340) cells pre-grown in SD + adenine or SD + hypoxanthine were washed into SD and pre-starved for 24 hrs to deplete intracellular storage. Subsequently, adenine or hypoxanthine was supplemented at various concentrations, and the net growth rate was measured via fluorescence microscopy (Methods, “Microscopy quantification of growth parameters”). Red circles and squares: pre-grown in adenine, and incubated in adenine; red crosses: pre-grown in hypoxanthine, and incubated in adenine; blue circles and squares: pre-grown in hypoxanthine, and incubated in hypoxanthine; blue crosses: pregrown in adenine, and incubated in hypoxanthine. Pre-growth in cognate metabolite versus non-cognate metabolite does not make a difference (e.g. compare red circles with red crosses, and blue circles with blue crosses, all of which were measured in the same experiment).

**Fig 2-Figure Supplement 3.**
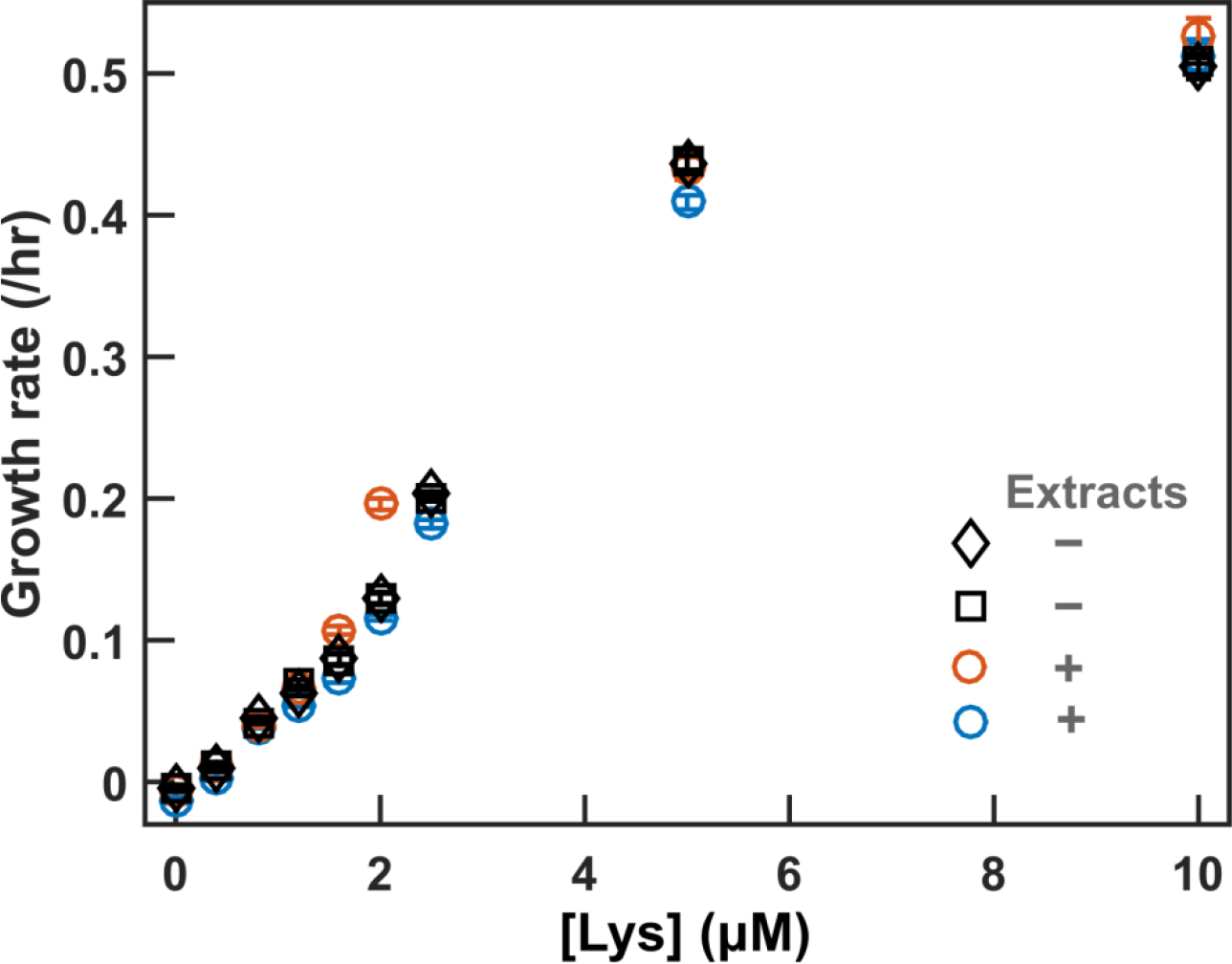
Cell extracts do not interfere with bioassays. Exponential *L*^*−*^*A*^*+*^ (WY1335) cells were starved in SD for 4 hrs to deplete intracellular storage of lysine. 2.5 ml of starved culture at OD~0.2 were used to extract intracellular metabolites (“Extraction of intracellular metabolites” in Methods). The dried pellet was re-suspended in ~1 ml H2O. In a separate experiment, exponential *L*^*−*^*A*^*+*^ were washed and pre-starved in SD for 4 hours. We then quantified the growth rates of *L*^*−*^*A*^*+*^ in SD supplemented with 1/3 volume of extracts (orange and blue) or water (black) as well as various concentrations of lysine (“Microscopy quantification of growth phenotypes” in Methods). The inclusion of extracts did not affect growth rates.

**Fig 3-Figure Supplement 1.**
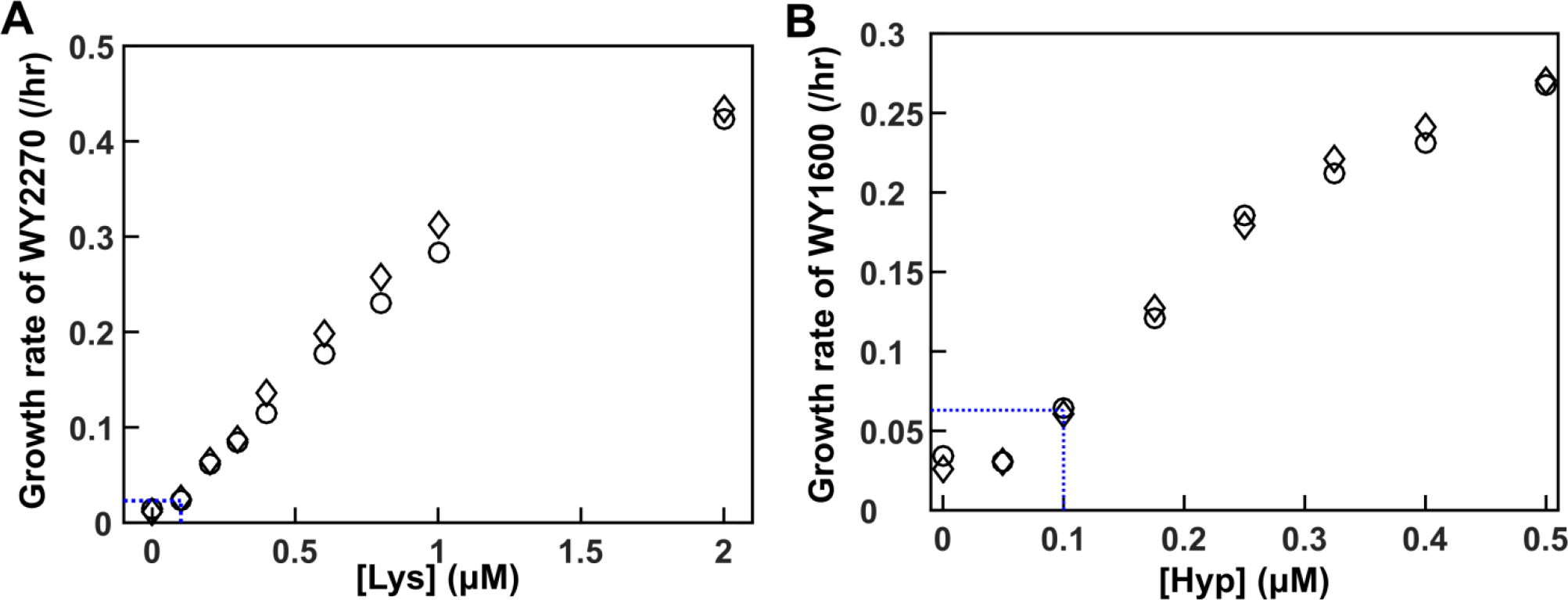
Using evolved clones to measure low concentrations of metabolites. (**A**) WY2270, an evolved *L*^*−*^*A*^*+*^ clone with significantly improved affinity for lysine, could detect sub-1 μM Lys. (**B**) WY1600, an evolved *A*^*−*^*L*^*+*^ clone with significantly improved affinity for hypoxanthine, could detect sub-1 μM hypoxanthine. Vertical dotted blue lines mark detection limits. Circles and diamonds mark two independent replicates.

**Fig 3-Figure Supplement 2.**
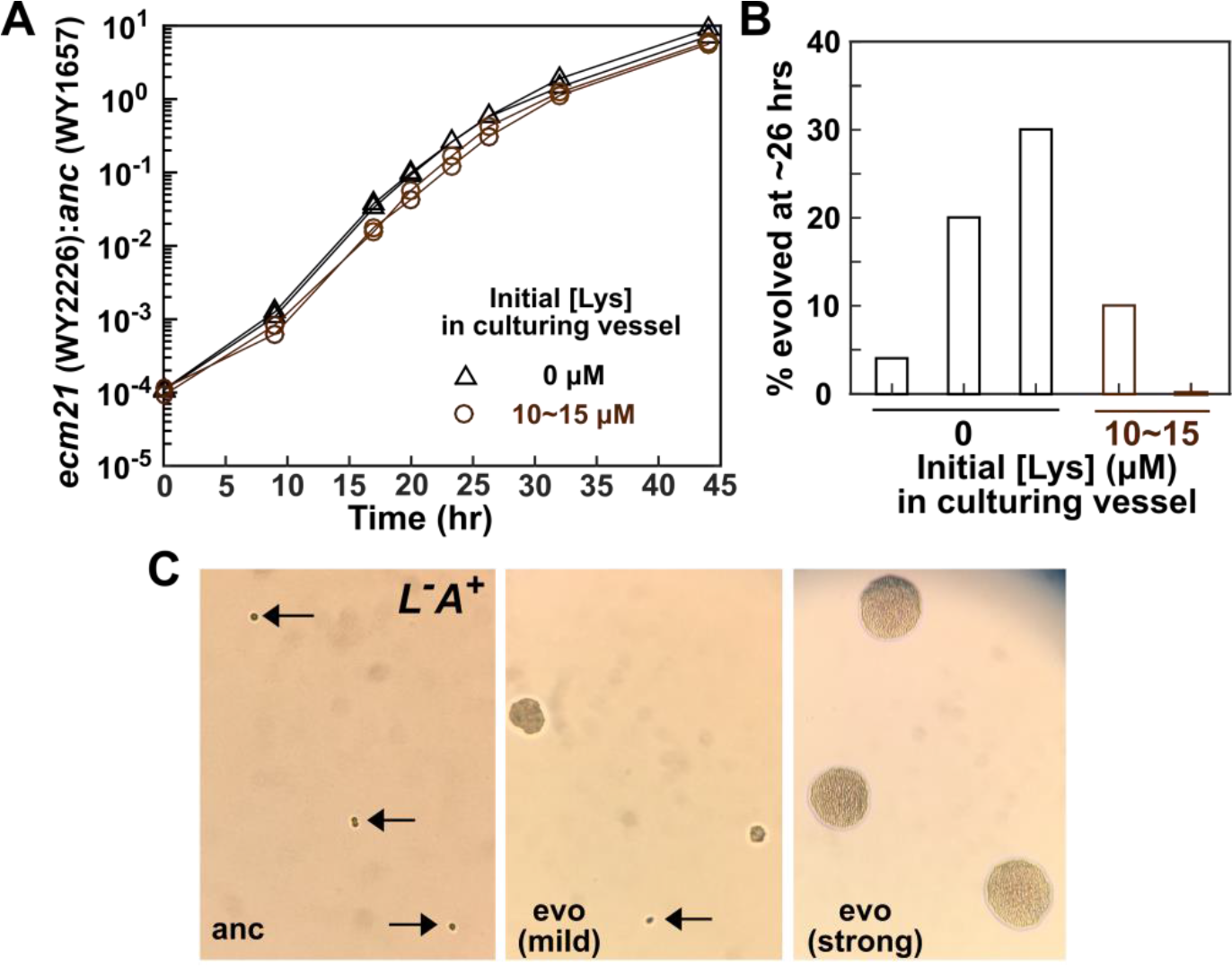
Characterization of evolved *L*^*−*^*A*^*+*^ clones. Whole-genome sequencing revealed that evolved *L*^*−*^*A*^*+*^ clones harbor mutations in genes such as *RSP5* (an E3 ubiquitin ligase) and *ECM21* (an arrestin-like adaptor for Rsp5) ^65^. In a stressful environment, wild-type Ecm21 and Rsp5 proteins target cell-surface permeases (including the high-affinity lysine permease Lyp1) for ubiquitination ^64^. Ubiquitinated permeases are then endocytosed and degraded in the vacuole ^64^. The resulting amino acids are then transported to the cytoplasm for protein synthesis to help cells cope with stress ^67^. In evolved cells with mutant *ecm21* or *rsp5*, lysine permease is stabilized. (**A**) Evolved *L*^*−*^*A*^*+*^ grows faster than the ancestor in lysine-limited chemostats. *L*^*−*^*A*^*+*^ with or without an *ecm21* deletion (WY2226 and WY1657, respectively) expressing different fluorescent proteins were competed in 8-hr doubling time chemostats. The initial lysine concentrations in culturing vessels was 0 (black triangles) or 10~15μM (brown circles). In all four chemostats, *ecm21* overtook ancestor. The fitness diffxserence between the two strains can be estimated: Let *E*(*t*) and *A*(*t*) be population densities of *ecm21* and ancestor at time *t*, respectively, and *rE* and *rA* be the growth rates of the two strains. Then, 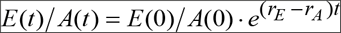, and we have ln (*E*(*t*)/*A*(*t*))=ln(*E*(0))/*A*(0))+(*r*_*E*_ − *r*_*A*_)*t*. We quantified (*r*_*E*_ − *r*_*A*_), the fitness advantage of *ecm21* over ancestor, as 0.31/hr (computed up to 32 hrs), compared to ancestor growth rate of 0.087/hr (8-hr doubling). Thus, *ecm21* grows ~3.6x faster than the ancestor. This fitness advantage is qualitatively consistent with what we observed in chemostats initiated with pure ancestor, since evolved clones increased from ~4% to ~40% within 5.7 hrs (from 26.3 to 32 hrs in Fig 3C), translating to a 49/hr fitness difference. We infer that evolved clones are initially present at a frequency on the order of ~0.04/exp(0.4/hr*26.3hr)=10^−6^. This is in-line with the phenotypic mutation rate of 0.5~30 × 10^−7^ per cell per generation ^68^ in the following sense. Since the inoculum population size is on the order of 4×10^6^ cells/ml × 19 ml= 8 ×10^7^ cells, we expect 2×8×10^7^*(0.5~30 × 10^−7^)=8~480 preexisting evolved cells (the coefficient of 2 results from (1 + 2 + 2^2^ + …+ 2^*n*^)/2^*n*^~2 where the numerator represents total mutation opportunity in a culture starting from a single cell, and the denominator represents the final size of chemostat inoculum). Thus, the early evolutionary dynamics in chemostats can be explained by pre-existing mutants outcompeting ancestral cells. (**B**) Percent evolved clones in ~26-hr samples from chemostats inoculated with ancestral *L*^*−*^*A*^*+*^ (WY1335). (**C**) A visual assay that distinguishes ancestral versus evolved *L*^*−*^*A*^*+*^ clones. *L*^*−*^*A*^*+*^ cells from an ancestral clone and two evolved clones were plated on SD plates supplemented with 1.5 μM lysine. Ancestral cells (WY1335, left) failed to divide (arrows). Cells from a mildly-adapted evolved clone (harboring duplication of Chromosome 14, center) showed heterogeneous phenotypes: some cells remained undivided (arrow), while other cells formed microcolonies of various sizes. Cells from a strongly-adapted evolved clone (harboring an *ecm21* mutation, right) formed microcolonies of a uniform and large size. These images were taken using a cell phone camera and thus do not have a scale bar. For reference, an average yeast cell has a diameter of ~5 μm.

**Fig 3-Figure Supplement 3.**
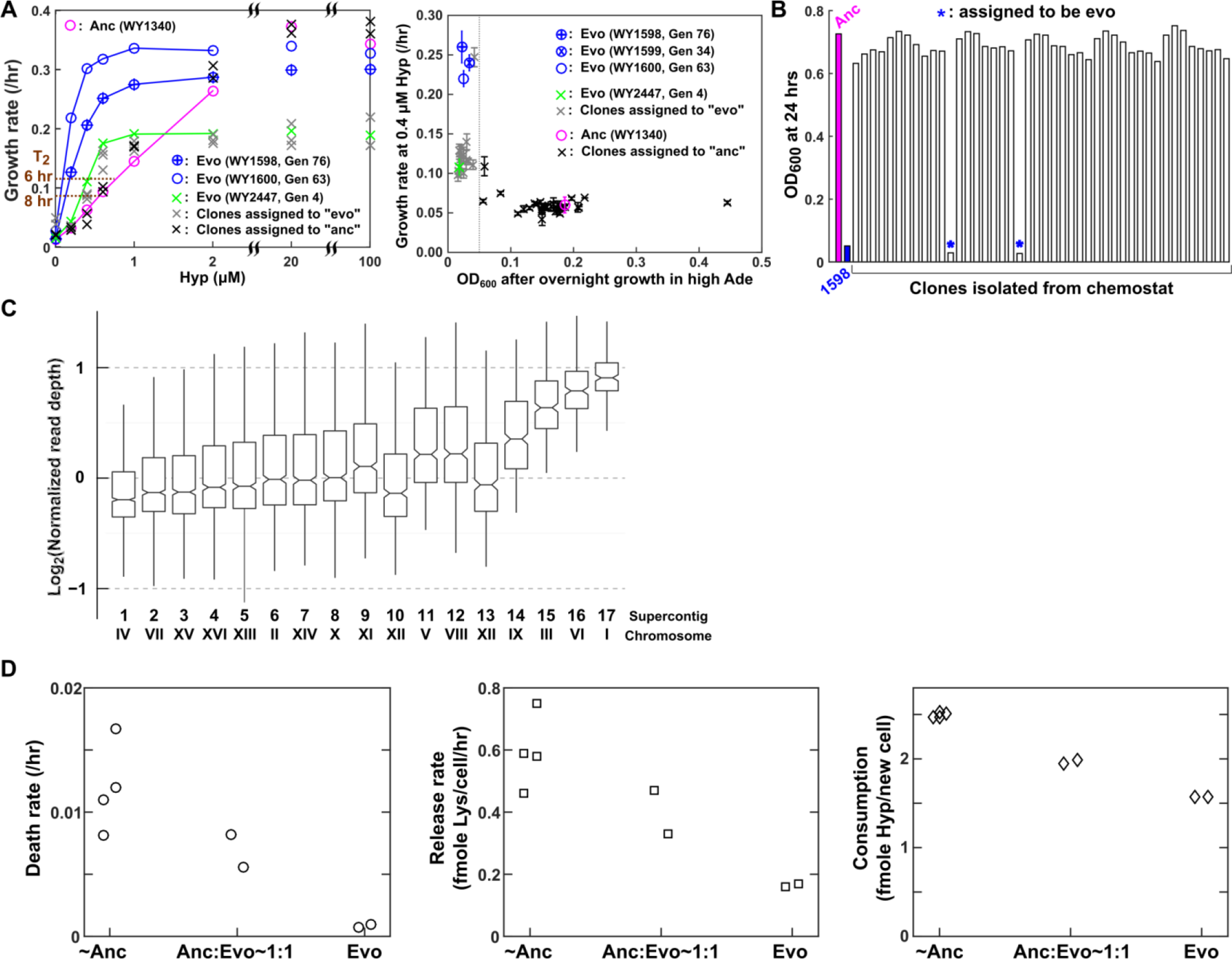
Characterization of evolved *A*^*−*^*L*^*+*^ clones. (**A**) A fitness tradeoff in *A*^*−*^*L*^*+*^. (**Left**) Growth rates of ancestral (magenta) and evolved (blue and green) *A*^*−*^*L*^*+*^ clones (pre-starved overnight) in various concentrations of hypoxanthine (Methods, “Microscopy quantification of growth parameters”) were plotted. Brown dotted lines mark 6-hr and 8-hr doubling time, a range experienced by CoSMO. In CoSMO, hypoxanthine concentrations were low (~1 μM). Evolved clones grew faster than the ancestor under low hypoxanthine concentrations, but grew slower than the ancestor under high hypoxanthine concentrations (e.g. 20~100 μM). Clones marked by crosses were isolated from Generation 4 (hour 31) of chemostat culturing. (**Right**) A negative correlation between growth rate at low hypoxanthine (~0.4 μM) versus turbidity in high adenine after overnight growth. Error bars on growth rate indicate 95% confidence interval on slope (rate) estimation. Grey line indicates the threshold by which we differentiated evolved clones (left of grey line) from ancestral clones (right of grey line), according to the growth rate assay in the left panel. In both panels, grey crosses represent Gen 4 clones assigned to be evolved, while black crosses represent Gen 4 clones assigned to be ancestral. (**B**) A high-throughput assay that distinguishes ancestral from evolved *A*^*−*^*L*^*+*^ clones. We used turbidity after overnight growth in high Ade (108 μM) to classify *A*^*−*^*L*^*+*^ clones as ancestral (no blue stars) or evolved (blue stars). The ancestral clone (WY1340) and an evolved clone (WY1598) are shown as controls. (**C**) Aneuploidy in the evolved clone WY2447. Whole-genome sequencing revealed that in addition to a synonymous nucleotide change, two nucleotide changes in non-coding regions, and a point mutation from Cys102 to Ser in OAR1 (Fig 3-Table Supplement 1), Chromosomes I, III, and VI are likely duplicated. For Chromosome III, read depth was not fully twice that of other chromosomes, which could be caused by cells losing the extra copy of Chromosome III during culturing prior to sequencing. (**D**) Evolved *A*^*−*^*L*^*+*^ cells have a lower death rate, a lower lysine release rate, and lower hypoxanthine consumption per birth compared to the ancestor. Phenotypes of ancestral *A*^*−*^*L*^*+*^ (WY1340 with preexisting WY2447-like mutants), 1:1 anc:evo (WY1340:WY2447), or evolved *A*^*−*^*L*^*+*^ (WY2447) were measured in 8-hr chemostats (Methods, “Quantifying phenotypes in chemostats”).

**Fig 3-Figure Supplement 4.**
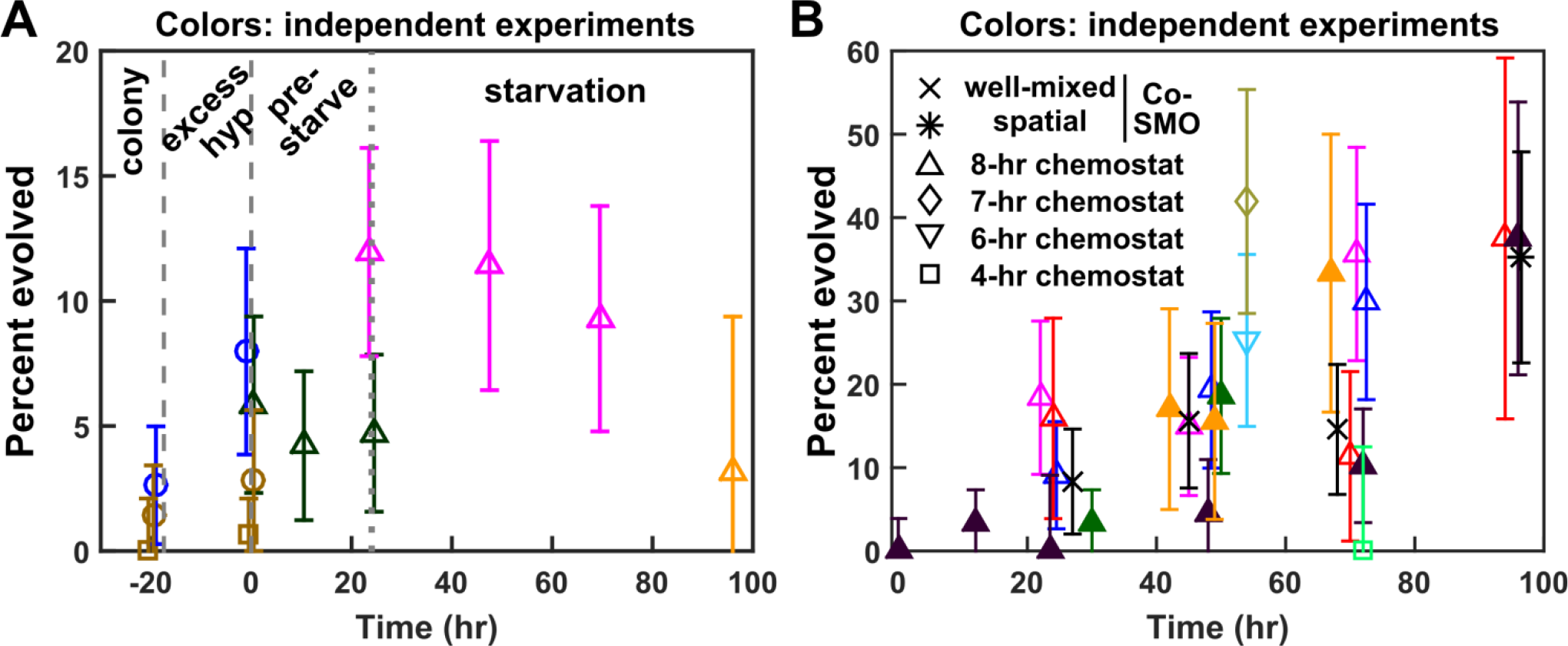
High levels of evolved clones in *A*^*−*^*L*^*+*^. (**A**) High levels of evolved *A*^*−*^*L*^*+*^ clones prior to a starvation experiment (left of dotted line) and during a starvation experiment (right of dotted line). Different colors represent experiments on different days. In experiments that terminated at 0 hr (circles and squares), an entire colony grown on rich YPD (circles) or minimal SD plus excess (100 μM) hypoxanthine (square) was resuspended in SD, and a fraction was used to inoculate SD plus excess hypoxanthine to grow exponential cultures. Otherwise (triangles), a fraction of YPD-grown colony was used to inoculate SD plus excess hypoxanthine to grow exponential cultures, and at time zero, the culture was washed free of supplements and starved of hypoxanthine. (**B**) Similar percentages of evolved *A*^*−*^*L*^*+*^ clones in chemostats and in CoSMO. For chemostat experiments, exponentially-growing cells washed free of supplements were pre-starved (unfilled symbols) or not pre-starved (filled symbols), and inoculated into chemstats (time zero). For CoSMO experiments, *A*^*−*^*L*^*+*^ cells were prestarved. In all experiments, we used the assay in Fig 3-Figure Supplement 3B to distinguish ancestral and evolved clones (Methods, “Detecting evolved clones”). If we sampled *n*_*tot*_ cells, and *n*_*evo*_ cells were evolved, then the fraction evolved was estimated to be *n*_*evo*_/ *n*_*tot*_, with error bar indicating 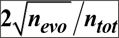 (assuming that the random variable *n*_*evo*_ followed a Poisson distribution). If zero assayed colonies were evolved, we found the maximal frequency of evolved clones such that the error bar of 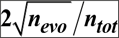 still covered zero, and used that error bar. For example, if 0 out of 88 was evolved (0%), and since 3 out of 88 had a frequency of 3.4% with an error bar of 3.9% which covered zero, we added an error bar of 3.9% above the 0% data point.

**Fig 3-Figure Supplement 5.**
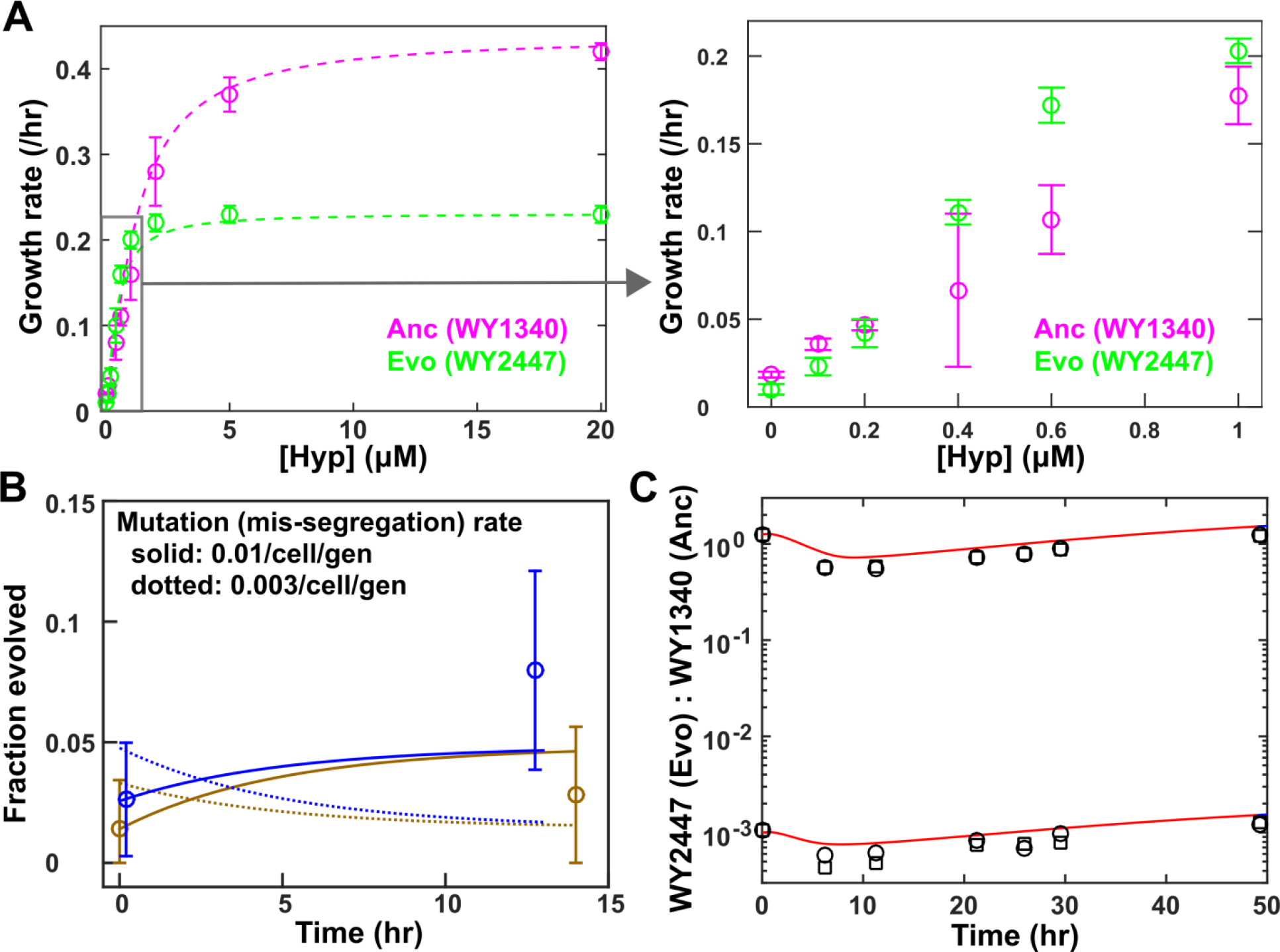
Evolutionary dynamics of *A*^*−*^*L*^*+*^ is consistent with high chromosomal mis-segregation rate. (**A**) Growth rates of ancestral (WY1340) and evolved (WY2447) *A*^*−*^*L*^*+*^ cells in various concentrations of hypoxanthine (24-hr pre-starvation; Methods, “Microscopy quantification of growth phenotypes”). Three experiments were averaged, and error bars indicate two standard deviations. (**B**) The evolutionary dynamics of *A*^*−*^*L*^*+*^ in excess hypoxanthine could be explained if we assumed that chromosome mis-segregation generated WY2447-like mutants at a rate of 0.01/cell/gen (solid lines). As a comparison, predictions from a mutation rate of 0.003/cell/gen (dotted lines) were also plotted. Brown and blue circles (measured in two different experiments) are identical to the corresponding ones in Fig 3-Figure Supplement 4A. Specifically, from the inoculum size and the final population size, we calculated the number of generations, which we then multiplied with the doubling time in SD with excess hypoxanthine to obtain the duration of exponential phase. We then inferred the lag phase to be ~6 hrs, and assumed that the fraction of evolved cells at time zero (the beginning of exponential phase) was similar to that at the time of inoculation. Our model (Fig 3-Code Supplement 1) considered the fitness advantage of ancestor over mutant in excess hypoxanthine (**A**), as well as the conversion from ancestor to mutant. Data at 0 hr were slightly jittered to aid visualization. (**C**) We competed WY2447 (expressing citrine-fluorescent protein) and WY1340 (expressing green-fluorescent protein) in 8-hr chemostats from two starting ratios, and measured strain ratios over time using flow cytometry (black circles). Using a mathematical model (Fig 3-Code Supplement 2) where growth parameters were measured experimentally (**A**) and where the ancestor converted to WY2447-like mutants at a rate of 0.01/cell/gen, we obtained a qualitative matching between model and experiments. In both models (**B, C**), death rate and hypoxanthine consumption per birth were from 8-hr chemostat measurements (Methods, “Quantifying phenotypes in chemostats”).

**Fig 3-Figure Supplement 6.**
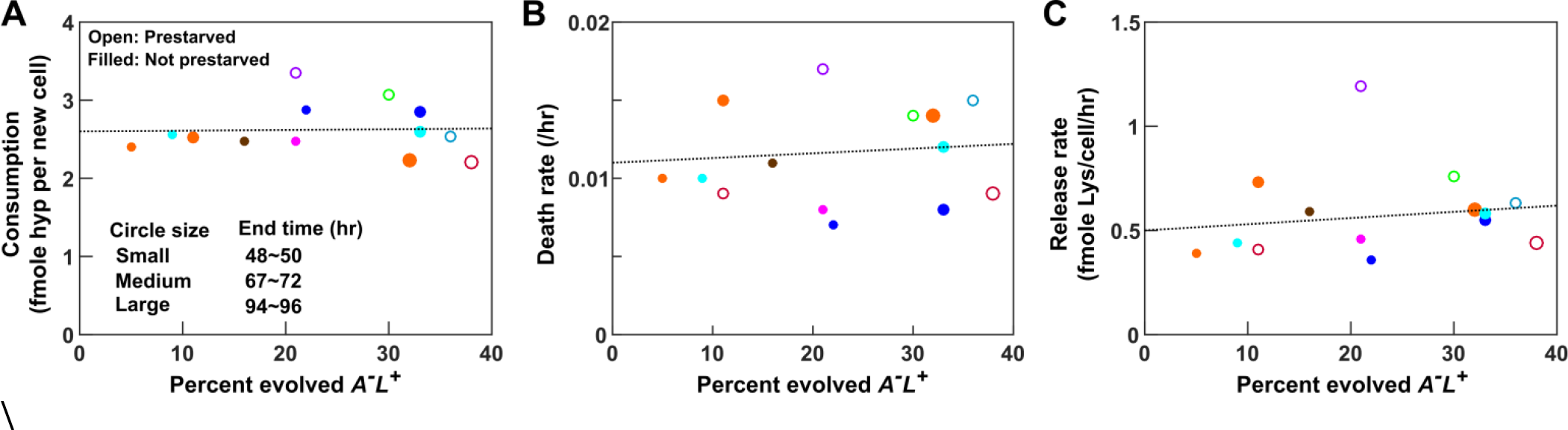
Measured *A*^*−*^*L*^*+*^ phenotypes are not significantly impacted by percent evolved clones during quantification. *A*^*−*^*L*^*+*^ (WY1340), either prestarved for 24 hrs (open circles) or not prestarved (filled circles), were cultured in 8-hr chemostats (different colors representing independent chemostat experiments). Hypoxanthine consumption per birth (**A**), death rate (**B**), lysine release rate (**C**) were quantified (Methods “Quantifying phenotypes in chemostats”) using dynamics up to 48~50 hrs (small-size circles), 67~72 hrs (medium-size circles), or 94~96 hrs (large-size circles). Percent of evolved clones was quantified at the end of each measurement. Despite phenotypic differences between ancestral and evolved *A*^*−*^*L*^*+*^ (Fig 3-Figure Supplement 3D), measured phenotypes did not show significant correlation with %evolved (slope +/− SEM being 0.1+/−0.8 (**A**), 0.003+/−0.008 (**B**), and 0.3+/−0.6 (**C**) - none significantly different from zero). This lack of correlation is presumably due to the relatively large measurement errors and the relatively narrow spread in % evolved. Take consumption as an example. Suppose that ancestral and evolved *A*^*−*^*L*^*+*^ consumed hypoxanthine at 2.5 fmole/birth and 1.5 fmole/birth, respectively (Fig 3-Figure Supplement 3D). At 10% mutants, consumption would be 2.5*0.9+1.5*0.1=2.4 fmole/birth. At 30% mutants, consumption would be 2.5*0.7+1.5*0.3=2.2 fmole/cell. This 10% difference is smaller than the measurement error. For example, at ~33% evolved *A*^*−*^*L*^*+*^ (filled dots in A), consumption varied from 2.2 to 2.8 fmole/birth.

**Fig 3-Figure Supplement 7.**
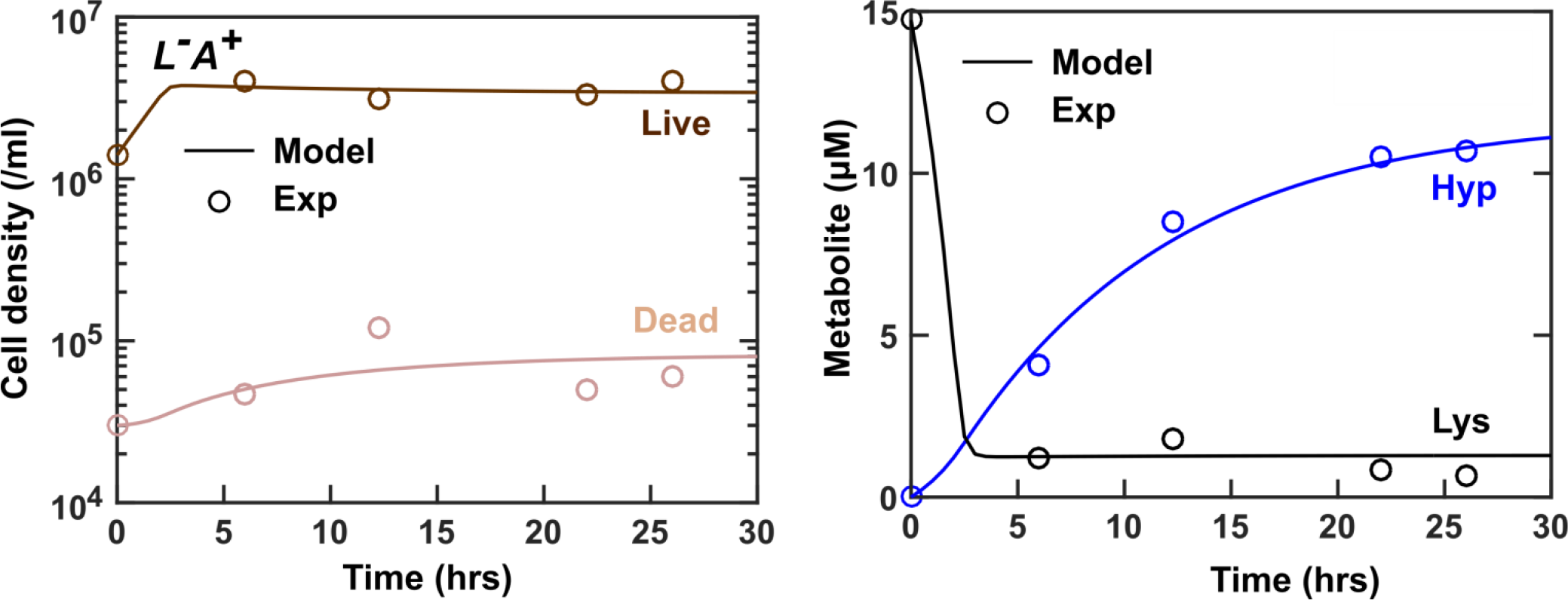
Parameters measured from *L*^*−*^*A*^*+*^ chemostats recapitulate chemostat dynamics. *L*^*−*^*A*^*+*^ cells in SD+ 15 μM lysine were inoculated into a chemostat culturing vessel (19 ml). SD + 20 μM lysine in the reservoir was pumped into the culturing vessel to achieve an 8-hr doubling time (i.e. 19 ml *ln(2)/8/hr = 1.646 ml/hr). *L*^*−*^*A*^*+*^ phenotypes in Table 1 (except for release rate of 0.30 fmole hypoxanthine/cell/hr and death rate of 0.0021/hr measured in this particular experiment) were used to simulate chemostat dynamics (Fig 3-Code Supplement 3). Simulations (lines) and experiments (circles) are in good agreement.

***Fig 3-Table Supplement 1. Mutations in WY2447***

***Fig 3-Code Supplement 1. Modeling A L+ evolution in excess hypoxanthine***

***Fig 3-Code Supplement 2. Modeling competition between evolved and ancestral A^*−*^L^*+*^ strains in hypoxanthine-limited chemostats***

***Fig 3-Code Supplement 3. Modeling chemostat dynamics of L^*−*^A^*+*^***

**Fig 4-Figure Supplement 1.**
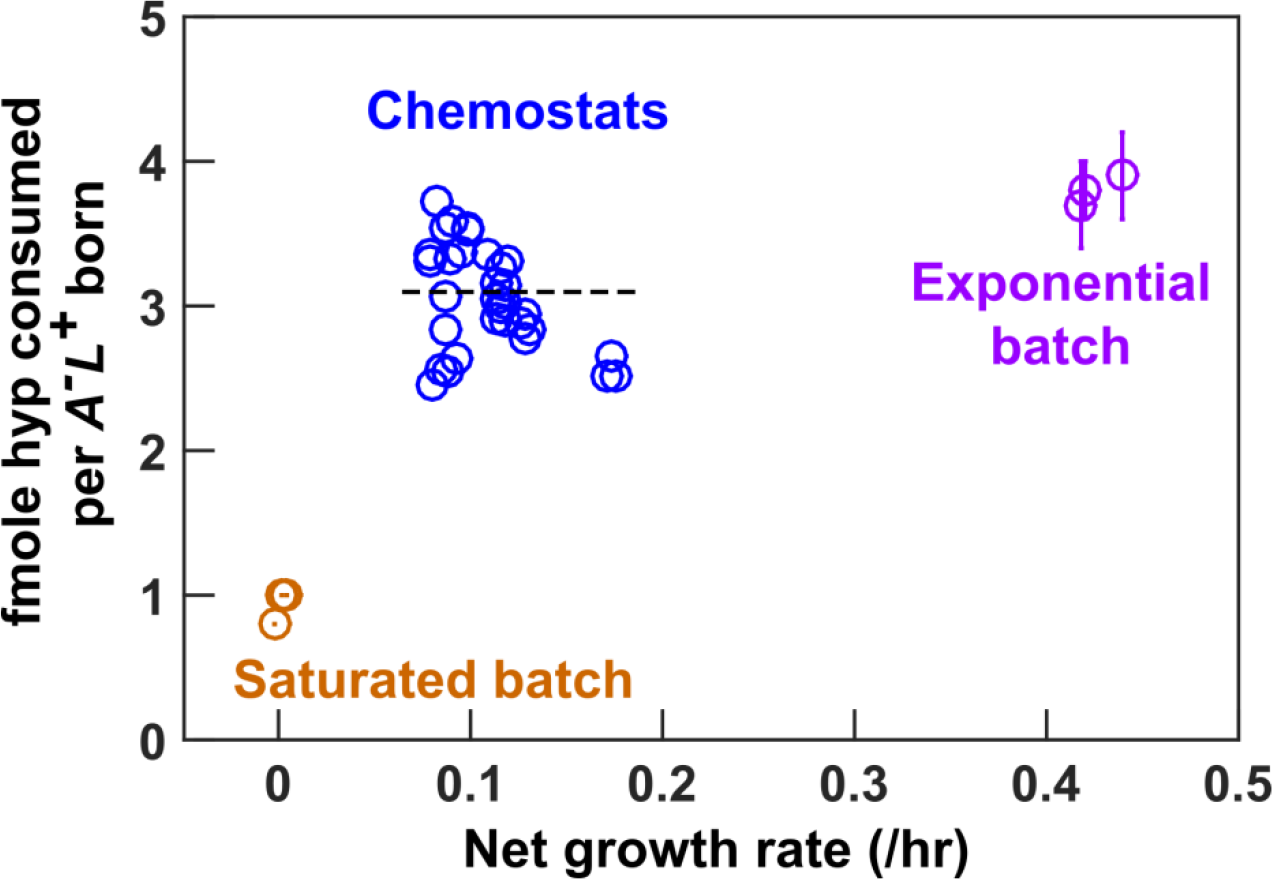
Purine consumption by *A*^*−*^L^*+*^ is relatively constant during purine-limitation. For hypoxanthine-limited chemostat measurements, data were jittered slightly along the x-axis to facilitate visualization. Consumption was measured over a similar time window as that of CoSMO growth rate to ensure similar evolutionary effects. For exponential and saturation consumption of adenine (which is similar to hypoxanthine, see Fig 2-Figure Supplement 1A), error bars mark 2 standard error of mean for slope estimation. The black dashed line marks the average hypoxanthine consumption per *A*^*−*^*L*^*+*^ birth in chemostats (Table 1; Table 1-Table Supplement 2).

**Fig 5-Figure Supplement 1.**
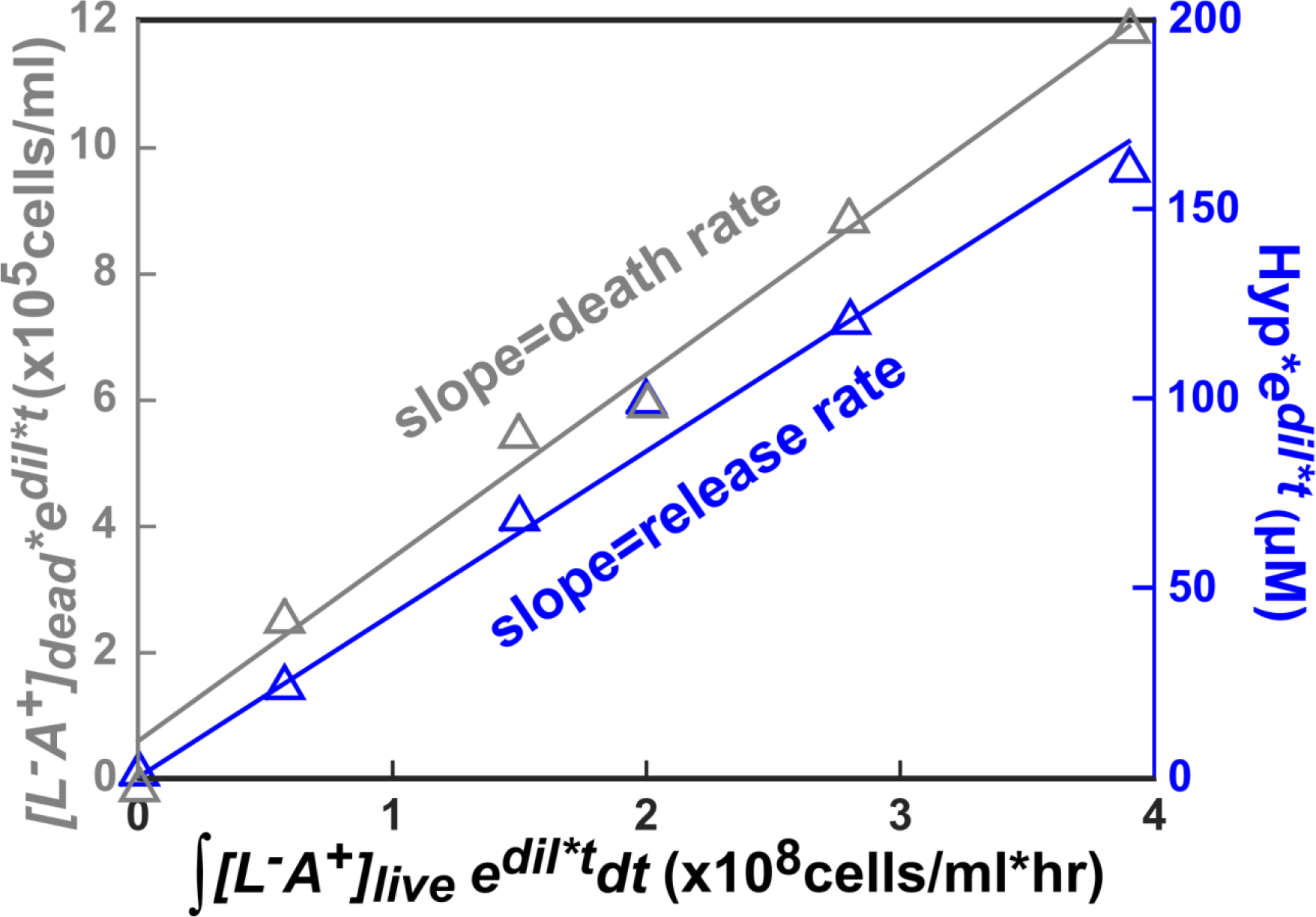
Regression analysis reveals death and release rates in a chemostat. Using regression to measure death rate (grey) and hypoxanthine release rate (blue) for the triangle-marked chemostat experiment from Fig 3. For an explanation, see “Quantifying phenotypes in chemostats” in Methods. Densities of fluorescent live cells and non-fluorescent/ToPro3-positive dead cells were measured via flow cytometry (Methods, “Flow cytometry”). Hypoxanthine was quantified using the yield bioassay (Methods, “Bioassays”).

**Fig 6-Figure Supplement 1.**
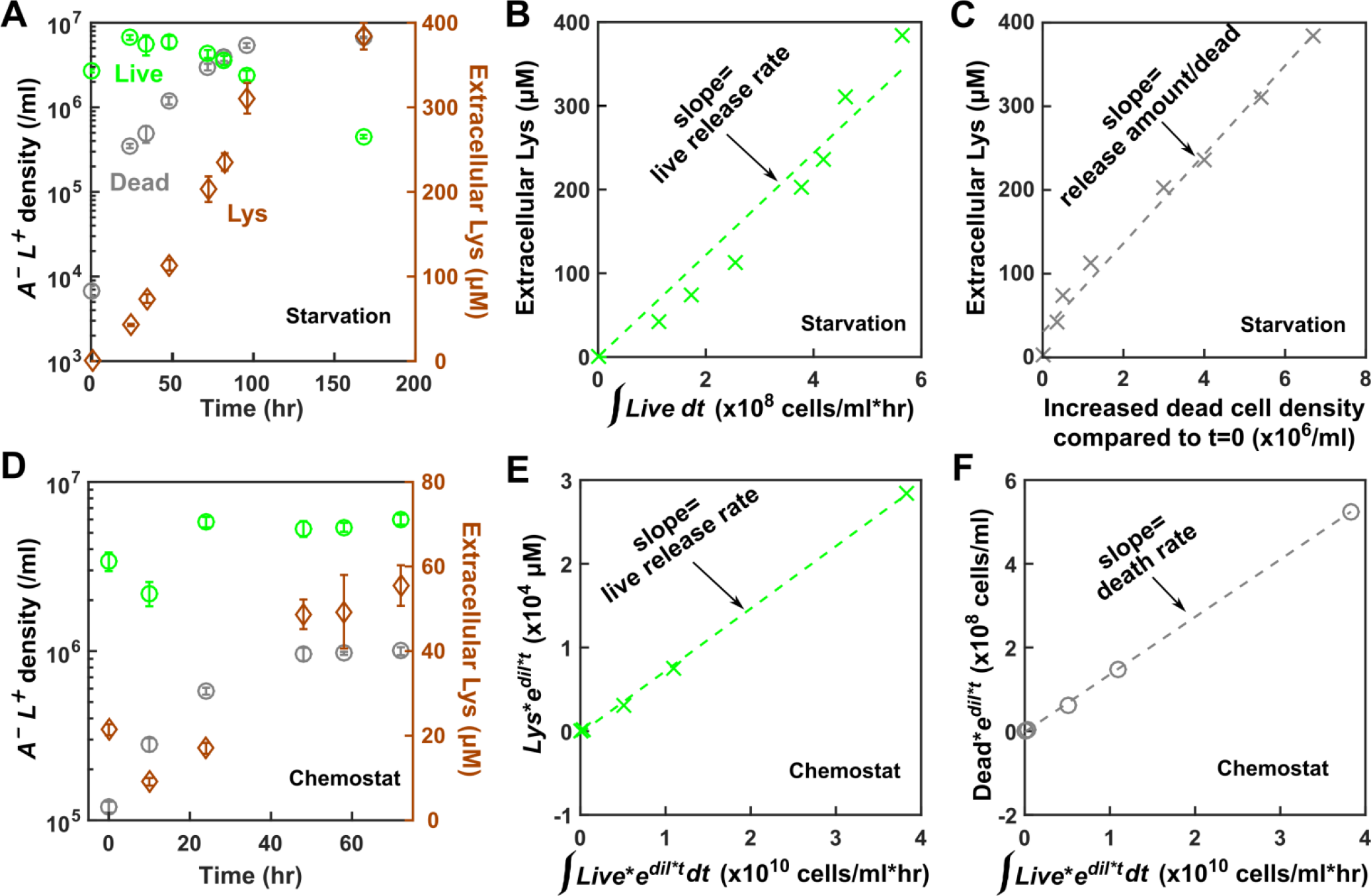
Quantifying release and death rates of *A*^*−*^L^*+*^. *A*^*−*^*L*^*+*^ cells grown to exponential phase in SD + excess hypoxanthine were washed and diluted into SD. (**A-C**) “Starvation”: At time zero, cells were inoculated into SD. From population and lysine dynamics (**A**), live release model (**B**) and dead release model (**C**) yielded a similar fit to the data. Thus from regression alone, we could not distinguish live from dead release. (**D-F**) “Chemostat”: Cells were pre-starved in SD for 24 hrs and then transferred to a hypoxanthine-limited chemostat (doubling time 8 hrs) at time zero. From population and lysine dynamics (**D**), lysine release rate by live cells (**E**) and death rate (**F**) can be calculated from slopes of respective regressions (Methods, “Quantifying phenotypes in chemostats”). Note that lysine release rate during starvation (**B**) remained relatively constant during the initial 90 hrs (the time window we later used to measure CoSMO growth rate; also see Fig 6-Figure Supplement 4B). In chemostat upon reaching the steady state, the release rate also remained relatively constant (the last three time points in **D, E**).

**Fig 6-Figure Supplement 2.**
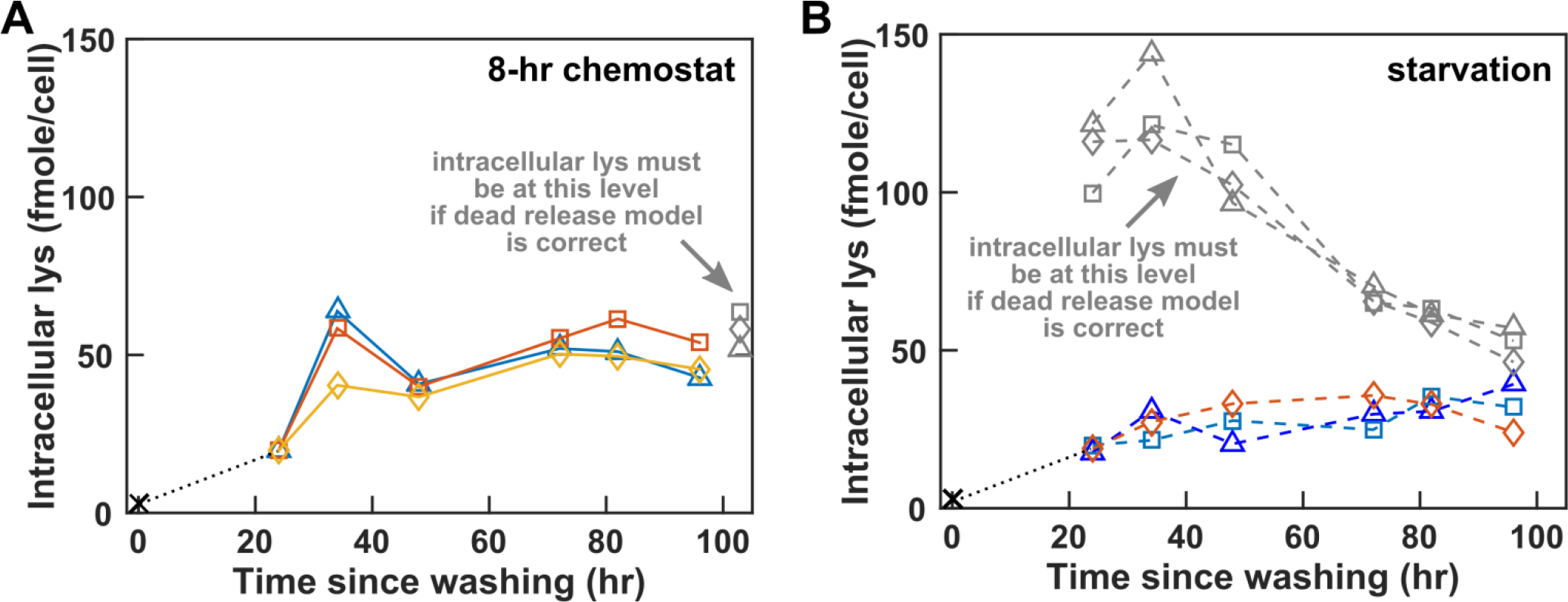
*A*^*−*^*L*^*+*^ intracellular lysine content varies with time and environment. *A*^*−*^*L*^*+*^ cells (WY1340) grown in SD+ excess hypoxanthine to exponential phase were washed and diluted into SD at time zero. Cells were either starved further (**B**) or inoculated into hypoxanthine-limited chemostats after 24 hrs of pre-starvation (**A**). At various times, cells were harvested and intracellular lysine was extracted and measured via yield bioassay. Different colors (except grey) represent different replicates and are identical to those in Fig 6A. Grey symbols in (**A**) represent the intracellular lysine content required to satisfy the dead release model at the steady state (calculated from the last three time points of Fig 6-Figure Supplement 1D). Grey symbols and dashed lines in (**B**) represent the intracellular lysine content required to satisfy the dead release model during starvation. Since live or dead release could explain chemostat results (**A**) but dead release could not explain starvation results (**B**), we made the most parsimonious assumption of live release.

**Fig 6-Figure Supplement 3.**
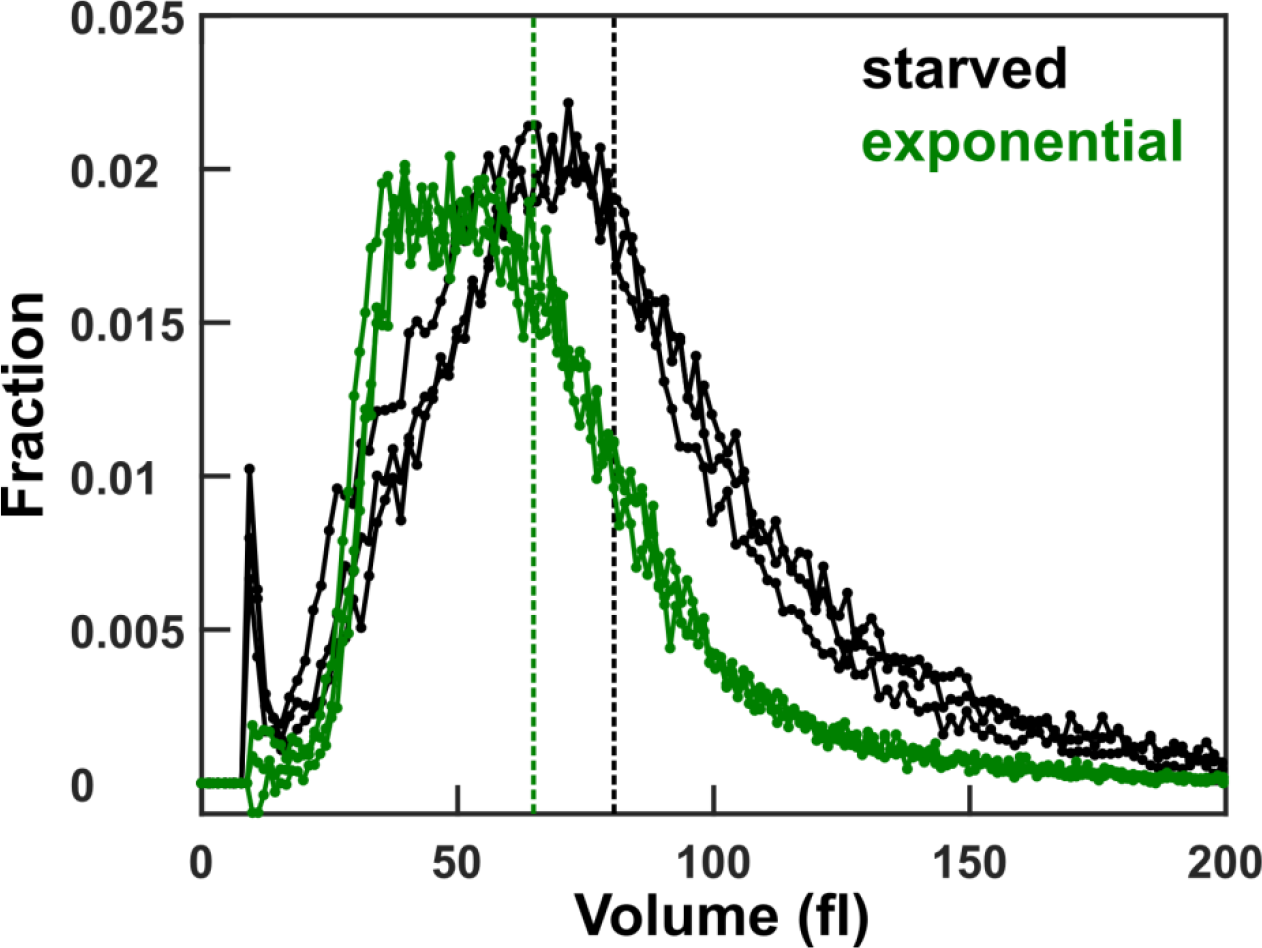
Slight cell size increase during *A*^*−*^*L*^*+*^ starvation. *A*^*−*^*L*^*+*^ cells (WY1340) grown in SD + excess hypoxanthine were either maintained at exponential phase (green) or washed and starved in un-supplemented SD for 24 hours (black). Cell sizes were measured using a Coulter counter. The initial peak in starved cells may represent dead cell debris. The average sizes of exponential and starved cells were 64.8 fl and 80.5 fl, respectively (dashed lines).

**Fig 6-Figure Supplement 4.**
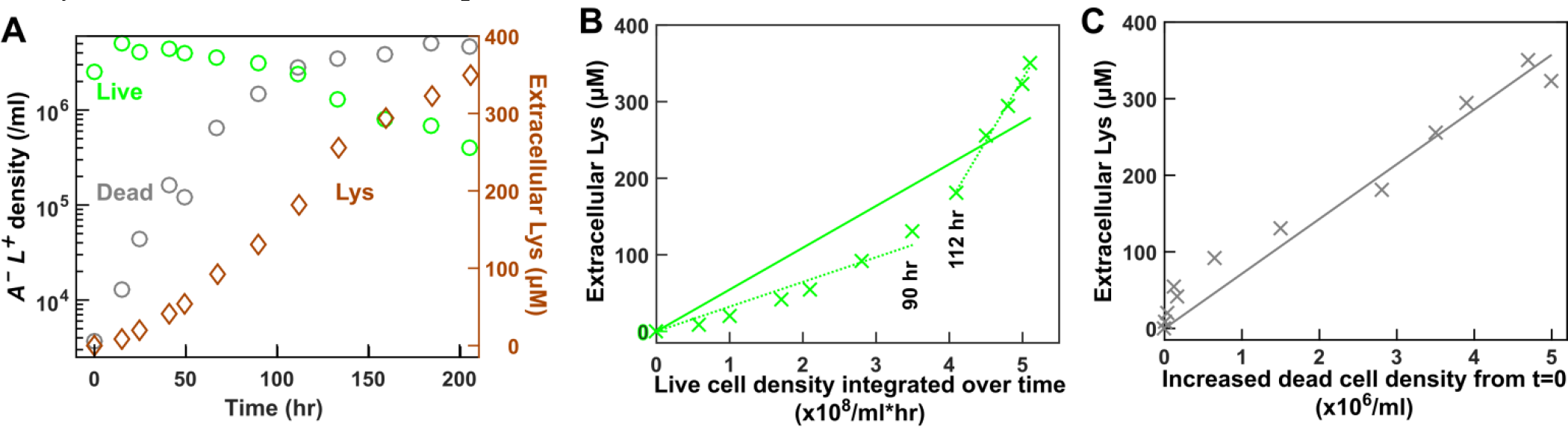
Lysine release by *A*^*−*^L^*+*^ during starvation is relatively constant during the initial 90 hrs. (**A**) Exponentially-growing *A*^*−*^*L*^*+*^ cells were washed and diluted into SD. Live and dead population densities were measured by flow cytometry, and lysine concentration was measured by the yield bioassay. Regression in both live release model (**B**) and dead release model (**C**) deviated from linearity. Since metabolite analysis suggests that live release is more likely (Fig 6-Figure Supplement 2B), we infer that release rate is time-variant - initially slow and then speeding up (a similar but less obvious trend can also be seen in Fig 6-Figure Supplement 1B). However, since CoSMO growth rate measurements rarely exceeded 96 hrs, we used the lysine release rate measured up to 90 hrs in Model ii.

**Fig 6-Figure Supplement 5.**
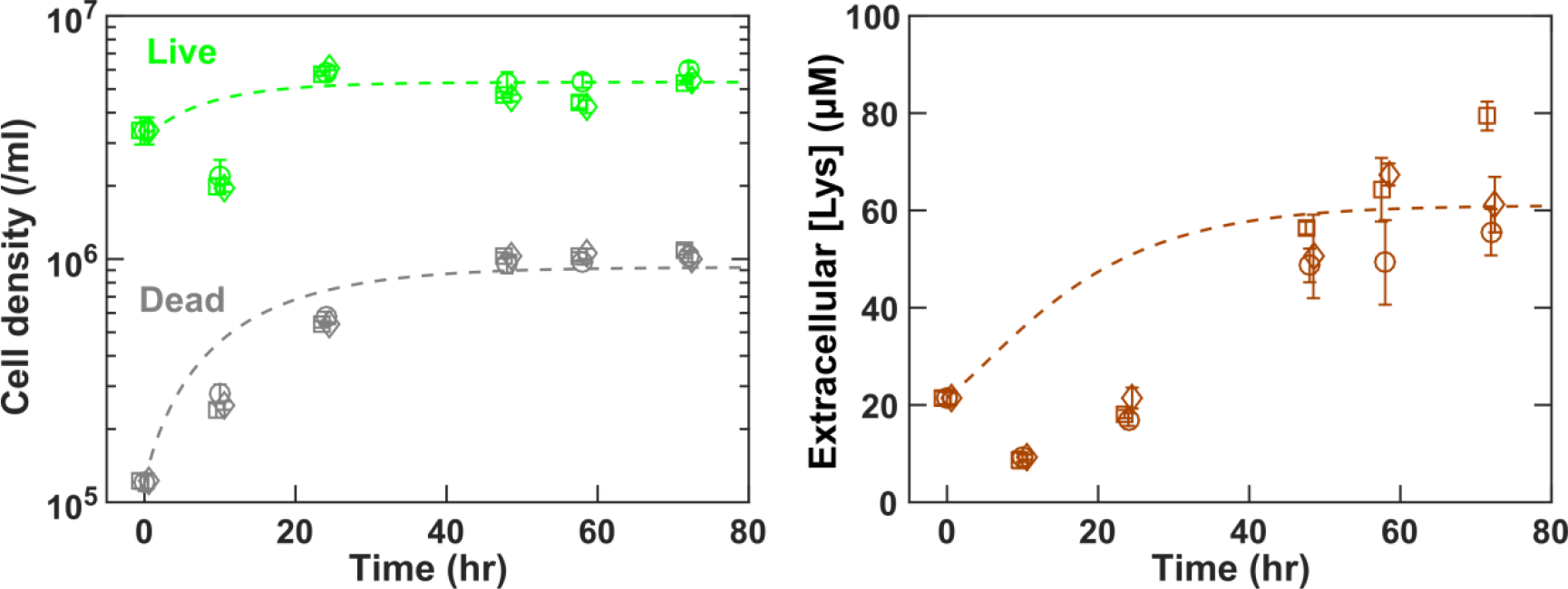
The *A*^*−*^*L*^*+*^ chemostat model can describe experimentally-observed chemostat steady state. *A*^*−*^*L*^*+*^ cells grown exponentially in SD + excess hypoxanthine were washed and diluted into SD, and pre-starved for 24 hrs. At time zero, starved cells (together with the medium which had already accumulated some lysine) were inoculated into chemostats, and fresh SD + 20 μM hypoxanthine was pumped in at a rate to achieve a doubling time of 8 hrs. Dynamics of live and dead populations (left) and of released lysine (right) were plotted (squares, circles, and diamonds representing three chemostats). Model (dashed lines; Fig 6-Code Supplement 1) was based on parameters in Table 1, except for a lysine release rate of 0.99 fmole/cell/hr which was averaged among the three chemostats. The initial decline in live cell density in experiments was presumably due to a growth lag when cells transitioned from starvation to chemostats, which was not modeled. The initial decline in extracellular lysine concentration in experiments is consistent with the live release model: reduced live cell density leads to reduced extracellular lysine.

**Fig 6-Figure Supplement 6.**
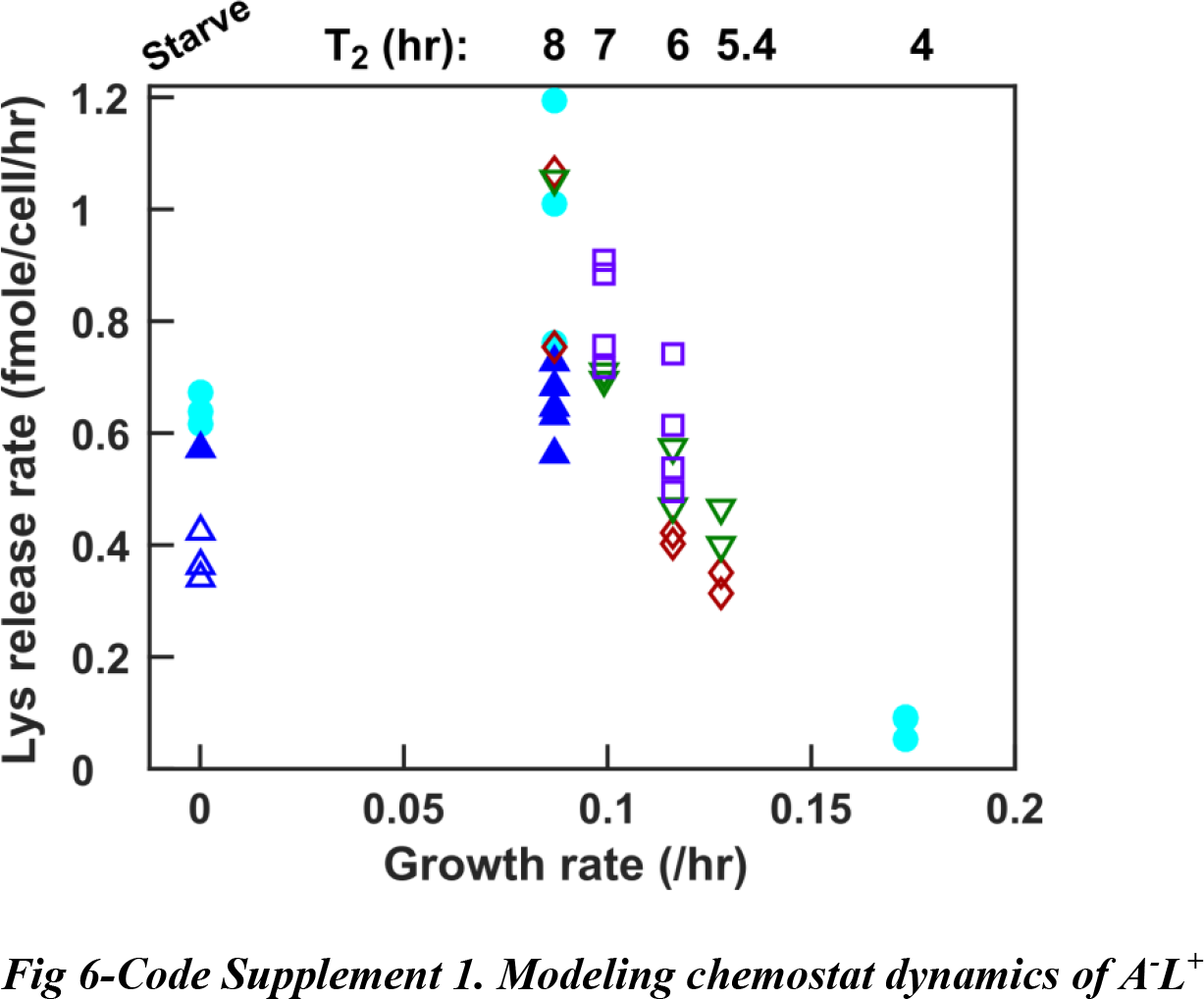
Measuring lysine release rate. *A*^*−*^*L*^*+*^ cells grown to exponential phase in SD plus excess hypoxanthine were washed and diluted into SD at time zero. Cells were either starved further (“Starve”) or inoculated into hypoxanthine-limited chemostats after 24 hrs of pre-starvation (e.g. Fig 6-Figure Supplement 1D-E; doubling times marked above). Each symbol represents an independent measurement, and measurements done at the same time were marked with the same color. Open and closed symbols represent pre-growth done in tubes versus flasks, respectively. We observed day-to-day variations in measurements (e.g. cyan circles higher than blue triangles). Lysine release rate data are in Table 1-Table Supplement 4.

***Fig 6-Code Supplement 1. Modeling chemostat dynamics of A^*−*^L^*+*^***

**Fig 7-Figure Supplement 1.**
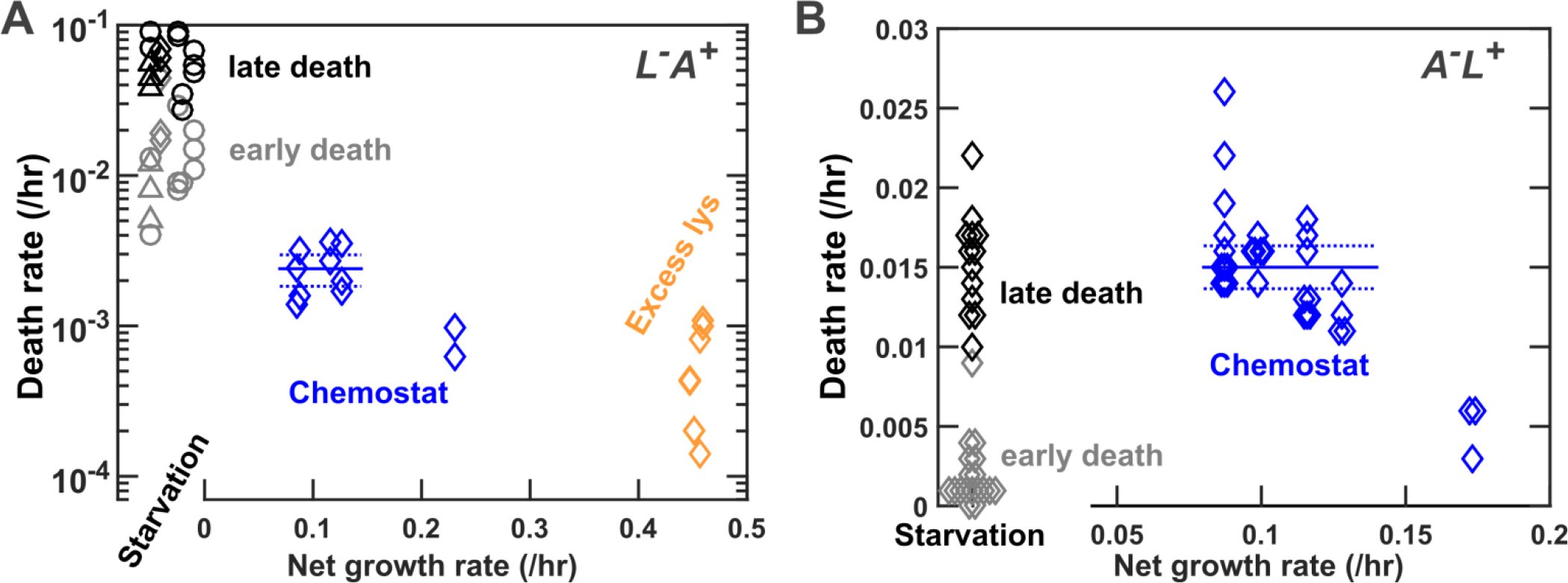
Death rates of *L*^*−*^*A*^*+*^ and *A*^*+*^*L*^*−*^ are relatively constant over a range of CoSMO-like environments. (**A**) Exponential *L*^*−*^*A*^*+*^ (WY1335) cells were washed in SD, and death rate was measured in chemostats at various doubling times (Methods, “Quantifying phenotypes in chemostats” Eq. 14; Fig 5-Figure Supplement 1 grey). As a comparison, death rates in batch cultures with zero or excess lysine are shown (see ^28^ for detailed methodology and data). With no lysine, the early-phase death rate (grey; from 5 to 12 hr postwash) was slower than the late-phase death rate (black; from 12 to 30 hr post-wash;). With excess lysine, death rates were very low (orange diamonds; Methods, “Calculating death rate in non-limited batch culture”). Death rates of *L*^*−*^*A*^*+*^ in chemostats (blue; doubling times from left to right being 8 hr, 6 hr, 5.5 hr, and 3 hr) were in-between death rates in starvation and in excess lysine. Blue solid and dashed lines mark the mean death rate ± 2 standard error of mean (SEM) from 5.5~ 8 hr doubling time chemostats (Table 1). Detailed data are in Table 1-Table Supplement 3. We used log plotting scale to visualize differences between small numbers. (**B**) Exponential *A*^*−*^*L*^*+*^ (WY1340) cells were washed and pre-starved for 24 hours. They were either further starved (black) or cultured at various growth rates in chemostats (blue). During starvation, death rate was initially slow and then sped up (see ^28^ for detailed methodology and data). Average death rate (blue solid line) and 2 SEM (blue dotted lines) were calculated from chemostats run at doubling times of 5.4 ~8 hrs (e.g. Fig 6-Figure Supplement 1F), and used in our model (Table 1). Detailed data are in Table 1-Table Supplement 4. We used linear plotting scale because early death rates were zero and could not be plotted on the log scale. In both **A** and **B**, death rates were quantified from the decline rate of ln(live population size), and live population size could be measured via microscopy total fluorescence intensity ^28^ (circles), microscopy live cell count ^28^ (triangles), or flow cytometry live cell density (diamonds, Methods “Flow cytometry”).

**Fig 7-Figure Supplement 2.**
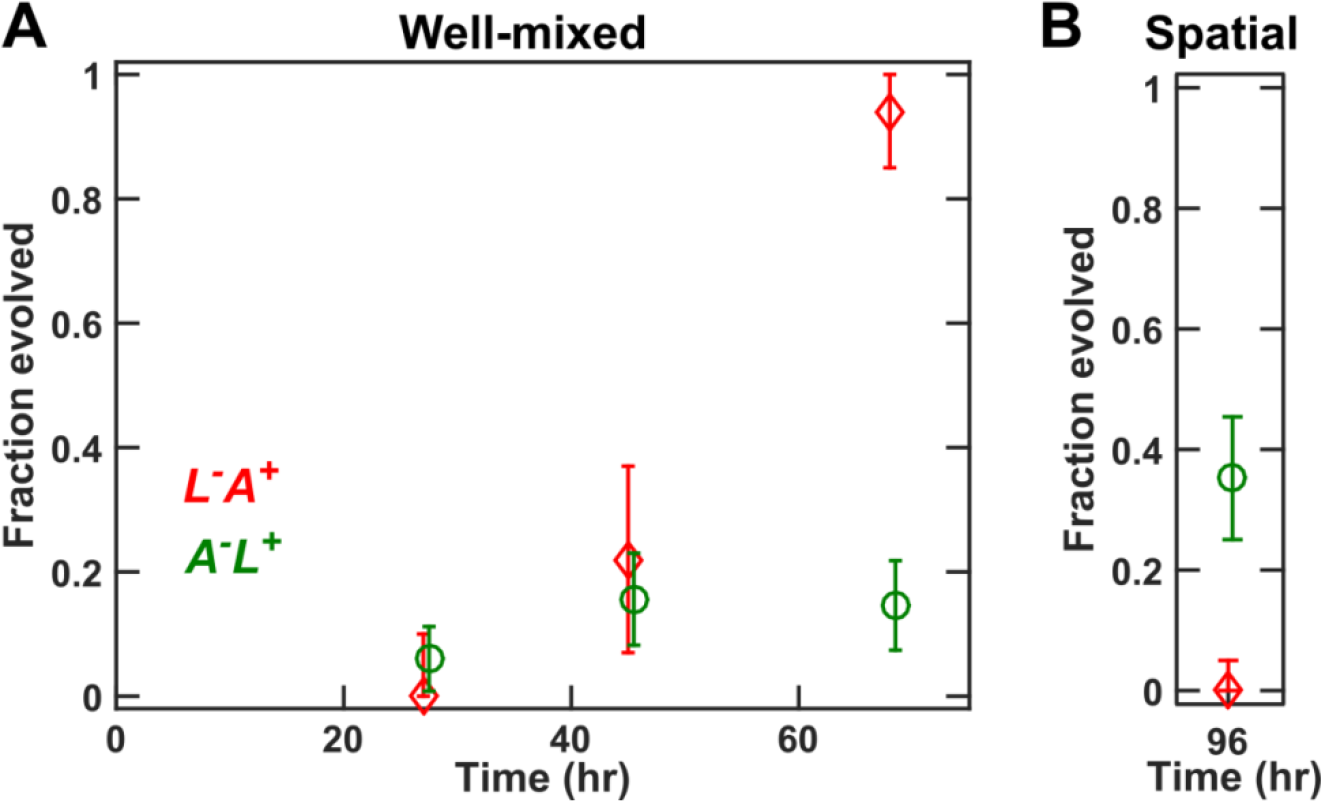
A spatially-structured environment slows down *L*^*−*^*A*^*+*^ evolution in CoSMO. (**A**) *L*^*−*^*A*^*+*^ evolves rapidly in a well-mixed environment. Exponentially-growing *L*^*−*^*A*^*+*^ (WY1335) and *A*^*−*^*L*^*+*^ (WY1340) were washed free of supplements, preconditioned, and mixed at 1:1 in SD at a total cell density of 10^5^/ml. The resultant CoSMO was grown in a well-mixed environment. At various times, samples were plated on YPD, and 32 *L*^*−*^*A*^*+*^ colonies (red diamonds) and 84~96 *A*^*−*^*L*^*+*^ colonies (green circles) were isolated to assay whether they were evolved or not (Methods, “Detecting evolved clones”). (**B**) *L*^*−*^*A*^*+*^ evolves slowly in a spatially-structured environment. *L*^*−*^*A*^*+*^ (WY1335 and WY1657) and *A*^*−*^*L*^*+*^ (WY1340 and WY1342) were mixed at approximately equal ratio and spotted onto the middle of an agarose slice containing 0.7 μM lysine (the “spotting” setting in Methods “Quantifying spatial CoSMO growth dynamics”; pre-starved *A*^*−*^*L*^*+*^ cells were washed again in SD so that CoSMO started with a defined level of lysine). At 96 hrs, CoSMO samples were plated on YPD, and 80 *L*^*−*^*A*^*+*^ colonies (red diamonds) and 88 *A*^*−*^*L*^*+*^ colonies (green circles) were isolated to assay whether they were evolved or not. For (**A**) and (**B**), error bars indicate two standard deviations according to binomial distribution. Specifically, if we observed *e* evolved clones among *N* total clones, then the fraction evolved was *p*=*e*/*N* and error bar was 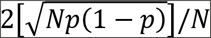. If no evolved clones were observed, then *p*=0 and the upper error bar was defined to be that of maximal *e* whose lower error bar spanned 0 (similar to Fig 3-Figure Supplement 4). Error bars were truncated at 0 and 1.

**Fig 7-Figure Supplement 3.**
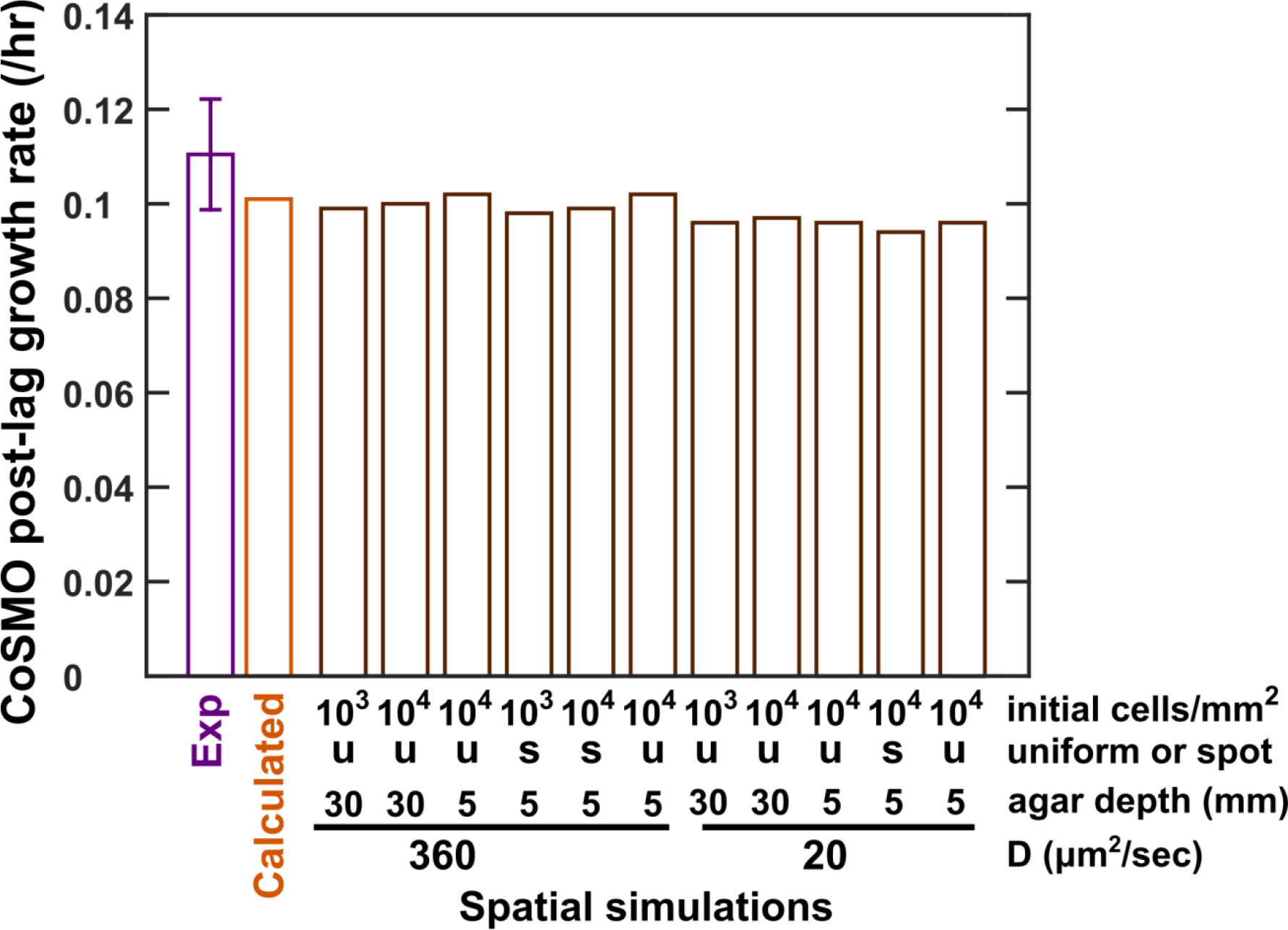
Predicting the steady state growth rate of spatial CoSMO. Spatial CoSMO growth were simulated under varying initial total cell density, inoculation setup (uniformly-plated “u” versus centrally-spotted “s”), agar depth, and diffusion coefficient (20 and 360 μm^2^/sec corresponding to diffusion coefficients in community and agarose, respectively ^24^). Spatial simulations yielded similar CoSMO growth rates (brown). Experimental measurements of spatial CoSMO (purple) and CoSMO growth rate calculated from Eq. 14 (orange) were taken from Fig 7 and plotted here for comparison. The spatial model and the calculation both considered variable lysine release rate.

**Fig 7-Figure Supplement 4.**
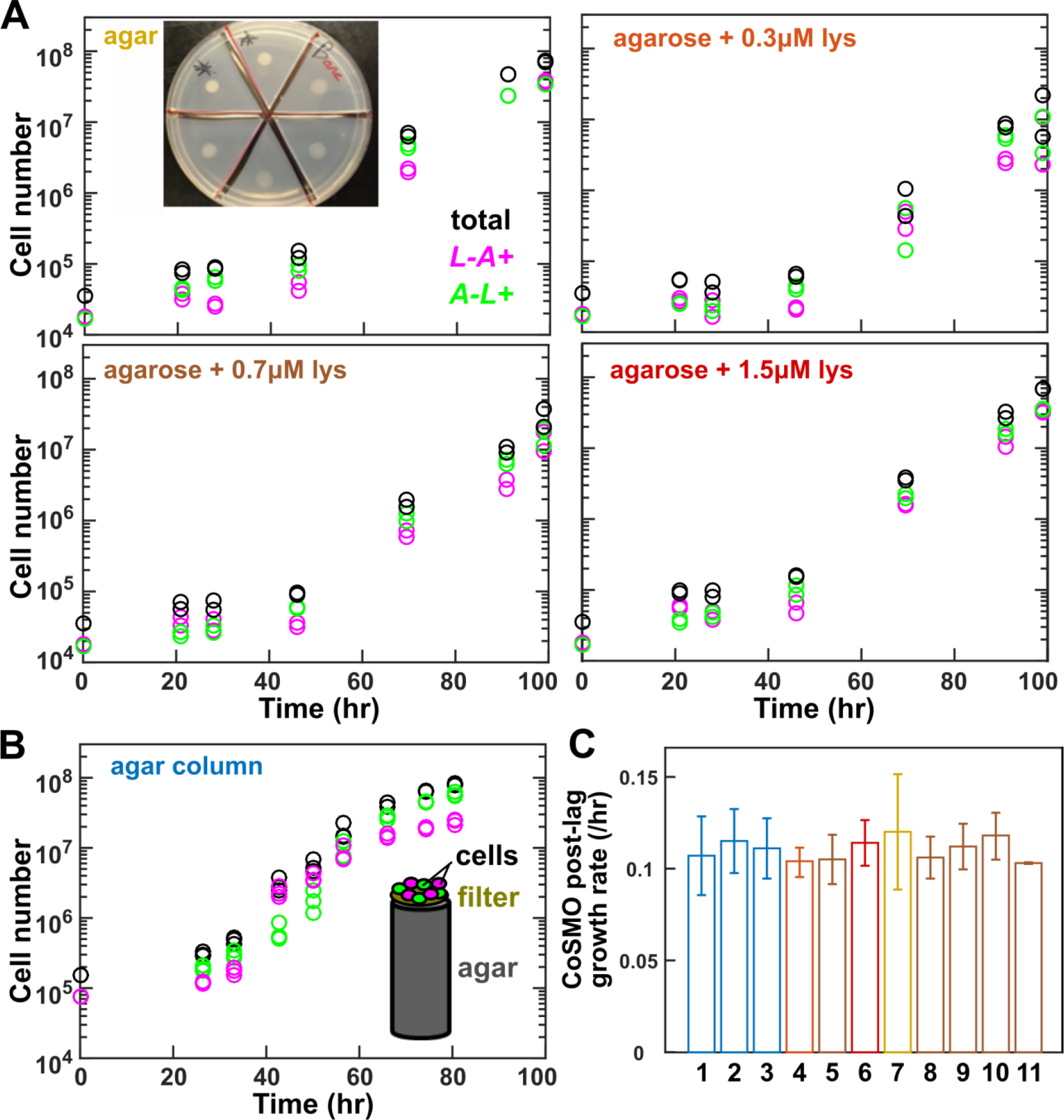
Population dynamics of spatial CoSMO. Preconditioned *L*^*−*^*A*^*+*^ and *A*^*−*^*L*^*+*^ were mixed at approximately 1:1 ratio and grown on 2xSD agar (which may contain trace nutrient contaminants) or agarose (Methods, “Quantifying spatial CoSMO growth dynamics”). (**A**) Growth dynamics of CoSMO on four media. Inset: shared experimental setup. 15 μl of 4×10^4^ total cells was spotted on the center of the cut pad, forming an inoculum spot of radius ~4 mm. (**B**) Growth dynamics of CoSMO in deep 96-well plates. 1.5×10^5^ initial total cells were filtered on top of a membrane filter to ensure uniform spatial distribution. This was equivalent to 3000 cells/mm^2^. (**C**) After the lag phase, steady state growth rates of CoSMO were calculated from 11 independent experiments, with color-coding corresponding to those in (**A**) and (**B**). Time points where total cell numbers exceed 1×10^8^ were excluded to avoid stationary phase. Error bars mark 2 standard error of estimating growth rate. In **A** and **B**, each data point represented the average of three flow cytometry measurements of a single spatial sample. Experimental data for **A** and **B** and summary data for C are provided in Fig 7-Table Supplement 1.

**Fig 7-Figure Supplement 5.**
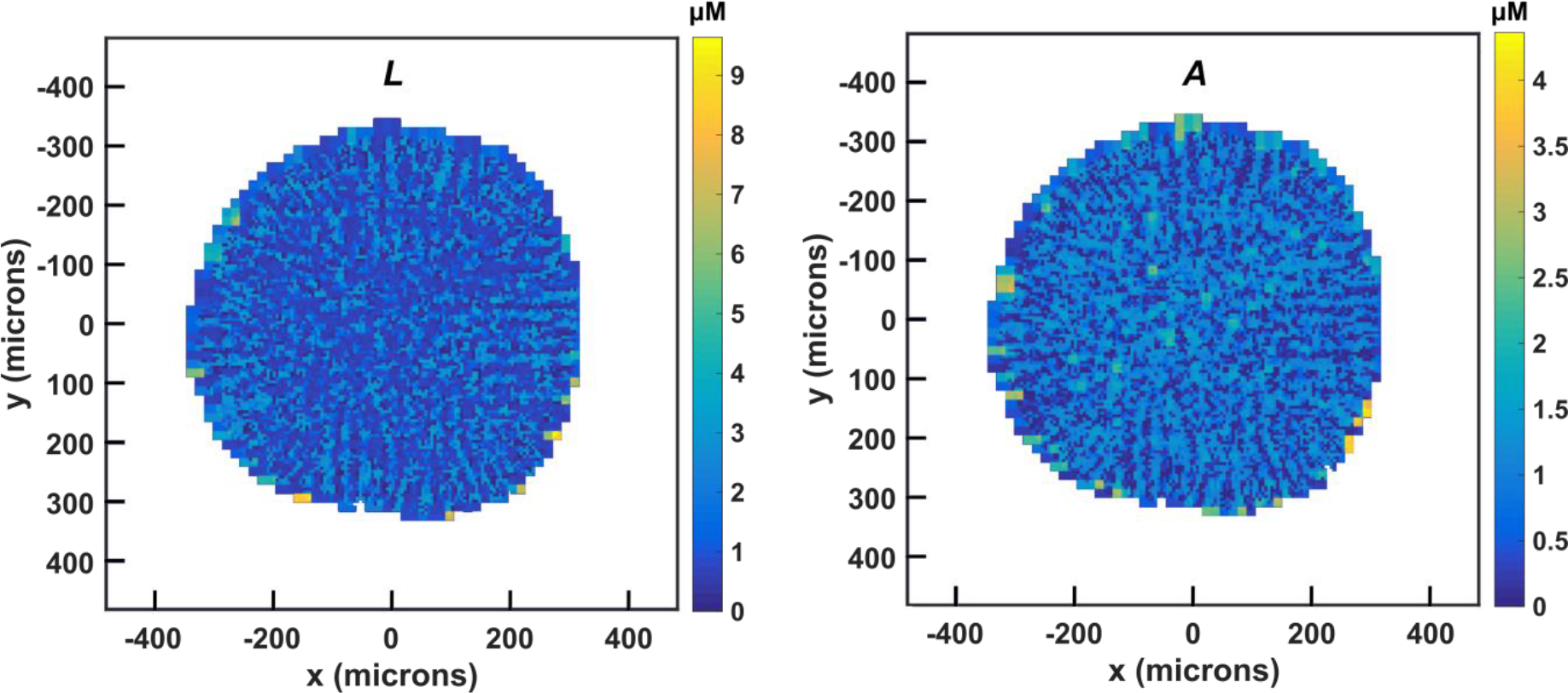
Spatial CoSMO eventually resembles well-mixed CoSMO. Metabolite concentrations in the agarose and the community eventually reach a nearly uniform state. Plotted are top views of lysine (*L*) and hypoxanthine (*A*) concentrations in a CoSMO community at the agarose surface 120 hrs after being spotted in the middle of an agarose pad. Since the populations were fairly intermixed within the community ^24^, the overall metabolite distributions remained fairly uniform within the community. The spatial averages of *L* and *A* in the community were 1.35 μM and 0.79 μM, respectively. The average concentrations in the agarose (1.31 μM for *L* and 0.73 μM for *A*) closely matched those inside the community. Thus, CoSMO growth rate in a spatially-structured environment is similar to that in a well-mixed environment. Here, the diffusion coefficients inside CoSMO and agarose were 20 and 360 μm^2^/sec, respectively.

***Fig 7-Code Supplement 1. Spatial CoSMO model with a similar diffusion coefficient for agar and community regions.***

***Fig 7-Code Supplement 2. Spatial CoSMO model with separate diffusion coefficients for agar and community regions.***

***Fig 7-Table Supplement 1. Experimental measurements of spatial CoSMO population dynamics and a summary of steady state CoSMO growth rate.***

## Acknowledgement

We thank Nick Buchler (Duke University) for TagBFP-AS-N constructs, Matt Rich (UW Seattle) for sequencing WY2447, Sarah Holte (Fred Hutch) for discussions on statistics, and Julia Laskin lab (Pacific Northwest National Laboratory) for providing preliminary nanoDESI MS data of chemical gradients between spatially-separated *L*^*−*^*A*^*+*^ and *A*^*−*^*L*^*+*^. Arvind “Rasi” Subramaniam, David Skelding, Maxine Linial, and Delia Pinto (Fred Hutch) critically read this manuscript.

